# A compact motif vocabulary is differentially interpreted across human cell types

**DOI:** 10.64898/2026.11.01.763377

**Authors:** Sam Shen, Wei Wang

## Abstract

How a common genome generates diverse human cell types may depend less on distinct cis-regulatory vocabularies than on context-dependent use of a shared one. We have developed CNN-CR, an interpretable neural network that learns regulatory motifs while predicting gene expression across 116 human biosamples. Promoter regulation proves highly compressible: dozens of recurrent motif families retain most of the predictive information of much larger cell-type-specific vocabularies and generalize to cellular contexts excluded from their discovery. These motifs are conserved at their binding sites and enriched at fine-mapped regulatory variants. Cellular context acts at two levels: chromatin state gates which motif instances are available for regulatory use, whereas reader identity determines how accessible motifs are interpreted. ChIP-seq data reveal frequent shifts in reader occupancy across cell types, and genome-scale perturbations support corresponding context-dependent reader functions. Thus, regulatory diversity can emerge through differential access to and interpretation of a compact shared DNA vocabulary.

## Introduction

How can the same genome generate hundreds of distinct human cell types? One possibility is that each cell type uses a largely distinct repertoire of cis-regulatory sequences. An alternative is that cells draw from a compact, shared vocabulary of regulatory motifs whose use and interpretation change with cellular context. Distinguishing between these models is central to understanding how regulatory information is encoded in DNA, how noncoding variants alter gene expression, and how synthetic regulatory programs can be designed.

Several observations make a compact regulatory vocabulary plausible. Although the human genome encodes approximately 1,600 transcription factors (TFs), many members of the same structural family recognize highly similar or nearly indistinguishable DNA motifs (*1–3*). A single motif can therefore be recognized by multiple candidate TFs, whereas the factor that actually occupies a particular motif instance may depend on chromatin accessibility, TF abundance, cooperative partners, and signaling state (*3–6*). Core regulatory circuit studies further suggest that relatively small sets of master TFs can dominate cell-identity programs during development (*7, 8*). Together, these observations raise the possibility that the diversity of human transcriptional states is generated not primarily by a large, diverse, cell-type-specific collection of DNA motifs, but by context-dependent use of a much smaller, shared cis-regulatory vocabulary.

Under this model, the motif vocabulary provides a common regulatory substrate, while cellular context acts at two distinct levels. First, chromatin state gates motif use: only motif instances located in accessible or otherwise active regulatory regions are available to participate in regulation. Second, reader identity determines motif interpretation: among the TFs capable of recognizing an available motif, the cellular environment determines which factor occupies it and therefore what regulatory consequence follows. The same DNA motif can thus be unavailable in one cell type, available in another, and interpreted by different TFs in different cellular contexts. Cell-type specificity can arise without changing the underlying vocabulary, by changing which copies of that vocabulary are usable and which factors read them.

This model makes two predictions. If chromatin provides the first contextual gate, the same motif patterns should carry much greater regulatory information when they occur within active regulatory regions than when their genomic occurrences are considered irrespective of chromatin state. If reader identity provides a second layer of interpretation, the same available motif family should be occupied by different compatible TFs in different cellular contexts, with corresponding differences in regulatory function. Whether a compact motif vocabulary can account for transcriptional variation across diverse human cell types under this two-layer model, and whether both mechanisms can be observed systematically, has not been established.

Testing this hypothesis requires learning regulatory sequence features without first prescribing the vocabulary to be found, and then comparing those features across many cellular contexts. Existing deep-learning models can predict gene expression from DNA sequence with high accuracy (*9–16*), but most do not yield compact, directly comparable motif vocabularies for individual cell types. Post hoc attribution methods (*17–20*) can identify sequence features associated with predictions, but those features are inferred after prediction rather than contributing through an explicit and interpretable model, so their contribution to a given prediction cannot be read off directly.

Here, we developed CNN-CR (Convolutional Neural Network with Contextual Regression (*21, 22*)), an interpretable architecture that couples de novo motif discovery to expression prediction within a single end-to-end model. Applying CNN-CR independently to each of 116 human biosamples from the EpiMap compendium (*23*) yields a per-biosample set of regulatory motifs with associated weights. We first tested the chromatin-gating layer and found that motif occurrences carry little additive predictive information when counted irrespective of chromatin state, but become strongly predictive when restricted to regulatory regions active in the corresponding biosample. That gated signal is, however, largely additive: within active regions, motif counts predict expression little better than the number of active regions alone, and only CNN-CR, which can combine occurrences non-additively, improves substantially on that baseline — indicating that the contribution of motif identity and combination is largely non-linear. Cross-sample comparison then revealed that promoter regulation is highly compressible: a recurrent set of 43 motif families, a number set by a recurrence cutoff whose consequences we test explicitly, retains most of the expression-predictive power in every context, approaching the accuracy of far larger biosample-specific motif sets and exceeding matched control motifs read through the same chromatin, so what the vocabulary contributes within the gate is motif identity rather than the chromatin it is read in. These families are conserved at their binding sites, enriched at fine-mapped regulatory variants, and predictive in biosamples excluded from their discovery. None of these properties — information content, evolutionary constraint, overlap with fine-mapped causal variants, occupancy in ChIP-seq, or reporter activity outside native chromatin — is used to define the families, and matched sets of 43 random JASPAR motifs fail each of them, so the compact vocabulary reflects biology rather than a computational artifact. Finally, because each motif family matches numerous candidate TFs, we test the reader layer directly: ChIP-seq shows that the factor occupying the same motif family frequently changes across cell types, and genome-scale perturbation supports corresponding context-dependent reader functions. Together, these results support a model in which a compact shared DNA vocabulary generates diverse regulatory outputs through two context-dependent operations: chromatin determines which motif instances are usable, and reader identity determines how those usable motifs are interpreted. Results

## Results

### CNN-CR predicts gene expression from epigenetically selected regulatory sequence with interpretable, cell-type-specific motif weights

We developed CNN-CR to jointly predict gene expression and extract the regulatory sequence features contributing to each prediction (Fig. 1A). For each gene, the model analyzes up to twenty 400 bp sequences — up to 10 located in promoters and 10 in distal enhancers, defined from sample-specific EpiMap (*23*) annotations based on epigenetic signals and predicted promoter-enhancer contacts from EpiTensor (*24*) (Methods). CNN-CR therefore learns from DNA sequence within epigenetically active regulatory regions. Convolutional filters discover de novo motifs, and Contextual Regression (*21, 22*) represents each prediction as a sum of motif occurrences weighted by gene-specific coefficients, providing an explicit motif-level decomposition.

**Fig. 1.**
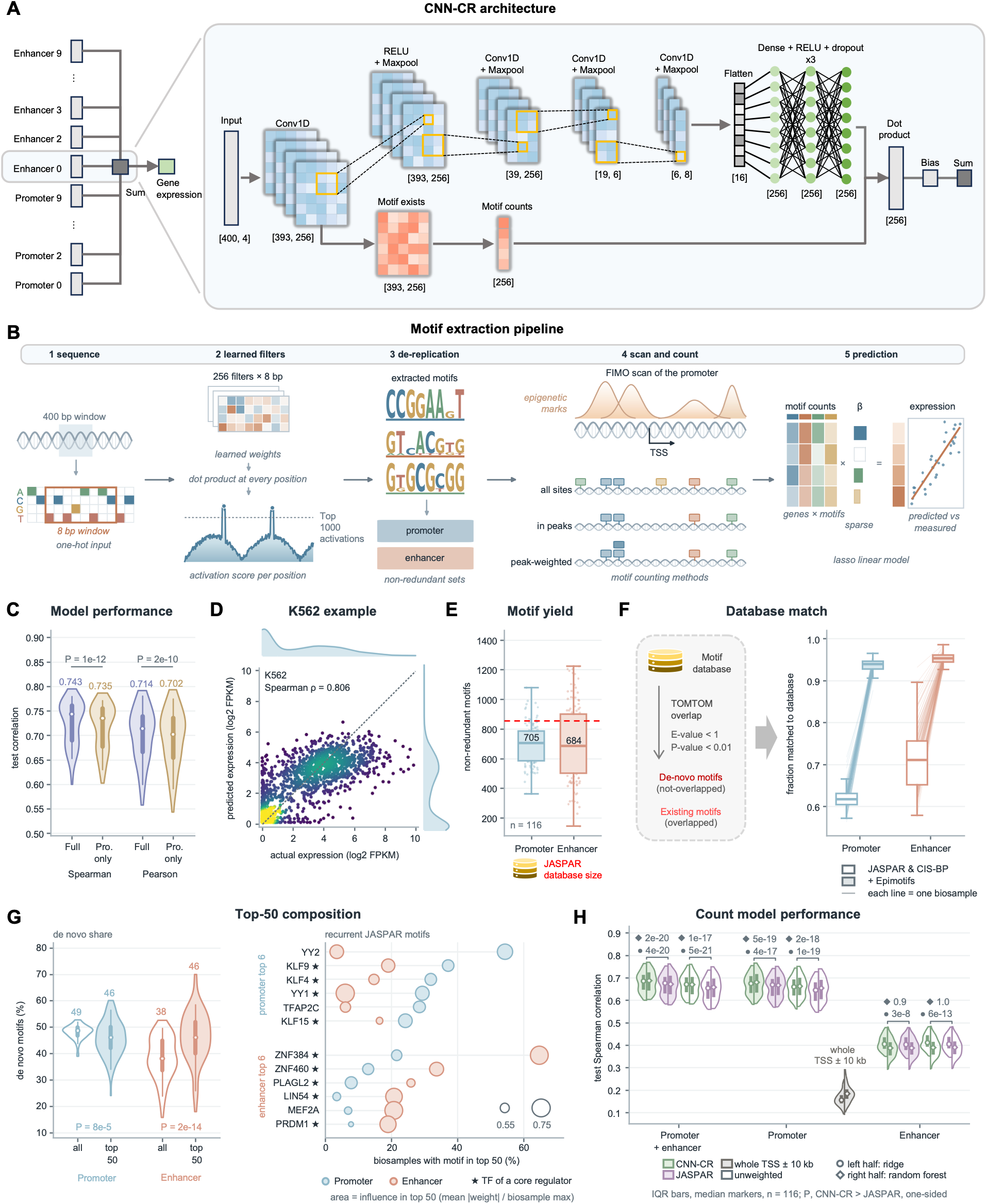
CNN-CR predicts gene expression from regulatory sequence and yields an interpretable motif vocabulary. **(A)** CNN-CR architecture: up to twenty 400-bp sequences per gene (ten promoters, ten enhancers) are processed through twenty parallel convolutional branches that do not share weights and meet only at the output, each combining a motif-finding module (256 filters of width 8 bp) with a contextual-weight module; each motif’s window-averaged activation is multiplied by its contextual coefficient and the products are summed (dot product) to predict expression. A full model therefore holds 20 × 256 = 5,120 first-layer motif filters and a promoter-only model 10 × 256 = 2,560. **(B)** Motif-extraction and de-replication pipeline: one-hot promoter windows -> convolutional filter activations -> top-activated k-mers - > per-model grouping and cross-model merge into non-redundant promoter/enhancer motif sets - > FIMO (*29*) scan under three counting rules -> count models (ridge regression and random forest). **(C)** Held-out test performance of the full (promoter + enhancer) versus promoter-only model, for Spearman and Pearson correlation, across the 116 biosamples for which both models were trained (median Spearman rho = 0.743 vs 0.734; median Pearson = 0.714 vs 0.702; paired one-sided Wilcoxon P = 1.3 × 10⁻¹² and 1.6 × 10⁻¹⁰). **(D)** Predicted versus measured expression in a representative cell line, K562 (log2 FPKM, the training target; Spearman rho = 0.806; dashed line, identity; colour, local point density). **(E)** Number of unique non-redundant motifs extracted per biosample for promoters and enhancers (median 705 and 684; n = 116 biosamples, each shown as a point). **(F)** Concordance of de novo motifs with known databases (JASPAR/CIS-BP, and all three including Epimotifs): promoter 62%->94%, enhancer 71%- >95%; each faint line joins one biosample across the two database sets. **(G)** Composition of the 50 highest-weighted motifs per biosample. Left, percentage of motifs with no JASPAR match, among all motifs versus the top 50 ranked by contextual weight, for promoters and enhancers (promoter 49%->46%, enhancer 38%->46%; paired Wilcoxon P as shown, n = 116). Right, the six JASPAR motifs recurring most often in the top-50 lists of each class; x axis, percentage of biosamples in which the motif ranks top-50; bubble area, mean |linear weight| of that motif relative to the strongest motif of the same biosample (its influence where it ranks); star, TF assigned to one of the core regulators of Fig. 2. **(H)** Chromatin gating of motif information. Count-model performance for CNN-CR versus JASPAR motifs under peak-weighted and unweighted (hatched) counting, for promoter+enhancer, promoter-only and enhancer-only feature sets. Each violin is one condition over the 116 biosamples: left half, ridge regression (circle, median); right half, random forest with 200 trees (diamond, median); bars, interquartile range (median Spearman ρ, ridge / random forest, CNN-CR then JASPAR motifs: promoter + enhancer, peak-weighted 0.686 / 0.687 and 0.676 / 0.671, unweighted 0.670 / 0.670 and 0.653 / 0.658; promoter only, peak-weighted 0.675 / 0.679 and 0.669 / 0.667, unweighted 0.659 /0.662 and 0.644 / 0.654; enhancer only, peak-weighted 0.405 / 0.387 and 0.403 / 0.387, unweighted 0.411 / 0.389 and 0.405 / 0.390). Grey, the same promoter motifs counted at every occurrence across the entire TSS ± 10 kb of each gene, without any chromatin restriction (median ρ = 0.156 linear, 0.184 random forest; below the peak-restricted model in all 116 biosamples under both models). Brackets, paired one-sided Wilcoxon for CNN-CR > JASPAR within each pair, lower value for ridge (●), upper for the random forest (◆).

The model’s input representation consequently contains information from both which regulatory regions are active and what sequence features occur within those regions. The chromatin landscape layer is substantial on its own. Used as the single feature of a linear model fitted on the same 90/10 gene split as the CNN-CR model, the number of active promoter windows a gene receives predicts its expression at a median Spearman ρ = 0.641 in held-out genes across the 116 biosamples (fig. S1): genes with more accessible, histone-marked promoter regions tend to be more highly expressed. As a comparison, CNN-CR reaches ρ = 0.743 on the same biosamples in held-out genes (Fig. 1C,D), consistent with additional predictive information being carried by sequence features within those regions, such as which motifs are present, in what combination, and in which windows.

Each CNN-CR model in a biosample is an ensemble of ten independently initialized networks trained with internal validation folds while preserving the same 10% of genes as an untouched test set (Methods, fig. S2). Accuracy was highest in well-characterized cell lines, with K562 and GM12878 the two best-performing biosamples (ρ = 0.806 and 0.803); prediction quality remained comparable across established cancer lines, primary immune cells, embryonic stem cells and a differentiated neuron sample (fig. S3). A promoter-only model reached median ρ = 0.734, showing that promoter sequence captured most of the predictable signal while the assigned distal enhancers provided a smaller aggregate improvement.

CNN-CR approaches the predictive accuracy of substantially larger sequence models at a small fraction of their cost. Across the 59 biosamples for which Enformer predictions were available, CNN-CR reached a median held-out ρ = 0.641 against Enformer’s 0.680 (paired Wilcoxon P = 4.6 × 10^-8^; fig. S4). CNN-CR therefore approaches the accuracy of a model with a 100 kb receptive field and approximately 32 times more parameters, while training in under 30 minutes per biosample on a single GPU and predicting at negligible cost (fig. S4E).

To confirm that CNN-CR’s filters learn genuine expression-predictive sequence patterns, we trained it on simulated data in which defined motifs were embedded into dinucleotide-shuffled promoter sequences and expression was generated from their motif counts plus noise (Methods). The models reached median ρ = 0.943 for the simulated expression and recovered a median of 93.3% of the embedded motifs per biosample from the first convolutional layer, counting a match at p ≤ 1.2 × 10^-3^, which normalizes the threshold to the JASPAR comparisons below (Methods; fig. S5). This positive control shows that, when expression is generated from known motif content, CNN-CR predicts it almost completely and recovers almost all of the underlying sequence features from which the prediction is made.

Beyond predicting expression, each trained model yields an explicit motif vocabulary. From its convolutional filters we extracted a non-redundant set of roughly 700 de novo motifs per biosample (median 705 for promoters and 684 for enhancers), ranked by contextual weight, the contribution associated with each motif in the fitted model (Fig. 1B,E; Methods). This importance distribution is strongly concentrated. Each model learns 2,560 promoter filters, of which the 750 highest-weighted account for 89% of the summed contextual weight. Motif information content falls steadily with contextual-weight rank, and the retained filters are more information dense than the discarded ones (11.53 versus 10.20 bits for promoters; fig. S6).

Most motifs extracted are recognizable TF binding patterns. A median of 62% of promoter motifs and 71% of enhancer motifs match JASPAR (*25*) or CIS-BP2 entries, rising to 94% and 95% respectively when histone modification or DNA-methylation associated motifs (i.e. EpiMotifs) (*26–28*) are included (Fig. 1F). The unmatched are candidate novel binding patterns. These uncatalogued motifs are not confined to low-weight features: among the 50 highest-weighted motifs of each biosample, a median 46% have no JASPAR match in both promoters and enhancers (Fig. 1G). For enhancer motifs this is a marked rise over their own background rate of 38% (paired Wilcoxon P = 1.7 × 10^-14^, n = 116 biosamples), while promoter motifs are already at 49% and change little (46%; P = 8.4 × 10^-5^). The recognizable TF motifs differ sharply by class: promoter top-50 lists recur on YY2 (in 54% of biosamples), KLF9 (37%), KLF4 (32%), YY1 (29%), TFAP2C (28%) and KLF15 (24%), whereas enhancer lists recur on ZNF384 (65%), ZNF460 (34%), PLAGL2 (26%), LIN54 (21%), MEF2A (21%) and PRDM1 (19%) (Fig. 1G). Nine of these twelve factors are among the TFs assigned to the core regulators defined below, and motifs of the ELF, ELK, KLF, ZNF, and YY families recur broadly across cell-type categories (fig. S7).

### Chromatin state gates the predictive information carried by regulatory motifs

We next asked whether the predictive information carried by motif occurrences depends on chromatin state. For each biosample we counted the occurrences of that biosample’s own de novo motifs per gene under different chromatin constraints and fitted two count-based models to the same genes and the same held-out split, a regularized linear model and a random forest (Methods). Holding the motif set, the genes and the model fixed isolates the effect of where the motifs are counted. The dependence on chromatin is pronounced (Fig. 1H). Counting every occurrence of the same promoter motifs across the entire TSS ± 10 kb of each gene, without reference to chromatin state, gave a median held-out ρ of 0.156 for the linear model and 0.184 for the random forest across the 116 biosamples (interquartile ranges 0.142–0.178 and 0.162– 0.204; about 2,100 occurrences per gene); counting only the occurrences inside the biosample’s active promoter peaks raised it to ρ = 0.659 and 0.662, and weighting each occurrence by the number of overlapping peaks raised it to ρ = 0.675 and 0.679 (0.686 and 0.687 with enhancer peaks added). The peak-restricted model outperformed the unrestricted model in every one of the 116 biosamples under both model classes.

These results establish the first contextual layer of the model proposed above: chromatin state gates which genomic copies of the motif vocabulary are available for regulatory use. The DNA sequence patterns themselves are present throughout the promoter region, but they carry little additive information when counted without regard to chromatin and become strongly predictive when restricted to regulatory regions active in the corresponding biosample.

Within this chromatin-gated framework, the de novo motifs discovered by CNN-CR modestly but consistently outperformed the full JASPAR motifs for promoter and promoter-plus-enhancer features under both model classes (promoter + enhancer, peak-weighted: ρ = 0.686 versus 0.676 for the linear model and 0.687 versus 0.671 for the random forest; paired one-sided Wilcoxon P ≤ 2.7 × 10^-8^ for all six linear comparisons and P ≤ 2.0 × 10^-17^ for the four random-forest comparisons on promoter-containing features, n = 116), whereas for enhancer-only features the two motif sets were indistinguishable under the random forest (P = 0.9 and 1.0). Promoter motifs alone performed nearly as well as the promoter-plus-enhancer motifs under every motif set, counting rule and model class (median ρ 0.675 versus 0.686 for the linear model and 0.679 versus 0.687 for the random forest on peak-weighted CNN-CR motifs, and within 0.012 in the other variants), whereas enhancer motifs alone fell to ρ = 0.387–0.411. The random forest never exceeded the linear model by more than 0.010 on promoter-containing features, so the information in motif counts is captured almost entirely by an additive model once chromatin is accounted for. The signal captured by this framework is therefore predominantly promoter-driven, motivating our subsequent focus on promoter motifs when asking whether the large biosample-specific vocabularies can be compressed into a smaller shared set (Fig. 1C).

### A compact motif vocabulary is shared across human cell types

Having derived a de novo motif vocabulary for each of the 116 biosamples, we asked whether these vocabularies organize around a smaller, shared set of recurrent motifs. From here onward we focus on promoter motifs and promoter sequences because promoter-associated features captured most of the predictive signal under the CNN-CR framework (Fig. 1C). For the cross-biosample analysis, each biosample’s promoter filters were first reduced to representative PWMs within that biosample: filters were clustered by similarity, a cluster was carried forward only when at least one member matched a JASPAR motif (E ≤ 0.05, p ≤ 0.01) whose TF was expressed in that biosample (log2 FPKM > 1.66), clusters annotated with the same expressed TF were merged as redundant representations of one factor, and each merged cluster was represented by one PWM (median 73 per biosample, 8,441 in total; Methods). These representatives constitute a more stringent, expression-filtered subset of the larger motif vocabularies used in Fig. 1E, which are used unchanged in the count-based models. We pooled the 8,441 representative PWMs and asked which recur across biosamples. Every motif was compared with every other by TOMTOM, which aligns two PWMs and scores their column-wise Pearson correlation, and a pair was accepted as a match when its E-value was ≤ 0.5 with p ≤ 0.01 over an overlap of at least 7 bp (Supplementary Methods S3). The matches were assembled into a similarity graph in which each node is a motif and an edge, weighted by the match E-value, links two motifs learned independently from different biosamples. Motif families were then identified by a shortest-path clustering: a motif joins a family only if its path distance to every current member is within a specified bound (Methods), so each family is a set of mutually similar motifs drawn from many biosamples; every family therefore represents one recurring motif independently rediscovered across multiple cellular contexts rather than a motif defined by a single biosample (Fig. 2A; Methods).

**Fig. 2.**
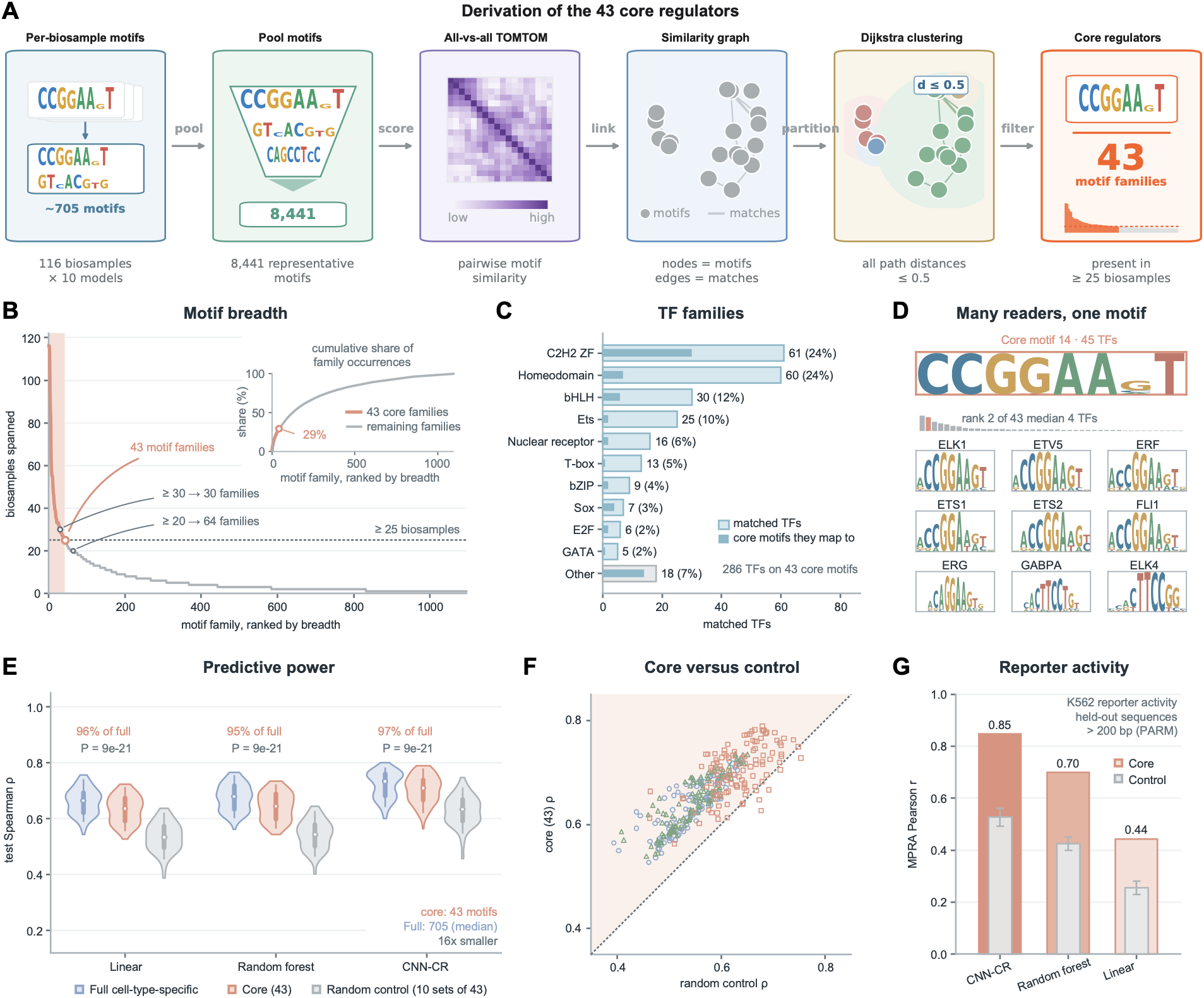
A compact set of 43 core regulators recurs across biosamples and retains near-full predictive power. **(A)** Derivation of the core motifs: the expression-filtered representative promoter PWMs of each biosample (median 73 per biosample, 8,441 pooled over 116 biosamples; Supplementary Methods S3) are compared all-versus-all by TOMTOM (*32*), assembled into a motif-similarity graph, clustered by greedy shortest-path (Dijkstra) clustering on the E-value-weighted graph, a motif joining a family only when its path distance to every member is <= 0.5 (Methods), and filtered to families recurring in at least 25 biosamples, yielding the 43 core regulators. **(B)** Cross-biosample coverage curve: motif families ranked by the number of biosamples they span; requiring recurrence in >= 25 of 116 biosamples (dashed line) leaves 43 motif families (orange, retained families); the cutoff falls where the curve has flattened but at no discontinuity, so it is a convention whose consequences are tested in fig. S8. Inset, cumulative share of all family occurrences (each family counted once per biosample it spans; 6,692 in total over the 1,095 families) captured by the families ranked up to each breadth; the 43 core families, 4% of the families, account for 29% of all occurrences. **(C)** The transcription factors matched by the 43 core motifs (JASPAR 2022, TOMTOM) distributed across their structural families, dominated by C2H2 zinc-finger and homeodomain factors. **(D)** One core motif matched by many candidate transcription factors. Top (orange border): the de novo core motif 14, an ETS binding pattern. Below: sequence logos (information content, bits) of nine biologically prominent ETS-family factors it matches by TOMTOM (JASPAR 2022) — ELK1, ETV5, ERF, ETS1, ETS2, FLI1, ERG, GABPA and ELK4 (GABPA and ELK4 reverse-complemented to the core orientation) — all recognizing a near-identical CCGGAA(G)T element, illustrating the many-to-one motif-to-factor structure. The full atlas pairing every core motif with all its matched JASPAR logos is in data S2. **(E)** Benchmark of three motif sets — the full cell-type-specific de novo vocabulary (median 705 motifs per biosample), the 43 core motifs, and random control motifs (ten independent sets of 43 JASPAR motifs resembling no core motif; Methods) — under three model classes (linear, random forest, CNN-CR) across 116 biosamples with matched gene splits. Each control value is the mean over the ten sets for that biosample. The 43 core motifs recover ∼96% of promoter only full-vocabulary performance (median test rho = 0.636/0.647/0.710 vs 0.675/0.679/0.734) with 10–20-fold fewer features, whereas random controls reach 0.544/0.551/0.640; core > control in all 116 biosamples in every class (paired Wilcoxon p = 9.0 × 10-21 for all 3 models). **(F)** Per-biosample core-motif-versus-control-motif held-out test Spearman rho for each model class; almost all 116 biosamples are above the diagonal (core motif > control motif). **(G)** Held-out prediction of measured reporter activity (MPRA) by three model classes trained on the core motifs; a motif-initialized convolutional network (CNN-CR) attains Pearson r = 0.848, versus 0.700 (random forest) and 0.443 (linear). Random 10 sets of controls attain mean r = 0.520 (CNN-CR), 0.411 (random forest) and 0.254 (linear).

The breadth of these families varied continuously. Family coverage declines smoothly with rank, without a discontinuity that would mark a unique boundary between broadly recurring and cell-type-restricted motifs (Fig. 2B); the family ranked 43rd spans 25 biosamples and the family ranked 44th spans 24. We therefore use recurrence in at least 25 of 116 biosamples as an operational definition of a compact core vocabulary, rather than as evidence for a biologically privileged number of motifs. This criterion yielded 43 core motif families (Fig. 2A,B, data S1). Although they are only 43 of 1,095 families, they account for 29% of all family occurrences across the 116 biosamples (Fig. 2B, inset), so the compact set is also the most frequently reused part of the vocabulary. Two analyses showed that the compact vocabulary was robust to the exact recurrence threshold. Re-deriving the families from 100 random subsamples of 93 biosamples produced sets of similar size (median 45 families); in a typical replicate 36 of the 43 families were called core again, and 40 of the 43 kept at least half of their membership (median best-match Jaccard index 0.79; fig. S8C,D; table S1). In a complementary analysis, recurrence thresholds spanning ≥10 to ≥ 45 biosamples generated vocabularies of 177 to 14 motifs, yet their held-out predictive performance remained within 5% of that obtained with the 43-motif set under the linear and random-forest models (fig. S8E,F; table S1; Methods), so the analyses below would be unchanged had a neighbouring threshold yielded 38 or 45 families instead. The principal result is therefore not the exact number 43, but that the much larger biosample-specific motif repertoires are strongly compressible into a small recurrent basis.

The 43 families also reveal a many-to-one relationship between motif sequence and TF identity. Each motif family matches several candidate TFs (a median of four, and as many as 51, per motif); across all 43 motifs these amount to 287 JASPAR entries, corresponding to 254 distinct TFs after removing heterodimers spanning ten structural families, most prominently C2H2 zinc fingers (including KLF, SP, and ZNF384) and homeodomain factors (Fig. 2C). Compared against the EpiMotifs, 12 of the 43 also resemble a histone-mark-associated motif and 2 a DNA-methylation-associated motif (E ≤ 0.05). This many-to-one structure is a defining feature of the core set: a single motif is typically matched by an entire family of related factors with near-indistinguishable binding preferences. The motif 14, for example, is matched by nine prominent ETS-family factors — ELK1, ETV5, ERF, ETS1, ETS2, FLI1, ERG, GABPA and ELK4 — whose JASPAR logos are nearly identical (Fig. 2D). Thus, the compact vocabulary specifies recurrent DNA-recognition patterns but does not, by sequence alone, specify which compatible TF reads them in a particular cell type.

An interesting observation is that every core family matches a catalogued motif in JASPAR but roughly half of each biosample’s most predictive motifs have no JASPAR match (Fig. 1G). We repeated the cross-biosample clustering specifically for the JASPAR-unmatched motifs (Methods) and obtained 2,623 families, but very few of them recur broadly: only 19 (0.72%) span ≥ 25 biosamples against 3.93% of the catalogued groups, a 5.4-fold difference (fig. S9A). In other words, JASPAR-unmatched motifs are much more cell-type specific than the JASPAR-matched motifs. Furthermore, most broadly recurring unmatched families are low-complexity patterns using at most two distinct bases (74%, against 21% of the core motifs and 8% of the uncatalogued motifs in K562’s top 50; Fisher exact P = 1.4 × 10^-4^), and carry below-median weight in the biosamples where they recur (contextual P = 1.1 × 10^-8^, linear P = 1.2 × 10^-4^; n = 115 biosamples; fig. S9C,D). Adding them to the 43 motifs did not increase predictive power, even with the 43+19=62-motif model tuned and the 43-motif model left at the configuration used throughout (mean Δρ = −0.002 across five biosamples spanning the performance range, improved in two of five; paired Wilcoxon P = 0.63; fig. S9E). Therefore, we focused on motifs that match a catalogued JASPAR motif with at least one candidate reader expressed in that biosample (Methods). A catalogue match names the factors able to read a motif, so their expression can be checked here and their binding tested directly later, whereas an unmatched motif names no factor to follow.

### A compact vocabulary retains most of the predictive power of full biosample-specific motifs

We evaluated how much of the predictive power carried by the full, biosample-specific motif vocabularies is retained by the compact set of 43 motif families. The full vocabularies are an order of magnitude larger at a median of 705 de novo motifs per biosample, with each biosample represented by its independently learned motif set, whereas the compact vocabulary contains the same 43 motif families in every context. This provides a direct test of whether regulatory sequence information distributed across hundreds of independently learned motifs can be approximated by a much smaller common basis.

Across all 116 biosamples, the 43 motifs retained most (94–97 %) of the predictive performance of the promoter only full vocabulary (Fig. 2E; Methods): median held-out test Spearman correlations were ρ = 0.636 versus 0.675 for the additive linear model, 0.647 versus 0.679 for a random forest, and 0.710 versus 0.734 for CNN-CR. To confirm that this is a property of the 43 core motifs rather than a trivial consequence of scanning promoters with any 43 position weight matrices, we compared them with ten independent sets of 43 random JASPAR motifs selected to have no detectable similarity to the core families. The 43 random JASPAR motifs performed worse in every model class (median ρ = 0.544, 0.551 and 0.640) across all 116 biosamples (paired Wilcoxon p = 9.0 × 10^-21^ for all 3 models; Fig. 2E,F).

We investigated whether the core motifs carry regulatory information outside the endogenous chromatin environment in which they were discovered. We therefore evaluated them using the K562 massively parallel reporter assay (MPRA) (*30*), in which promoter fragments drive reporter expression outside their native genomic context (the PARM library (*31*); Fig. 2G, Methods). Scanning the 43 recurrent motifs across the reporter sequences, models predicted held-out promoters’ reporter activity, with Pearson correlations of r = 0.443 for the linear model, 0.700 for random forest and 0.848 for a motif-initialized CNN-CR (i.e. the CNN’s first layer filters frozen to the 43 PWMs, see Methods), compared with 0.254, 0.411 and 0.520, respectively, for 10 sets of control motifs (Fig. 2G), which suggests that the 43 motifs are not a statistical artifact and motif identity matters for prediction accuracy. The CNN-CR result approached the reported Pearson r = 0.920 for the full PARM model (K562, sequences > 200 bp; Fig. 2G). Because MPRA removes the native chromatin environment, these results provide complementary evidence that the compact motifs encode transferable, sequence-intrinsic regulatory information rather than dataset-specific correlations. In the genome, chromatin determines which instances of that vocabulary are available for use; in the reporter assay, where the assayed sequence is experimentally made available, the same vocabulary retains substantial predictive power. This distinction is consistent with the chromatin-gating result above while showing that the sequence patterns themselves carry regulatory information rather than merely serving as markers of active chromatin.

### Core motifs show sequence specificity and evolutionary constraint at their genomic sites and among candidate readers

We examined whether the 43 core motifs exhibit independent properties consistent with biological importance by checking their information content, evolutionary conservation of their predicted genomic occurrences, and human mutational constraint of candidate TFs whose binding preferences match the motifs (Fig. 3).

**Fig. 3.**
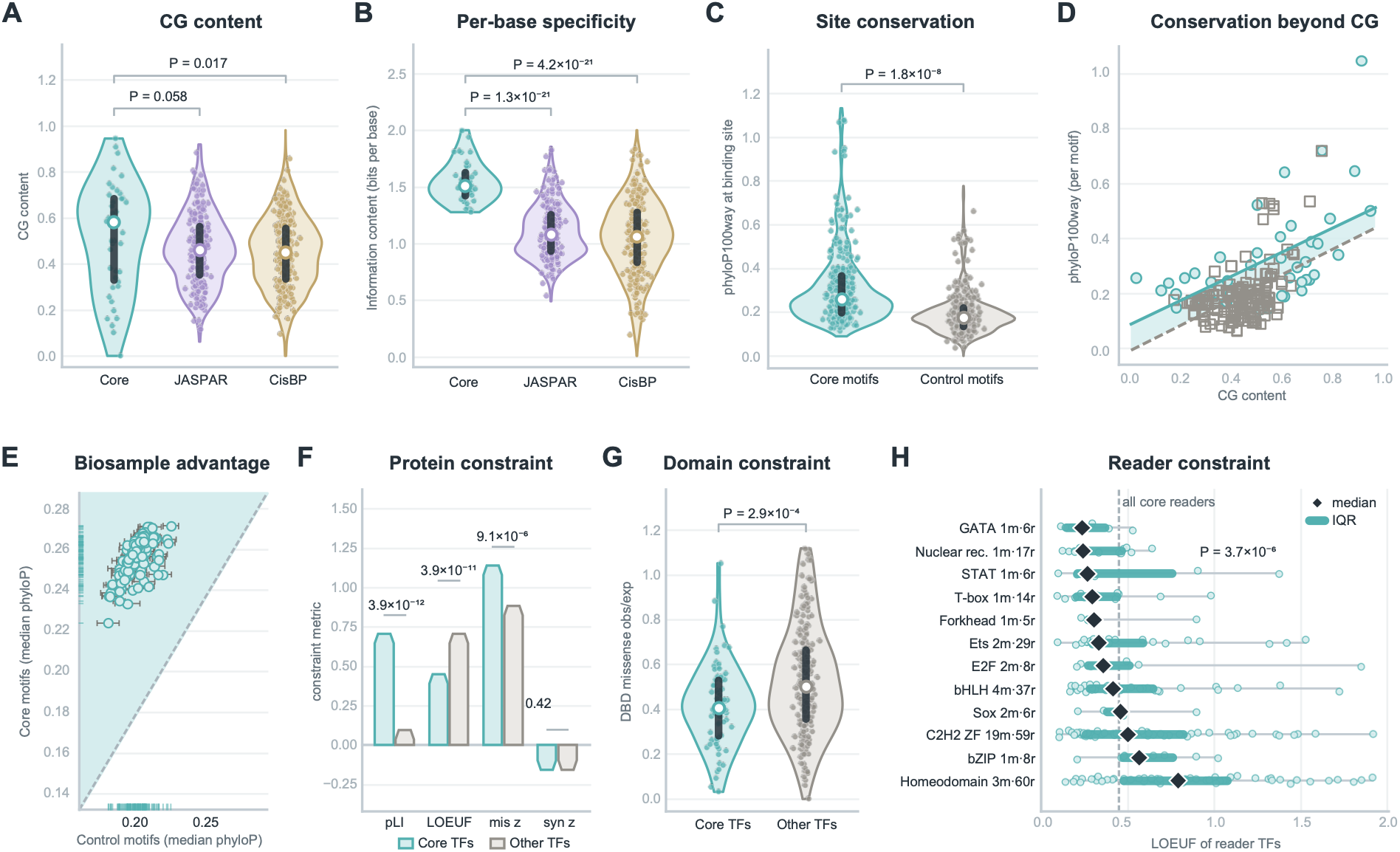
Core regulators are evolutionarily constrained at the motif, binding-site and protein levels. Core is teal, JASPAR violet, CisBP ochre, and control motifs and other TFs grey. **(A and B)** CG content and information content per base of the 43 core motifs against the JASPAR (n = 841) and CisBP human (n = 6,607) catalogues. **(C)** phyloP100way conservation of FIMO-mapped binding sites over the motif bases alone, core versus control motifs. The control motifs are random JASPAR motifs with no similarity to any core motif, drawn as ten independent sets of 43 (Methods); the control violin pools the 141 distinct motifs of the ten sets, each motif once (median 0.257 vs 0.174). Each motif is summarized by the median phyloP of its sites, and the test compares these per-motif values (43 core versus 141 control motifs, two-sided Mann– Whitney; P shown in the panel). Tested against each of the ten control sets separately (43 versus 43), P ranges from 1.9 × 10⁻⁶ to 8.6 × 10⁻⁴. **(D)** Per-motif median phyloP against CG content with per-set least-squares fits; the slopes are similar (+0.430, +0.454) and the shaded band is the CG-adjusted core advantage (+0.085; p = 1.4 × 10⁻⁵, common-slope ANCOVA; +0.059 to +0.105 across the ten control sets, all p < 0.05). **(E)** The advantage holds in 116 of 116 biosamples (p = 1.2 × 10⁻³⁵, sign test); one point per biosample, x = mean over the ten control sets with s.d. error bar, shaded field marks core > control; the core median exceeds every individual control set in all 116 biosamples. **(F)** gnomAD constraint of the core reader TFs versus 1,338 other human TFs. The 43 core motifs match 287 entries with 254 unique TFs after removing heterodimers in JASPAR by TOMTOM; the 250 of these with gnomAD constraint estimates are plotted. Synonymous z is a negative control. **(G)** Regional missense constraint within DNA-binding domains, 81 core reader TFs versus 268 other TFs: the subsets of the two groups in (F) that carry a UniProt-annotated DNA-binding domain covered by a gnomAD regional missense-constraint segment (Methods). **(H)** LOEUF of all 254 reader TFs grouped by the dominant family of the core motifs that recruit them, lower being more constrained; labels give motif reader TFs, diamond = median, bar = interquartile range (IQR), dashed line = median across all core readers (p = 3.7 × 10⁻⁶, Kruskal–Wallis).

At the motif level, the information content per base of the 43 motifs was higher (mean 1.546 bits) than the full JASPAR vertebrate (1.108 bits; p = 1.3 × 10^-21^) or CisBP human motifs (1.048 bits; p = 4.2 × 10^-21^; Fig. 3A,B). They are modestly CG-rich relative to both catalogues, significantly so against CisBP (median 0.582 versus 0.461 and 0.450; p = 0.058 and p = 0.017) but not JASPAR, and the enrichment does not account for their conservation (below). Thus, the compact vocabulary consists predominantly of information-rich recognition patterns rather than broadly occurring low-complexity sequence features.

At their predicted genomic occurrences, the core motifs localized to more evolutionarily conserved positions than control motifs. The control motifs are JASPAR motifs drawn at random after excluding any motif that resembles a core motif, in ten independent sets of 43 (Methods). Across the 100-vertebrate phyloP (*33*) alignment, all core-motif occurrences had higher conservation scores over the motif bases alone (median 0.257) than control-motif occurrences (median 0.174 over the 141 distinct control motifs of the ten sets; p = 1.8 × 10^-8^ across the per-motif medians; per-set control medians 0.169–0.191, p = 1.9 × 10^-6^ to 8.6 × 10^-4^; Fig. 3C).

Conservation rises with CG content at a similar rate in both motif sets (slopes +0.430 and +0.454), yet core motifs remain more conserved at matched CG content (+0.085 phyloP; p = 1.4 × 10^-5^, common-slope ANCOVA; +0.059 to +0.105 across the ten sets, all p < 0.05; Fig. 3D), so the advantage is not a CpG-island effect. It held in 116 of 116 biosamples against the mean of the ten control sets, and against every individual set (Fig. 3E), indicating a genome-wide property of these sequences rather than a biosample-specific artifact. The core motifs therefore preferentially occupy promoter positions that have been maintained during vertebrate evolution, independent of their modest CG enrichment.

At the protein level, candidate TFs that recognize the core motifs are among the most mutation-intolerant in the human genome. Relative to all other human TFs, the 250 of 254 core-matched TFs carrying gnomAD (*34*) constraint estimates showed dramatically stronger constraint across all three standard metrics: a probability of loss-of-function intolerance (pLI) of 0.706 versus 0.094 (p = 3.9 × 10^-12^), a loss-of-function observed/expected upper-bound fraction (LOEUF) of 0.449 versus 0.705 (p = 3.9 × 10^-11^), and a missense z-score of 1.139 versus 0.884 (p = 9.1 × 10^-6^). Synonymous z, which is not under coding selection and thus as a negative control, showed no difference (−0.159 versus −0.155; p = 0.42), excluding coverage, gene-length or mutation-rate artifacts (Fig. 3F). The LOEUF association also remained under a motif-level permutation that preserved the number of candidate TFs associated with each motif (Methods).

Constraint extended specifically to annotated DNA-binding domains. Core-motif-matched candidate TFs were more depleted of missense variation within these domains than other TFs (median missense observed/expected ratio 0.405 versus 0.501; p = 2.9 × 10^-4^; Fig. 3G), showing that evolutionary selection acts most strongly on the very residues that read the core motifs.

Constraint was not uniform across reader families. Assigning each core motif to the dominant family among its matched TFs and pooling all readers of each family, the 254 reader TFs differed markedly (p = 3.7 × 10^-6^, Kruskal–Wallis; Fig. 3H): GATA, nuclear receptor and STAT readers were the most constrained (median LOEUF 0.233–0.264) and homeodomain readers the least (0.789). The core motifs therefore recruit readers spanning the full constraint spectrum rather than a single intolerant family.

### Core motif occurrences are enriched at fine-mapped eQTLs and in TF-bound regulatory regions

We next asked whether the core motifs sit at genomic positions with independent evidence of regulatory function. We used the fine-mapped GTEx eQTL sets (*35*) released with Enformer (*10*), which provide tissue-specific variants assigned high posterior probability (>0.9) of affecting gene expression. Because our framework treats chromatin as a gate on motif use, we counted motif occurrences only within promoter windows defined by the matched biosample’s Epimap concatenated promoter landscape. Twenty-six GTEx tissues could be paired with corresponding EpiMap biosamples, of which 16 contained more than 500 fine-mapped variants and were used for the primary statistical comparison (table S2, Methods).

Across those 16, fine-mapped variants overlapped core-motif occurrences far more frequently than occurrences of ten independent sets of 43 random JASPAR motifs with no similarity to any core motif (paired one-sided Wilcoxon p = 1.5 × 10^-5^ against the mean over the ten sets; Fig. 4A). The core-to-control difference was positive in all 16 primary comparisons and also in each of the 10 lower-variant-count tissue pairs (<500 variants) excluded from the formal test (Fig. 4B; table S2). Thus, within chromatin-defined active promoters, the shared motif vocabulary preferentially marks positions implicated by human genetic variation in transcriptional regulation.

**Fig. 4.**
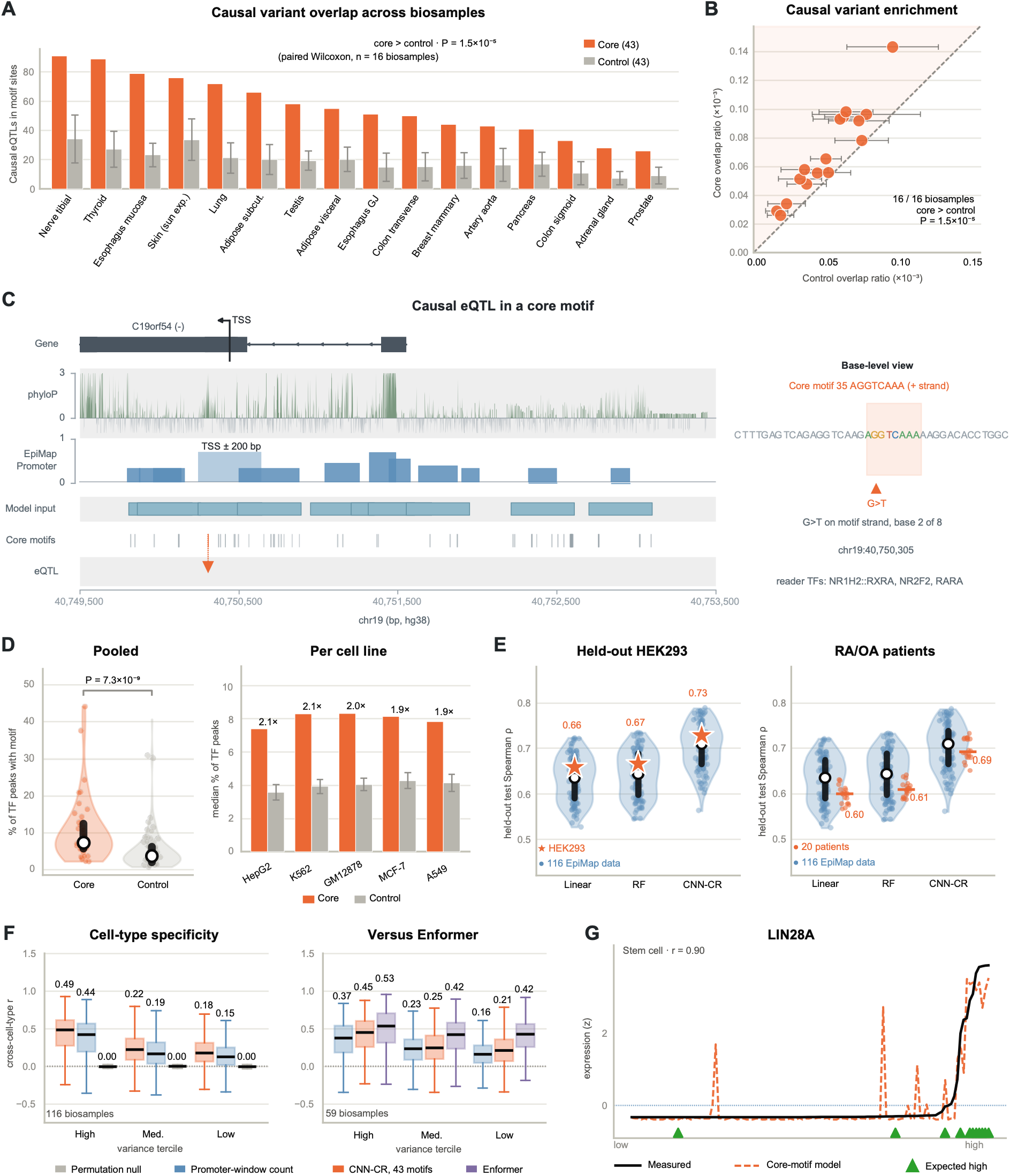
Core motifs mark functional variants, are broadly occupied, generalize to held-out cellular contexts, and predict cell-type-specific expression. **(A)** Fine-mapped causal eQTLs (GTEx, PIP > 0.9) overlapping the 43 core motifs (orange) versus control motifs (grey) in the 16 biosamples with > 500 causal variants. The control motifs are ten independent sets of 43 random JASPAR motifs with no similarity to any core motif (Methods); the grey bar is the mean over the ten sets and the error bar their s.d. (paired one-sided Wilcoxon p = 1.5 × 10⁻⁵ against the mean). **(B)** Causal-variant overlap ratio (overlaps / motif occurrences), core versus the mean over the ten control sets (error bars, s.d. over the sets); all 16 biosamples lie above the diagonal. **(C)** The C19orf54 promoter in the matched tibial-nerve biosample: gene model with TSS, phyloP conservation, EpiMap promoter track, model input windows, core-motif occurrences (the variant-containing occurrence in orange) and the fine-mapped causal variant (orange arrow); grey bands separate alternating tracks. Right, the 40-bp sequence with the motif, the base change and its candidate reader TFs. **(D)** Fraction of a TF’s TSS-proximal ChIP-seq peaks containing a motif match (ENCODE4, 1,151 TF–cell-line pairs). Left, pooled core versus control, the control violin pooling the 141 distinct motifs of the ten control sets, each once (median 7.2% vs 3.6%, one-sided Mann– Whitney p = 7.3 × 10⁻⁹; per set 1.6–2.4-fold, all p ≤ 1.8 × 10⁻⁴); right, per-cell-line medians, control shown as the mean over the ten sets with s.d. error bar (1.9–2.1-fold excess). **(E)** Held-out test Spearman ρ for linear, random-forest and CNN-CR models on the fixed 43-motif features; violins, the 116 EpiMap biosamples (open circle, median; black bar, IQR). Left, held-out HEK293 (orange star), at the 69th/63rd/65th percentile. Right, twenty patient synovial-fibroblast samples (orange dots; orange bar, median): 0.598, 0.621 and 0.694 versus EpiMap 0.636, 0.647 and 0.710. **(F)** Per-gene cross-cell-type Pearson r by expression-variance tercile, expression standardized within each biosample (Avsec et al.10); medians printed above the boxes. Left, all 116 biosamples: the 43-motif CNN-CR (orange), the promoter-window count baseline (blue) and the shuffled permutation control (grey), all scored on the same genes. Right, the 59 Enformer-covered biosamples on Enformer’s own held-out sequences (1,633 test genes), with CNN-CR retrained on Enformer’s gene partition so that neither model has seen them: promoter-window count (blue), the 43-motif CNN-CR (orange) and Enformer (purple). Each model is scored against the measurement it was trained to predict, Enformer against measured CAGE and the 43-motif CNN-CR against EpiMap RNA-seq; variance terciles are fixed for all three series, so the boxes cover identical gene groups. **(G)** Measured (black) versus predicted (orange) expression across the 116 biosamples, sorted by measured level, for LIN28A (r = 0.90); green triangles, biosamples expected to express it.

Furthermore, because every core motif is tied to TFs, each eQTL variant inside a core-motif occurrence can generate a concrete, testable mechanism. Figure 4C shows one such locus: an eQTL variant 138 bp from the C19orf54 TSS falls inside a nuclear-receptor core-motif occurrence, within an H3K27ac-marked promoter window in the matched tibial-nerve biosample; the motif’s bases are more evolutionarily conserved than the surrounding sequence (mean phyloP 1.70 versus 0.05), and its candidate reader factors include NR2F2 and RARA. Such loci therefore nominate a specific motif instance and a restricted pool of candidate TFs through which a regulatory variant might act. Two additional examples are shown in fig. S10.

We next examined whether core motifs are preferentially located within regions occupied by TFs (Methods). Across 1,151 ENCODE4 ChIP-seq data (*36*) from five cell lines, core motifs occurred in a larger fraction of TF binding peaks than did the ten sets of control motifs (median 7.2% versus 3.6% of peaks over the 141 distinct motifs of the ten sets, a 2.0-fold excess; one-sided Mann–Whitney p = 7.3 × 10^-9^). The enrichment was reproduced against every individual control set (1.6- to 2.4-fold and p ≤ 1.8 × 10^-4^) and in every individual cell line (1.9- to 2.1-fold against the mean of the ten sets; Fig. 4D, right). Together, the genetic and occupancy analyses show that the core vocabulary is preferentially positioned at promoter sites with independent evidence of regulatory activity.

### Core motif vocabulary generalizes to independent cellular contexts and predicts cell-type-specific expression

A defining prediction of a shared regulatory vocabulary is that the same motif set should remain informative in cellular contexts that did not contribute to its discovery. We therefore applied the fixed 43 core motif families to HEK293, an embryonic kidney line absent from the EpiMap discovery panel, and to twenty primary synovial-fibroblast samples from patients with rheumatoid or osteoarthritis (*37*) (Methods). Only the motif vocabulary was transferred: model parameters were fitted independently within each new context using its own chromatin and expression data. EpiMap chromatin-state annotations do not exist for these samples, so promoter windows were built from the pooled peaks of each sample’s own chromatin assays (ATAC-seq or DNase-seq with H3K27ac, H3K4me3 and H3K4me1, plus PRO-cap for HEK293), under the same window rules as for the 116 biosamples (Methods; Supplementary Methods S8).

In HEK293, models using the fixed 43-motif feature set reached held-out Spearman correlations of 0.659 for linear regression, 0.666 for random forest and 0.728 for PWM-initialized CNN-CR, placing HEK293 within the performance distribution (69th, 63rd and 65th percentile) of the original 116 biosamples (Fig. 4E; Methods). Across the twenty synovial-fibroblast samples, median correlations were 0.598, 0.621 and 0.694, respectively, compared with 0.636, 0.647 and 0.710 across the original EpiMap panel. Thus, the 43-motif vocabulary selected without either dataset retained similar predictive power in the new cellular contexts.

The chromatin-gating relationship established above (Fig. 1H) also transfers unchanged. Under the same chromatin-coverage baseline, promoter-window count alone, carrying no sequence information, predicts expression with Spearman ρ = 0.634 in HEK293 and a median of 0.565 across the twenty patient samples. Against this baseline the 43-motif model achieved gains of 0.025, 0.032 and 0.094 in HEK293, and median gains of 0.033, 0.056 and 0.129 across the 20 patient samples for linear, random forest, and CNN-CR models respectively. Thus, the first contextual layer is reproduced outside the discovery panel: chromatin determines which regulatory sequences are available, while the shared motif vocabulary contributes sequence-specific information within those active regions.

We next asked whether the shared vocabulary can predict expression variation of individual genes across biosamples. Following Avsec et al.10, expression was standardized within each biosample, a correlation between predicted and measured expression was computed for each gene across the 116 biosamples, and genes were split into three equal groups (terciles) by how variable their expression is across biosamples (Methods). A high correlation therefore requires the model to predict where a gene is up- or down-regulated between cell types, not merely its average level. Correlations were compared with a permutation control in which the correspondence between gene expressions and biosamples is shuffled.

Across all 116 biosamples, the 43 core motifs resolved cell-type-specific expression precisely where such regulation exists. These correlations run in the opposite direction to the rest of this paper: each gene is taken on its own and its predicted and measured expression are correlated across biosamples, so a high value means the model places that gene in the right cell types, not that it ranks genes correctly within one biosample. Predictions came from CNN-CR reading the 43 core motifs, with the biosample held out (Methods). For the most variable genes the median per-gene correlation was r = 0.486, with 92.1% of genes above zero and the most cell-type-specific genes reaching r ≈ 0.9; the medium- and low-variability groups gave positive but lower medians (0.222 and 0.179), whereas the biosample-label permutation control was centred near zero throughout (Fig. 4F, left). The 43-motif CNN-CR also improved on the chromatin-coverage baseline — the number of active promoter windows a gene receives, used as the single feature of a model carrying no sequence information — which on the same genes reached 0.436, 0.189 and 0.151 (paired one-sided Wilcoxon P = 4.7 × 10^-73^, 1.2 × 10^-41^ and 1.6 × 10^-107^), so the cell-type-specific signal is only partially predicted by how much open chromatin a gene has. The signal is strongest exactly where genes differ most between cell types, as expected if the shared vocabulary, read through each biosample’s own chromatin, encodes cell-type-specific regulation. We then repeated the comparison with Enformer on Enformer’s own held-out sequences. The 43-motif CNN-CR was retrained on Enformer’s published partition of the genome, and both models were scored on the 1,633 test genes, which neither had seen, in the 59 biosamples Enformer covers; each model was scored against the measurement it was trained to predict, Enformer against measured CAGE and CNN-CR against EpiMap RNA-seq (Methods). The 43-motif CNN-CR again improved on that baseline in all three groups (median r = 0.451 versus 0.374, 0.245 versus 0.234 and 0.210 versus 0.161; paired one-sided Wilcoxon P = 1.3 × 10^-19^, 3.3 × 10^-4^ and 4.6 × 10^-10^) and approached Enformer most closely among the most variable genes, the group in which cell-type-specific regulation is strongest (0.451 versus 0.534), remaining below it in the medium- and low-variability groups (0.245 versus 0.422 and 0.210 versus 0.424; Fig. 4F, right). A convolutional network reading forty-three motifs therefore comes close to a multi-hundred-million-parameter sequence transformer on variable genes.

The predictions tracked canonical lineage-restricted genes, peaking in the expected biosamples, such as the pluripotency factors POU5F1 (OCT4, r = 0.80) and LIN28A (0.90), the pan-hematopoietic marker PTPRC (CD45, 0.83), the T-cell marker CD4 (0.61), muscle MYOD1 (0.63), erythroid GATA1 (0.53) and the hepatocyte marker FOXA2 (0.54), nearly all in the high-variance tercile (LIN28A in Fig. 4G; further examples in fig. S11A–D); the second hepatocyte marker ALB is the exception at 0.13. Housekeeping genes, by contrast, were not tracked at all (GAPDH r = 0.06, TUBB 0.12, TBP 0.06, SP1 −0.06; fig. S11E–H), as expected of genes whose expression does not vary with cell identity. Together these results show that the 43 core regulators generalize to unseen cellular contexts and resolve cell-type identity at the level of high-variance regulatory genes.

### Core motifs show diverse, context-dependent reader usage

The analyses above establish the first contextual layer: chromatin determines which genomic copies of the shared motif vocabulary are available for regulatory use. We next tested the second. Once a core motif is available, does the same TF interpret it in every cell type, or can different compatible factors read the same sequence in different contexts?

Each core motif family is compatible with multiple TFs whose DNA-binding preferences are similar enough that sequence alone cannot identify the factor occupying a particular site (Fig. 5A). We therefore used ENCODE ChIP-seq across five cell lines to resolve occupancy experimentally, scoring each cell line at the core-motif copies that lie inside its own promoter windows (246,571 to 317,970 motif copies per cell line, 375,180 distinct copies in all). For each TF, binding was restricted to the strongest 10% of peaks within that factor’s own experiment, a within-factor percentile that keeps each factor’s own strongest sites without ever comparing one factor’s signal with another’s (Methods; peak selection in Supplementary Fig. S12). A gene was counted as bound by a factor when one of these peaks overlapped the motif copy in its promoter. Occupancy was then summarized by how widely a factor binds: the dominant binding reader of a motif in a cell line is the candidate factor bound at the largest number of that motif’s bound genes, provided it reaches at least 20 genes. Counting binding rather than ranking signal keeps the comparison free of the antibody, epitope-tag and experiment-specific differences that make ChIP-seq intensities incomparable between factors and between cell lines. Switching was likewise scored without reference to signal, between pairs of cell lines and only over those candidate readers that have ChIP-seq in both cell lines of the pair; a gene counts as switched when the set of these readers bound there in one cell line is not the set bound there in the other. A change of dominant reader can be seen only where a motif has a dominant reader in at least two cell lines, which 28 of the 43 core motifs do; 14 of those 28 changed dominant reader between lines, involving 31 distinct factors. A switch rate can be measured only where enough genes carry the motif and are bound in both cell lines of a pair, which brings 12 of the 43 above the 20-gene requirement; for the four motifs drawn in Fig. 5B the readers bound differed at 43% to 72% of the genes, and across all 12 at 3.9% to 72.2%, pooling to 53.1% over all cell-line pairs (25,608 gene comparisons) (Fig. 5C). Eight of the 12 changed dominant reader: four are drawn in Fig. 5b and the other four in Supplementary Fig. S12F.

**Fig. 5.**
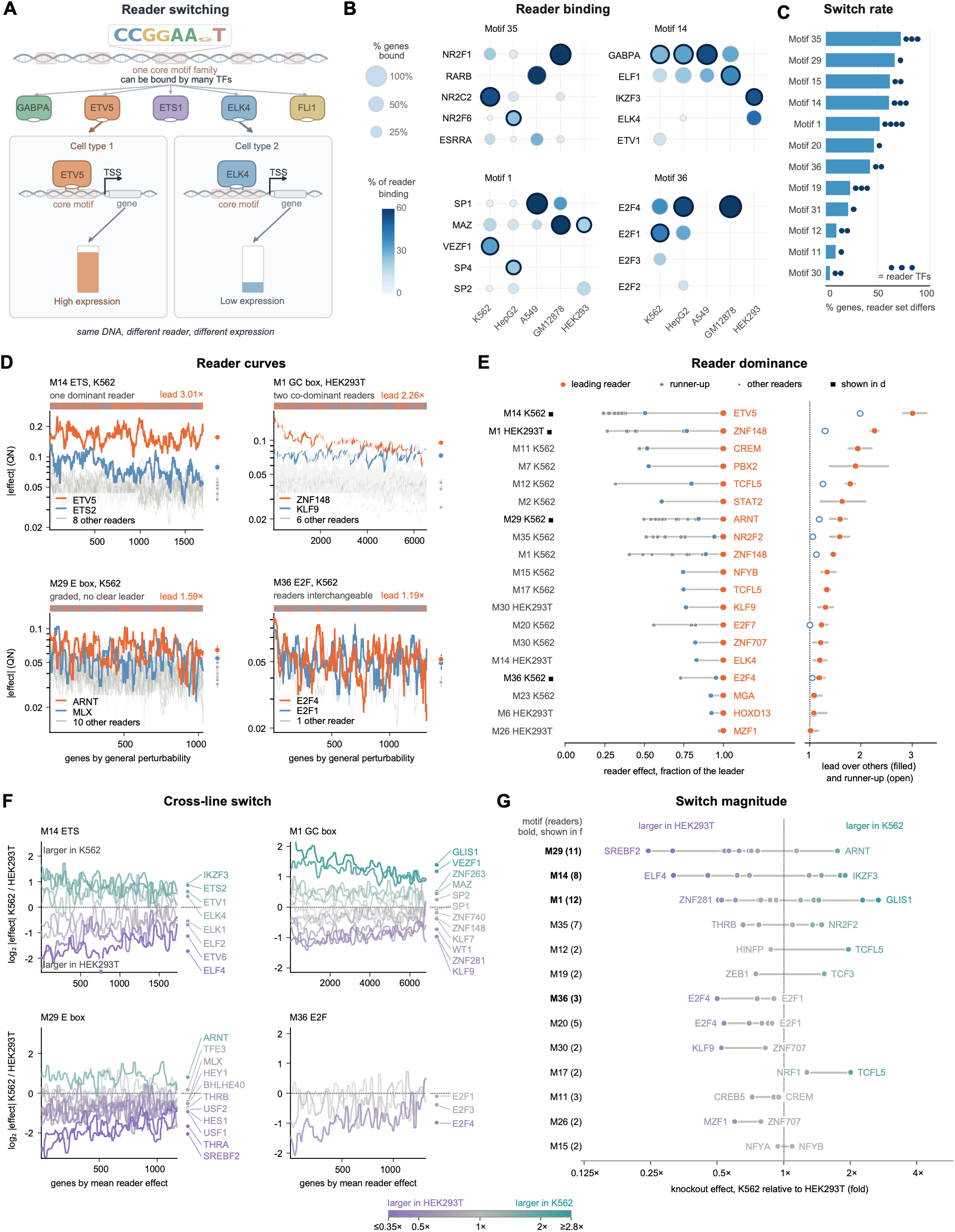
Core motif families exhibit diverse, context-dependent reader architectures. **(A)** Thesis: a core motif family is bound by many candidate reader TFs, and the same motif copy at the same gene is occupied by a different member of that pool in each cell type. **(B)** Reader binding at core-motif sites, for the four motifs whose dominant reader changes between lines and whose reader set turns over most. Rows, the motif’s candidate reader TFs, at most five per motif; columns, the five ChIP-profiled cell lines. Bubble size, the percentage of the motif’s bound genes at which that TF binds; shade, that TF’s share of all candidate-reader binding at the motif in that line; the two channels have the same numerator and differ only in the denominator. Dark ring, the dominant reader of that line, the TF bound at the most genes. No ChIP-seq signal is compared between factors or lines anywhere in (B) or (C). **(C)** Per-motif switch rate: the percentage of genes, bound in both lines of a pair, at which the set of bound candidate readers differs, pooled over all cell-line pairs and restricted to readers ChIP-profiled in both lines, for the 12 motifs with at least 20 such genes. Dots, the number of distinct dominant readers of that motif across the five lines; a cell line contributes one only when its most widely bound candidate reaches 20 of the motif’s bound genes. Motifs 29 and 20 carry a single dot beside a long bar: their leading reader is the same in every cell line while the readers around it change. **(D)** Reader curves within one cell line, four representative motif and cell line panels. Each curve is one reader’s quantile-normalized absolute knockout effect across the motif’s ChIP-defined target genes, all readers scored on the same genes (Methods). Orange, leader; blue, runner-up; grey, the other readers, 10th to 90th percentile shaded. Right strip, each reader’s median effect with a bootstrap 95% interval. Top strip, orange where the leader leads, blue where the runner-up does. Lead, the leader over the median of the other readers. **(E)** The same measurement for all nineteen motif-cell combinations. Left, every reader at its median effect as a fraction of the leader’s. Right, that lead over the other readers (filled dot, with its bootstrap 95% interval) and over the runner-up alone (open dot); 1.0 means no leader. Rows ordered by the lead; the four panels of (D) are marked. **(F)** Cross-cell-type comparison for the four core motifs whose readers spread most widely. Every candidate reader of the motif is one curve: the log2 ratio of its absolute knockout effect in K562 to its effect in HEK293T, over one fixed gene set (the union of the motif’s ChIP-defined target genes across its candidate readers) scored in both screens. Curves above zero are readers that matter more in K562, below zero readers that matter more in HEK293T, and readers with no cross-line preference lie on the line. Colour runs violet to teal with the key between (F) and (G), saturating at a 2.8-fold median ratio; curves are smoothed over 2.5% of the genes for display. The marginal at the right of each panel names every reader at its median ratio. The nine motifs not drawn here, with the absolute knockout magnitudes behind their ratios, are in fig. S14. **(G)** Each reader’s cross-line preference, one lane per motif, for the 13 motifs that qualify. Dots are readers, coloured as in (F), with the two extremes named and the bar spanning the motif’s readers; motifs drawn in (F) are in bold. A motif with dots on both sides of the dashed line has readers that favour different cell lines; a motif with all dots on one side differs in overall knockout size rather than in reader identity.

These changes generally occurred among TFs with closely related DNA-binding preferences. A nuclear-receptor motif is bound most widely by NR2C2 in K562, NR2F6 in HepG2 and NR2F1 in GM12878; an ETS motif by GABPA in K562, HepG2 and A549 but by ELF1 in GM12878 and IKZF3 in HEK293; a GC-box motif by VEZF1, SP4, SP1 and MAZ across the five lines; and an E2F motif by E2F1 in K562 and E2F4 in HepG2 and GM12878 (Fig. 5B). The turnover is not confined to the leading factor. USF1 is the most widely bound reader of an E-box motif in all four lines in which it can be called, and E2F8 leads a second E2F motif in all three of its lines, yet the readers present at a gene still differ between lines at 67% and 46% of genes respectively (Fig. 5C). Thus, motif identity constrains the pool of compatible factors but does not determine which of them occupies the motif in a particular cellular context. This provides direct evidence for context-dependent reader selection.

Occupancy alone, however, does not show which TF carries the motif’s function. We therefore used two genome-scale Perturb-seq datasets (*38*) in K562 (Replogle (*39*)) and HEK293T (X-Atlas/Orion atlas (*40*)). We quantile normalized the 8,236 genes they share (Methods). For each motif and cell line, target genes were defined by the presence of that motif within any ChIP-seq peak of any candidate readers (i.e. matched TFs) in the gene’s promoter (Methods). All candidate readers of a motif were then evaluated across the same union of target genes. Nineteen motif-cell-line combinations contained at least two perturbed readers and at least 100 target genes.

Ranking the readers of a motif in the same cell line by their median absolute knockout effect, the leading reader’s advantage over the median of the others varies from 1.02 to 3.01 fold (Fig. 5D,E, fig. S13). Some motifs have one dominant reader. The ETS motif in K562 is led by ETV5, whose effect is 1.98-fold that of the runner-up ETS2 and 3.01-fold the median of the other nine readers, and which is the larger of the two at 71% of the 1,694 target genes (paired Wilcoxon P < 10^-99^). Other motifs have two co-dominant readers. The GC box in HEK293T is led by ZNF148 with KLF9 close behind, 1.30 fold apart, and the two together stand 2.26 fold above the remaining six readers. In the rest no reader stands out. The twelve readers of the E box motif form a graded band with no step in it, and the three E2F readers of an E2F motif are level, the leading pair separated by 1.05 fold (P = 0.63). Reader interpretation is therefore not necessarily one-factor-per-motif: some motifs are functionally dominated by one or two readers, whereas others are distributed across several compatible factors.

We then used the two screens together to check whether a reader’s effect is specific to a certain cell type. For each core motif we took every candidate reader with an effective knockdown in both screens and scored all of them on one fixed gene set: the genes that carry a copy of the motif inside a ChIP-seq peak of any of its candidate readers in K562 or HEK293 (at least 300 genes). The ChIP-seq data therefore define only the genes; the effects come from the two screens. Every reader of a motif is therefore measured on the same genes in both cell types, and any difference between them comes from the perturbation rather than the gene set. For each reader we took the absolute knockout effect on each of those genes in both cell lines and formed the ratio of the K562 value to the HEK293T value, gene by gene. Averaged in log2, that ratio says which of the two cell lines a reader’s knockout matters more in, and the larger its absolute value, the more strongly the reader favors one line. We call that absolute value the reader’s switch magnitude. Thirteen core motifs had at least two readers meeting these conditions. In seven of the thirteen the readers fall on both sides of the line, so that for the same motif some readers matter more in K562 and others more in HEK293T; in the remaining six every reader leans the same way, which is a difference in overall knockout size rather than a change of reader. Figure 5F draws every reader of the four motifs whose readers spread most widely: the ETS motif divides four readers towards K562 against four towards HEK293T, the GC box five against seven and the E-box motif two against nine, whereas the three E2F readers of motif 36 all lean to HEK293T. Across the thirteen motifs the spread between a motif’s most K562-leaning and most HEK293T-leaning reader ranges from 1.2-fold (the NFY motif, NFYB and NFYA) to 7.2-fold (the E-box motif, ARNT and SREBF2) (Fig. 5G). The switches therefore differ widely in size, and not every motif shows one. The remaining nine motifs, together with the absolute knockout magnitudes behind their ratios, are in fig. S14.

Together, the occupancy and perturbation analyses support the second contextual layer of the model: once chromatin makes a motif instance available, the TF that interprets that sequence can differ between cell types. The shared DNA vocabulary therefore remains fixed while cellular context acts twice—first by determining which motif instances are usable and then by determining which compatible factor reads them.

## Discussion

A compact promoter-motif vocabulary captures much of the regulatory information associated with gene expression across diverse human cell types. Starting from independently learned motif representations in 116 biosamples, we identified an operational set of 43 core motif families that retains most of the predictive information carried by much larger biosample-specific vocabularies. The exact boundary is continuous rather than discrete, so the central result is compressibility rather than the number 43 itself. Despite representing only a small fraction of the motifs learned in individual biosamples, the core set retains approximately 96% of full-vocabulary predictive performance and remains informative in cellular contexts excluded from its discovery. Importantly, the same models built on random sets of 43 JASPAR motifs performed significantly worse, suggesting that the identified core motifs are not computational artifacts. These results suggest that a substantial component of promoter regulation can be represented by a compact shared vocabulary rather than requiring a largely distinct set of regulatory motifs in every cell type.

Our results further suggest that this shared vocabulary acquires cell-type specificity through two context-dependent operations. First, chromatin state gates motif use. The same promoter motifs carry little additive predictive information when counted without regard to chromatin, but become strongly predictive when restricted to regulatory regions active in the corresponding biosample. What is linearly predictable appears to come mainly from accessibility itself: an additive model of motif counts improves only marginally on the chromatin-coverage baseline (median held-out ρ = 0.675 versus 0.641), whereas CNN-CR, which can combine motif occurrences non-additively, reaches 0.743 on the same biosamples. This suggests that the contribution of motif identity and combination is largely non-linear, as expected if these motifs are read cooperatively rather than independently. Second, reader identity determines how an available motif is interpreted. Individual core motif families are compatible with multiple TFs with closely related DNA-binding preferences, which can shift further when two such factors bind DNA as a pair (*41*), and ChIP-seq shows that the factor occupying a motif family frequently changes across cellular contexts; genome-scale perturbation further supports context-dependent differences in the functional contribution of these readers. Thus, the underlying DNA vocabulary can remain largely shared while cellular context changes both which copies can be used and which proteins interpret them.

The compact core does not imply that the remainder of the motif landscape is unimportant. Indeed, a substantial fraction of the highest-weighted de novo motifs in individual biosamples lack matches to JASPAR, yet these database-unmatched motifs rarely recur broadly across cell types. Only 19 of 2,623 unmatched motif families reached the recurrence criterion used to define the core, substantially fewer than among catalogued motifs. This organization suggests a two-scale regulatory vocabulary: a broadly reused core that provides a common transcriptional basis, accompanied by a much larger tail of context-restricted sequence features that may refine regulation in particular cellular states. The regulatory architecture may therefore be less a choice between universal and cell-specific motifs than a combination of a shared backbone and context-specific refinements.

The two contextual layers may themselves be coupled rather than strictly hierarchical. Twelve of the 43 core motif families resemble sequence motifs previously associated with histone modifications, and two also resemble DNA-methylation-associated motifs (table S5)26– 28. This overlap raises the possibility that some DNA elements participate on both sides of the chromatin gate: chromatin state determines whether a motif is available for binding, while proteins recognizing particular motifs may in turn recruit or stabilize chromatin-modifying activities that alter accessibility or histone state. Pioneer factors, which engage their motifs in closed chromatin and open it for other factors, are the clearest precedent for a reader acting on the gate itself (*42*). In this view, chromatin gating and reader interpretation are analytically distinct but potentially connected by feedback. Establishing such feedback will require direct perturbation of motif instances and their candidate readers together with measurements of chromatin state, but the overlap with epigenetic motifs provides a testable route toward linking the two layers.

More broadly, the results suggest a way to think about how regulatory information is encoded and processed. The combinatorial sequence space of DNA is enormous, yet cells need not occupy that space by evolving an independent regulatory vocabulary for every cell type or environmental state. Instead, a relatively compact collection of recognition sequences can provide a stable information substrate, while chromatin and the available TF repertoire dynamically determine how that information is decoded. This separation between storage and interpretation expands the regulatory output available from a limited vocabulary: the same sequence can be silent in one context, active in another, and interpreted by different factors in still another. Such an architecture may also provide evolutionary flexibility. Mutations can alter motif occurrence or affinity in cis, whereas changes in chromatin programs, TF abundance or reader interactions can alter regulation in trans without requiring wholesale rewriting of the underlying motif vocabulary. Developmental programs and responses to environmental or signaling cues could therefore evolve by modifying both the encoded sequence and the machinery that interprets it.

This distinction also has practical implications for noncoding variation, which accounts for the majority of common disease-associated variants and is concentrated in regulatory DNA that is active in only a subset of cell types³⁶. A variant disrupting a motif does not necessarily implicate the TF whose name is attached to the canonical motif; its consequence should depend on whether that motif instance is available in the relevant cell type and which compatible factor reads it there. Conversely, the same sequence change could have different effects across tissues because either chromatin gating or reader identity differs. A compact motif vocabulary therefore provides an interpretable intermediate between raw sequence and cellular phenotype: variants can first be mapped onto shared motif families and then resolved through cell-type-specific chromatin and reader information.

The compactness of the motif vocabulary may also inform synthetic regulatory design. Rather than searching an effectively unconstrained sequence space, synthetic promoters could in principle be assembled from a smaller set of recurrent motif families and tuned through their combination, abundance and cellular context. In this framework, broadly shared core motifs would provide reusable regulatory building blocks, whereas context-restricted motifs, chromatin state and reader availability could modulate when and where those elements are active. Whether this reduced representation is sufficient for predictable design remains to be tested experimentally, particularly with respect to motif spacing, orientation, copy number and higher-order interactions, but it suggests a tractable route from descriptive regulatory genomics toward programmable control of gene expression.

The scope of the present study nevertheless remains primarily promoter-centered, and the 43 families should be regarded as an operational compact representation rather than an exhaustive regulatory alphabet. Reader-function analyses are also limited by the cellular contexts in which sufficiently dense ChIP-seq and perturbation data coexist, and population-level occupancy and transcriptome perturbation do not yet establish reader identity at every individual motif instance, which single-molecule footprinting is able to resolve directly (*44*). Extending the framework to distal regulation, where the rules relating sequence to activity are less settled than at promoters (*45*), and testing motif–reader–chromatin relationships with matched, site-resolved perturbations should reveal where this compact representation holds and where additional regulatory vocabulary is required.

Together, these results support a model in which human cells reuse a compact motif vocabulary while generating regulatory diversity through differential access and interpretation. Chromatin determines which encoded instructions are available; reader identity determines how those available instructions are interpreted; and a longer tail of context-restricted motifs may provide additional local refinement. Regulatory complexity can therefore arise not only from what is written in DNA, but from how a shared sequence vocabulary is selectively exposed and dynamically read. METHODS

## Materials and Methods

The procedures below are described at the level needed to reproduce them. Where a procedure cannot be specified briefly (regulatory-window construction, motif de-replication and clustering, derivation and robustness of the core families, control-set construction, the eQTL tissue pairing, the external-validation window construction, the cross-cell-type analysis and the Perturb-seq processing), the complete parameter-level description is given in Supplementary Methods. Throughout, “core motif families” refers to the operational set of 43 sequence families, “candidate TFs” to proteins matched by motif similarity, and “readers” to candidate TFs supported by occupancy or functional evidence in a cellular context. All code, and the data behind every figure panel (organized as data_Fig1a, data_Fig1b, etc.), are deposited on GitHub (Code and data availability).

### Expression data and regulatory sequences

RNA-seq expression (log2 FPKM) and biosample-specific promoter and enhancer peak annotations for 340 human biosamples were downloaded directly from the EpiMap release (*23, 46*) without reprocessing (table S3). These were reduced to 116 non-redundant biosamples by removing redundant biological replicates, sex-specific counterparts and two samples failing quality control (table S4). Transcription start sites are FANTOM5 CAGE peaks (*47*), taking the strongest strand-specific peak per gene (hg38). Analysis was restricted to 18,356 protein-coding autosomal genes. EpiMap coordinates were lifted from hg19 to hg38 with liftOver (*48*). Each gene received up to ten promoter sequences from TSS ± 10 kb: a mandatory core promoter of ± 200 bp around the CAGE TSS, plus up to nine EpiMap promoter peaks lying at least 50% inside the window, the nine widest when more were available. Each gene also received up to ten enhancer sequences, which are not confined to any window around the gene. EpiTensor-predicted (*24*) three-dimensional contacts (interaction score > 10; 655,133 promoter–enhancer pairs) were anchored to the gene where an EpiMap promoter peak overlapped the contact’s promoter region by > 50% (BEDTools (*49*)); the contact’s distal region was retained as an enhancer where an EpiMap enhancer peak of that biosample overlapped it by > 50%, the ten widest when more qualified. Every sequence was standardized to 400 bp by symmetric expansion or contraction about the peak centre. Missing sequences were zero-padded, so every gene enters the model as twenty 400-bp slots (fig. S1; Supplementary Methods).

Because every gene receives the mandatory TSS-centred window, promoter-window count has a minimum of one; additional windows reflect biosample-specific epigenomic peak calls. Promoter-window count was therefore used as a simple summary of chromatin-defined regulatory coverage in the relevant analyses, rather than as a sequence feature.

### CNN-CR architecture and training

CNN-CR is a multi-branch PyTorch (*50*) network (Fig. 1A). The twenty 400-bp sequences of a gene are processed by twenty weight-independent branches. Within each branch, a first convolutional layer of 256 filters (width 8 bp, no bias term) feeds two sub-modules. A motif-activation module averages each filter’s raw activation over the 393 positions of the window, with no activation function, threshold or binarization, giving a 256-dimensional motif-activation vector (Supplementary Methods S1). A second module (ReLU and max-pooling of the first-layer activations, three further convolutions of 64 filters each followed by max-pooling, two dense layers of 256 units with ReLU and dropout 0.15, and a final linear layer) outputs a matching 256-dimensional vector of per-motif contextual coefficients. Their dot product, multiplied by one trainable scalar per branch, is summed over branches to predict expression (7,719,700 parameters; the contextual coefficients of a branch are set to zero when the gene has no sequence for it). Training and evaluation used a fixed hold-out design per biosample: 10% of genes were set aside once as the test split and never used for training or model selection; the remaining 90% were divided into ten folds, and ten models were trained, each on nine folds with the tenth fold as the validation set for early stopping: training stopped when the validation loss had not improved for 10 consecutive epochs, or at 100 epochs, and the weights at the stopping epoch were used. Every test gene was therefore predicted by ten models, none of which saw it, and the reported prediction was the median of the ten; held-out accuracy was the Spearman correlation between these predictions and measured expression across the test genes (1,160 models across the 116 biosamples). Training used Adam (learning rate 10⁻⁴, no weight decay), batch size 64 and an MSE loss to which L2 penalties on the two dense layers and the contextual-weight layer (coefficients 0.1, 0.2 and 0.3) were added. The promoter-only variant has the same architecture with ten branches, one per promoter sequence. Enformer (*10*) was compared on the 59 biosamples with published predictions, using its pre-trained CAGE tracks over ± 100 kb inputs; both models were scored on the same held-out test genes.

As a positive control, CNN-CR was trained on simulated data: dinucleotide-shuffled promoters with known motifs re-embedded at observed frequencies, and expression generated as the sum of the embedded motif counts with a single weight per biosample, chosen so that the signal has the standard deviation of the real log2 FPKM, centred on the real mean and plus Gaussian noise (σ = 0.05); ten models per biosample were trained with the production settings, and motif recovery was scored by TOMTOM between the extracted first-layer PWMs and the embedded motifs, at a threshold rescaled to the number of motifs searched so that it is as strict as the E ≤ 1 cutoff used against JASPAR comparisons below (fig. S5; thresholds and counts in Supplementary Methods S1).

### Motif extraction, de-replication and annotation

For every first-layer filter, which represents a motif, a position weight matrix (PWM) was built from the 1,000 highest-activation subsequences across training genes. The importance of a motif in a biosample, termed its contextual weight, was defined as the median, over the genes in which the product exceeds half of its maximum across genes, of the product of the filter’s window-averaged activation and its contextual coefficient from the second module in the CNN-CR model. The 750 highest-ranked motifs per model were retained, and those with information content > 10 bits were kept.

Redundant motifs were then merged in two rounds of graph clustering, first within each model and then across a biosample’s ten models. In each round the motifs were compared all-versus-all with TOMTOM (*32*) and every significant match (E ≤ 0.01 and p ≤ 0.01) formed an edge weighted by its E-value; two motifs were treated as similar when the summed E-value of the Dijkstra shortest-path connecting them was ≤ 0.01. Groups were grown greedily, a motif joining a group only when its path distance to every current member was within this bound, so that each group was a tight clique of nearly identical PWMs rather than a loosely connected component. Each group was represented by the member closest to the component-wise median PWM. The cross-model round retained only groups found by at least two of the ten models. This yielded a median of 705 promoter and 684 enhancer motifs per biosample (Supplementary Methods).

Motifs were annotated by TOMTOM against JASPAR 2022 CORE vertebrates non-redundant (*25*), CIS-BP2 and EpiMotifs (*26–28*). For the cell-type-specific analyses (Fig. 1E–H), these annotations are descriptive only. For the core-motif derivation (Fig. 2A,B), a within-biosample motif group was carried forward only when at least one member matched a JASPAR motif (E ≤ 0.05, p ≤ 0.01) whose TF was expressed in that biosample (log2 FPKM > 1.66), and groups sharing an expressed TF annotation were merged as the same factor. Motifs with no catalogue match were excluded from the core-motif derivation and analysed separately in fig. S9. The expressed-TF annotations were used again to define the candidate readers of each core motif (Fig. 3F–H and Fig. 5).

### Motif counting, chromatin gating and linear models

Motif occurrences were mapped with FIMO (*29*). Each PWM was stored in one orientation with FIMO scanning both the sequence and its reverse complement, so one PWM captured occurrences of the motif on either strand. All motifs were scanned in this way at the FIMO default p < 10⁻⁴. Motif occurrences were counted in three ways: every hit across the entire TSS ± 10 kb of the gene (the 20-kb sequence scanned in full, with no chromatin information), hits inside the biosample’s EpiMap peaks (the 400-bp promoter and enhancer windows), or peak-overlap-weighted, in which a hit under N overlapping peaks counts N times. The motif vocabulary, genes and model class were held fixed across these counting rules, so the comparison isolates the effect of chromatin restriction on the information carried by the same sequence patterns. Downstream count-based models used the peak-overlap-weighted representation, which gave the highest predictive accuracy.

The linear models used one feature per motif, the peak-overlap-weighted count of that motif’s occurrences in the gene’s regulatory sequences, and predicted log2 expression by ridge regression (RidgeCV; features z-scored with a scaler fitted on the training genes only). Each model was fitted on a fixed 90% training split of the genes and scored by Spearman correlation on the held-out 10%; the same split was used for every feature set with each biosample, so that de novo and JASPAR features, and promoter, enhancer and combined feature sets, were compared on identical genes by paired Wilcoxon tests across the 116 biosamples (Fig. 1H). Throughout, ‘split’ denotes this fixed 90/10 train/test partition and ‘fold’ the ten cross-validation folds within the training split.

### Derivation of the 43 core motif families

The cross-biosample derivation started from a reduced, expression-filtered set of representative PWMs built for each biosample independently of the non-redundant vocabularies described above (which are used only for Fig. 1E,H and as the full vocabularies of Fig. 2E). Per biosample, the 2,560 promoter filters of each of the ten models were filtered to a total information content above 10 bits, and the 750 highest-weighted survivors of each model were kept, giving 7,500 PWMs per biosample. These were compared against themselves with TOMTOM (Pearson distance, minimum overlap 7 bp; matches at E ≤ 0.01 and p ≤ 0.01) and against JASPAR 2022 (matches at E ≤ 0.05 and p ≤ 0.01, used for annotation), and grouped by Dijkstra shortest-path clustering on the self-TOMTOM similarity graph, a motif joining a group only if its path distance to every current member was ≤ 0.01. A group was retained only when at least one of its JASPAR-matched transcription factors was expressed in that biosample (log2 FPKM > 1.66 in the biosample’s own RNA-seq), so that each representative corresponds to a factor that can act in that cellular context (median 329 annotated groups per biosample). Groups annotated with the same expressed TF were then merged, transitively (table S6), so that one representative stands for each expressed factor rather than for each PWM variant of its site; this annotation-based merge, accounts for most of the reduction (median 73 representatives per biosample, range 45–97; 8,441 over the 116 biosamples), and 47% of the representatives are merges of two or more similarity groups. Because merging is transitive, a representative can combine groups annotated with different TFs that share one name. The member closest to the group’s component-wise median PWM (Pearson distance) was taken as the representative. Cross-biosample clustering. The 8,441 representative promoter PWMs from all biosamples were then compared all-versus-all with TOMTOM (E ≤ 0.5, p ≤ 0.01, overlap ≥ 7 bp). Edges were drawn only between motifs from different biosamples, weighted by E-value, and families were extracted by the same greedy shortest-path clustering used within biosamples, here with the lenient distance bound of 0.5 so that the same motif family groups across biosamples despite biosample-to-biosample PWM variation. Families spanning ≥ 25 distinct biosamples define the 43 core motifs (data S1). All JASPAR 2022 matches of the 43 PWMs were kept (TOMTOM E ≤ 1 and p ≤ 0.01, the criterion also used for the control-motif exclusion below). They give 324 motif–TF associations spanning 287 TF entries and 254 distinct TFs, with DNA-binding-domain families assigned through CIS-BP.

The recurrence cutoff was evaluated in two complementary sensitivity analyses (fig. S8; table S1; Supplementary Methods). First, the entire derivation was re-run on 100 random subsamples of 93 of the 116 biosamples under a proportionally scaled cutoff (≥ 20 of 93). The recovered sets had a median of 45 families (interquartile range 42–46); a median of 36 of the 43 full-panel families (range 32–42) were called core again in each replicate, and 40 of the 43 kept at least half of their membership (Jaccard index ≥ 0.5 to their best-matching replicate family; median Jaccard 0.79), the three exceptions being families that were recovered but split in two (table S1b,c). The set is therefore reproducible under resampling of the biosamples, which is the property the cutoff must have; the loss that does occur is concentrated in families at the cutoff boundary, as any threshold requires. Second, the cutoff was swept from ≥ 10 to ≥ 45 biosamples, giving strictly nested sets of 177 to 14 motifs. Each set was re-benchmarked under the identical protocol and gene split of Fig. 2E; held-out accuracy stayed within 5% of the 43-motif result (linear ρ 0.604–0.660, random forest 0.607–0.664), and the JASPAR TFs matched by every set are a superset or subset of the 287 matched by the 43 (table S1a). These analyses treat 43 as an operational representation of a stable, compact motif basis rather than a unique biological boundary.

For the uncatalogued-motif analysis (fig. S9), the derivation was repeated with one change: the JASPAR-expression filter above was replaced by a reproducibility requirement, a within-biosample group being carried forward when it contained motifs from at least three of the ten independently trained models, so that motifs without a catalogue match, including low-complexity motifs, travelled through the identical clustering with all similarity thresholds unchanged. The resulting 19 broad families were scored by their within-biosample percentile ranks of contextual and ridge weight.

### Benchmarking and controls

The full per-biosample vocabularies, the 43 core motifs and the control motifs were each evaluated by three model classes on identical fixed 90/10 gene splits. The classes are RidgeCV regression, a random forest (200 trees, sqrt features, minimum leaf 2) and CNN-CR with the PWMs installed as first-layer initialization (4 × 8 kernels, zero-padded to 256 filters). Control motifs were drawn from the 841 JASPAR 2022 CORE vertebrate motifs after excluding every motif with a TOMTOM match to any core motif (E ≤ 1 and p ≤ 0.01) and every motif in the same family of a core motif; the 149 motifs that remained form the candidate pool, and ten independent sets of 43 were drawn from it uniformly at random (Supplementary Methods). Reporter-assay generalization used the K562 PARM MPRA library (*30, 31*). Reporters ≥ 200 bp were scanned for the 43 motifs (FIMO p < 10⁻⁴). Reporters were split 90/10 by promoter, so no promoter appears in both training and test sets, and performance is the Pearson correlation on the held-out reporters (Fig. 2G).

### Motif, binding-site and protein constraint

Information content per base and CG content of the core PWMs were compared with JASPAR 2022 CORE vertebrates non-redundant (n = 841) and with CIS-BP 2.0 human (n = 6,607 PWMs, comprising directly determined and inferred motifs for the 1,200 human TFs with a motif in the catalogue) by two-sided Mann-Whitney tests. Binding-site conservation is the mean phyloP100way (*33*) score over the exact motif interval of each FIMO hit, reduced to the best hit per gene to avoid inflation from multi-site genes. Core and control motifs were scanned in a single FIMO run per biosample (p < 10⁻⁴), and the pooled comparison uses the 141 distinct control motifs of the ten sets, each once, with every statistic also computed per set. The test uses per-motif medians, not the pseudo-replicated individual sites. CG dependence was removed by a common-slope ANCOVA, and per-biosample consistency was assessed by sign test on the mean over the ten sets. Protein constraint used the gnomAD v2.1.1 constraint table (*34*). The 43 core motifs match 254 non heterodimer TF entries in JASPAR by TOMTOM (a TF can be matched by more than one motif); 250 of these have a gnomAD entry (highest-pLI transcript per gene) and were compared with the 1,338 other human TFs in CIS-BP that have one, on pLI, LOEUF and missense z, with synonymous z as a negative control. Because the 250 TFs derive from 43 motifs, the LOEUF result was re-verified at the motif level against a permutation null (20,000 draws preserving the per-motif TF-count structure). DNA-binding-domain constraint used gnomAD regional missense observed/expected ratios averaged over each TF’s UniProt-annotated DNA-binding domains (DNA_BIND, ZN_FING and DNA-binding DOMAIN features); a TF enters this comparison only when it has such a domain covered by a regional-constraint segment, which leaves 81 core reader TFs and 268 other TFs. Constraint differences across reader families used the modal DBD family per motif (assignment purity 0.93) and a Kruskal–Wallis test over the 12 families with at least five readers.

### Fine-mapped eQTL overlap

Fine-mapped GTEx v8 eQTLs (*35*) were taken unmodified from the benchmark released with Enformer (*10*) and Borzoi (*51*) (SuSiE posterior inclusion probability > 0.9, with count-matched non-causal variants at PIP < 0.01). Motif occurrences are counted inside biosample-specific promoter windows, so each GTEx tissue was hand-paired one-to-one with the EpiMap biosample of the same tissue; 26 GTEx tissues have such a counterpart among the 116 biosamples (table S2; Supplementary Methods), and the remaining tissues were not used. The 16 pairs contributing > 500 causal variants were tested; the excluded ten all show the same direction of effect. Variants were matched to motif-occurrence midpoints within ± 4 bp, the motif interval for 8-bp motifs. The overlap ratio is the number of distinct causal variants inside occurrences divided by occurrences scanned. Core motifs were compared with the mean of the ten control sets by one-sided paired Wilcoxon test.

### ENCODE ChIP-seq occupancy

ENCODE4 narrowPeak files (*36, 52*) for five cell lines were used (1,151 TF–cell-line pairs; 7.5 million peaks). Peaks were reduced to summit-centred ± 150-bp windows within TSS ± 10 kb and scanned with FIMO (p < 10⁻⁴) for the 43 core motifs and the 141 distinct motifs of the ten control sets in a single run. Each motif was summarized by its mean fraction of a TF’s peaks containing a match. Core versus control distributions were compared by one-sided Mann-Whitney test, pooling the control motifs of the ten sets (each once) and also per set; per-cell-line control values are the mean over the ten sets.

### External validation and cross-cell-type specificity

The fixed 43 motifs were applied unchanged to HEK293, a line absent from EpiMap. Because EpiMap-style promoter peaks do not exist for HEK293, promoter windows were built from eleven public peak sets in place of the EpiMap chromatin-state annotation for HEK293 and HEK293T (all GRCh38): ENCODE H3K27ac (36,471 peaks), H3K4me3 (26,738) and H3K4me1 (77,523) for HEK293; ENCODE DNase-seq for HEK293T (85,624); H3K27ac for HEK293T from GEO GSE299613 (34,805); and ENCODE PRO-cap bidirectional, unidirectional and divergent TSS peaks for HEK293 (41,572, 19,436 and 35,923) and HEK293T (37,640, 16,438 and 33,240). The 445,410 summits were pooled, and each gene received the mandatory TSS ± 200 bp window plus up to nine 400-bp windows centred on the summits nearest the TSS within ± 10 kb, at least 100 bp apart, the same construction as for the 116 biosamples (122,248 windows; 6.66 per gene, 12.1% of genes with the TSS window only; Supplementary Methods). The 43 motifs were scanned with FIMO like before, the target was Human Protein Atlas RNA-seq (*53*) (log2(TPM + 1)), and linear, random-forest and CNN-CR models were trained and tested within HEK293 under the internal benchmark protocol: a fixed 90/10 gene split, with the CNN-CR validation fold drawn from the training split. Linear and random-forest hyperparameters were identical to the 116-biosample models; for the CNN-CR the dropout, L2 weight and learning rate were selected on the validation fold only, never on the test genes. The same procedure was applied to twenty synovial-fibroblast samples from patients with rheumatoid or osteoarthritis (*37*). For each patient, promoter windows were rebuilt under the identical construction from the pooled MACS2 summits of that patient’s own ATAC-seq, H3K27ac, H3K4me1 and H3K4me3 peaks (6.74 windows per gene), the 43 motifs were scanned, and the target was that patient’s RNA-seq expression (log2(TPM + 1)); linear and random-forest models used the identical untuned protocol, and CNN-CR learning rate and dropout were selected per patient on the validation fold only (Fig. 4E). The promoter-window-count baseline (fig. S1) was computed for both panels in the same way as for the 116 biosamples: the number of promoter windows each gene received, as the single feature of a linear model fitted on the same 90/10 gene split, scored by Spearman correlation on the held-out genes, alongside the raw correlation between window count and expression over all measured genes.

Cell-type specificity followed Avsec et al.10. Expression was standardized within each biosample. The unit held out is the biosample, not the gene: five-fold cross-validation was partitioned by biosample, and the expression of every gene in a held-out biosample was predicted by the 43-motif CNN-CR models of the training-fold biosamples, each run on the held-out biosample’s own promoter sequence and their predictions averaged, so that no model contributing to a prediction had seen any data from the biosample it predicts. This is the intended test, since it asks whether the 43 motifs predict how a gene’s expression differs between cell types rather than its level in a cell type used for training. Pooled out-of-fold predictions gave one cross-cell-type Pearson correlation per gene across the 116 biosamples (genes with cross-biosample s.d. < 0.01 excluded). Correlations were referenced to a null in which the biosample labels of the predictions are permuted once per round, shared across genes, over 100 rounds.

Genes were stratified into terciles by cross-biosample variance. The chromatin-coverage baseline, the number of active promoter windows a gene receives used as the single feature of a model, was fitted and scored under the same biosample-partitioned cross-validation, so that it differs from the motif model only in which features it is given. The comparison with Enformer was run on Enformer’s own partition of the genome, so that the genes plotted are held out from both models. Enformer’s published human training, validation and test intervals were mapped onto the 18,356 gene promoters, CNN-CR was retrained from scratch on the training genes with the architecture and settings used elsewhere in this work, and both models were then scored on the 1,633 test genes in the 59 biosamples Enformer covers, under the same biosample-held-out cross-validation, standardization and tercile definition as the 116-biosample panel. Each model was scored against the measurement it was trained to predict: Enformer against the measured FANTOM5 CAGE library matched to each biosample, using its published predictions as released (log10-transformed predicted CAGE counts), and CNN-CR and chromatin-coverage baseline against EpiMap RNA-seq. Variance terciles were defined once on measured EpiMap RNA-seq and held fixed for every series, so that all three are scored on identical gene groups.

### Reader assignment at motif sites from ChIP-seq

Readers were assigned in the five ENCODE ChIP-profiled cell lines. Core-motif occurrences (FIMO p < 10-4) were mapped to genomic coordinates within each line’s own promoter windows. A candidate reader (a motif’s JASPAR-matched TF) was scored as binding a gene when one of its top-decile peaks overlapped a copy of that motif in the gene’s promoter. The decile is the strongest 10% of that factor’s own peaks, a within-factor percentile of its own signalValue distribution, so that every peak entering the comparison is a strong site for its own factor and no threshold is carried across experiments (peak counts, signal distributions and the effect of the cutoff for all five lines in fig. S12A-E); this is the only use of peak signal, and no signal value is compared between factors, experiments or cell lines. Occupancy is summarised by breadth of binding: a factor is called the dominant reader of a motif in a cell line when it is bound at the largest number of that motif’s bound genes, provided it has at least 20 genes. Cell lines were then compared two at a time. For each pair of cell lines we used only the genes whose motif copy lies inside the promoter window in both cell lines, and only those candidate readers with ChIP-seq data in both, so that a difference between the two cell lines cannot simply mean that a factor was never measured in one of them. More than one candidate reader can bind the same motif copy at a gene, so each cell line gives a list of the readers bound there rather than a single factor. A gene counts as switched when the two lists are not the same: a reader bound in one cell line but not the other, a reader replaced by a different one, or both. The switch rate of a motif is the percentage of genes that switch, counted over every pair of cell lines (Fig. 5B,C; fig. S12).

### Functional reader switching by Perturb-seq

Knockout effects were the published K562 profiles of Replogle et al.41 (z-scores versus non-targeting controls; 11,258 perturbations) and the HEK293T X-Atlas/Orion atlas (*40*) (18,312 perturbations) which we processed the same way, so that both screens give each gene’s expression as a z-score against non-targeting controls. A TF was a positional reader of a motif at a gene when a FIMO match (p < 10−4) inside any of its called ChIP-seq peaks fell within the gene’s promoter. Only readers that were knocked down effectively (own transcript z < −0.1) in the screen were considered here. The two screens were placed on a common scale by quantile normalization to the average of their quantile functions over the 8,236 genes they share. Within-line dominance (Fig. 5D,E) ranks a motif’s readers in one line by their median absolute effect on the union of the genes they bind, so every reader of a panel is scored on the same genes, and compares the leader with the runner-up by paired Wilcoxon over genes. The size of the lead is the leader over the median of the other readers. Gene order in the curve panels is by general perturbability, the median effect of knockdowns of TFs that do not bind that gene, so no statistic taken from the plotted curves sets the axis. The cross-cell-type comparison (Fig. 5F,G) uses every candidate reader of a core motif that has an effective knockdown in both screens, scored on one fixed gene set: the union of the motif’s ChIP-defined target genes over all its candidate readers in K562 and HEK293, with at least 300 genes. Every reader of a motif is scored on that union in both screens, so the readers differ only as perturbations; thirteen motifs qualified. Switch magnitude is the absolute value of the mean over those genes of log2 of a TF’s absolute K562 effect divided by its absolute HEK293T effect, taken per gene; genes are used where both screens give a finite non-zero magnitude, which excludes 0.02% of observations. Figure 5F draws every reader of the four motifs with the widest spread, Fig. 5G the readers of all thirteen as one lane per motif, and fig. S14 the nine motifs not in Fig. 5F together with the absolute knockout magnitudes behind their ratios. The spread of a motif is the difference between the cross-line ratios of its most K562-leaning and most HEK293T-leaning reader, reported as a fold.

### Statistics and software

Tests are named at point of use. Tests are two-sided unless a directional null is stated. Effect sizes accompany P values, and Bonferroni or Benjamini-Hochberg correction was applied wherever multiple tests were run. Analyses used Python 3.10 with NumPy (*54*), SciPy (*55*), scikit-learn (*56*), pandas (*57*), matplotlib (*58*), seaborn, NetworkX (*59*), pyBigWig and pybedtools. The MEME Suite (*29, 32*) provided TOMTOM and FIMO. BEDTools (*49*), the UCSC Genome Browser (*48*) (phyloP, liftOver) and PyTorch (*50*) were also used.

## Acknowledgments

We are grateful to Hanbei Xiong and Mina Yao for insightful discussions and technical assistance on test run of all the codes.

## Author contributions

WW conceived the project; SS developed the computational models and performed data analysis; SS and WW interpreted the results and wrote the manuscript.

## Data, code, and materials availability

Code is available at https://github.com/Wang-lab-UCSD/CNNCR_WangLab and the processed data archives at Zenodo https://doi.org/10.5281/zenodo.22945154.

