## Additional supplemental files for "A compact motif vocabulary is differentially interpreted across human cell types": data_S2_core_motif_logo_atlas.pdf

Core 0 — 1 JASPAR match: ZNF384

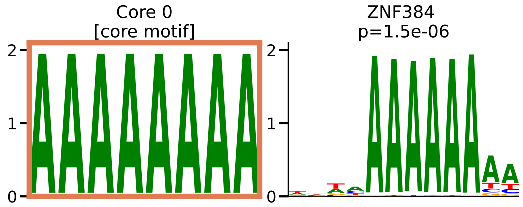

Core 1 — 25 JASPAR matches: ZNF740, VEZF1, KLF12, SP2, KLF15, KLF10, SP1, MAZ, KLF7, KLF14, SP4, ZNF148, RREB1, KLF4, SP5, GLIS3, GLIS1, KLF1, Wt1, ZNF281, ZNF263, ZIC5, ZIC1, GLIS2, KLF9

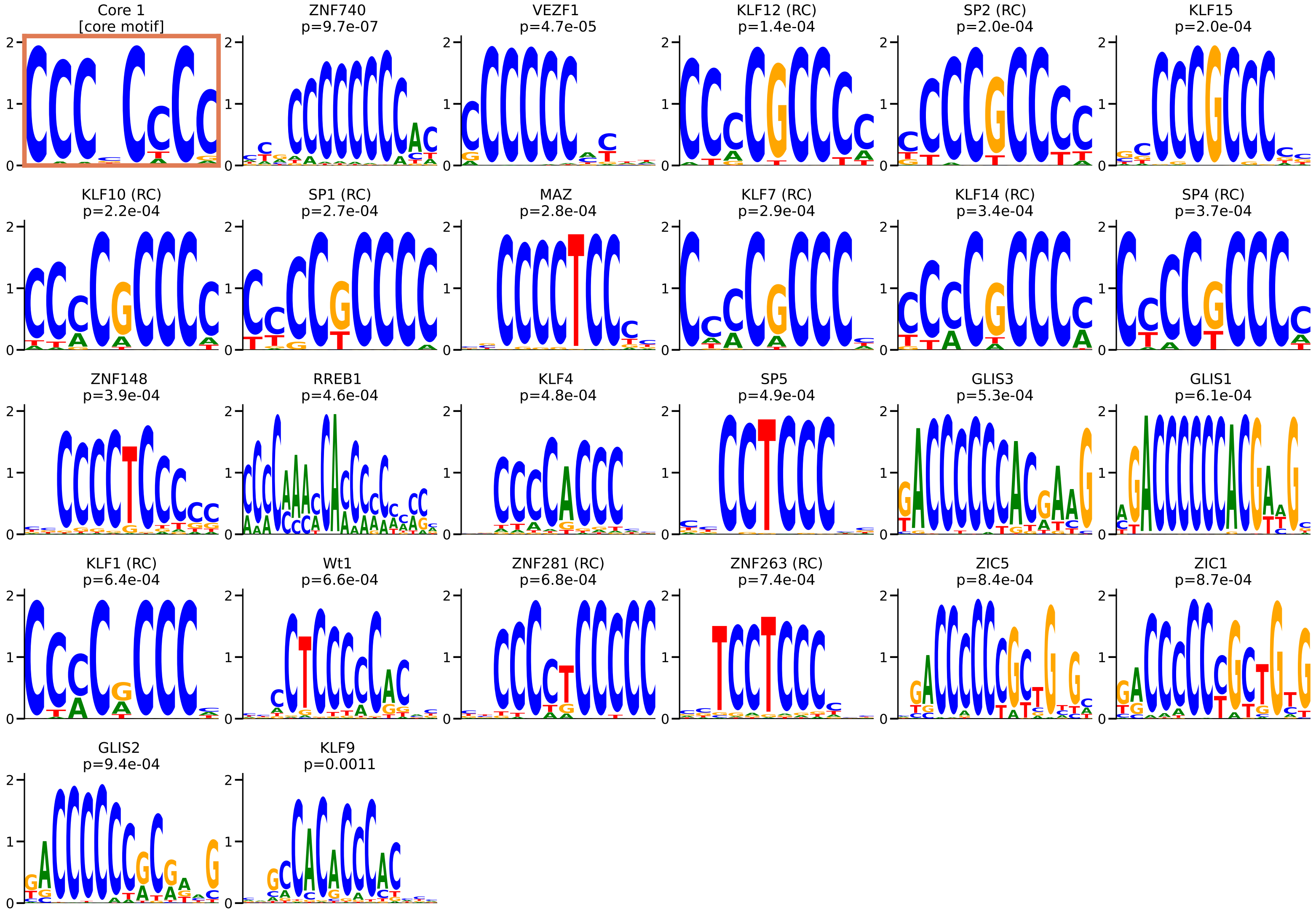

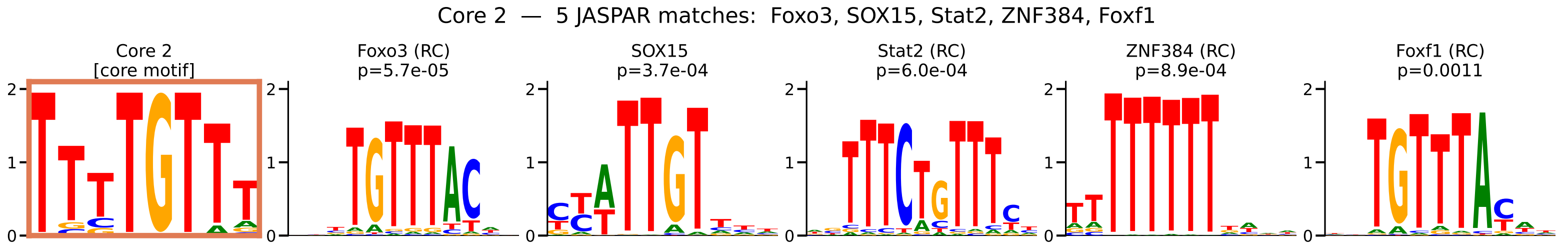
Core 2 — 5 JASPAR matches: Foxo3, SOX15, Stat2, ZNF384, Foxf1
Core 2
[core motif]
Foxo3 (RC)
p=5.7e-05
SOX15
p=3.7e-04
Stat2 (RC)
p=6.0e-04
ZNF384 (RC)
p=8.9e-04
Foxf1 (RC)
p=0.0011

Core 3 — 6 JASPAR matches: Stat5a, Stat5b, Hoxa13, STAT3, ZNF136, ZNF680

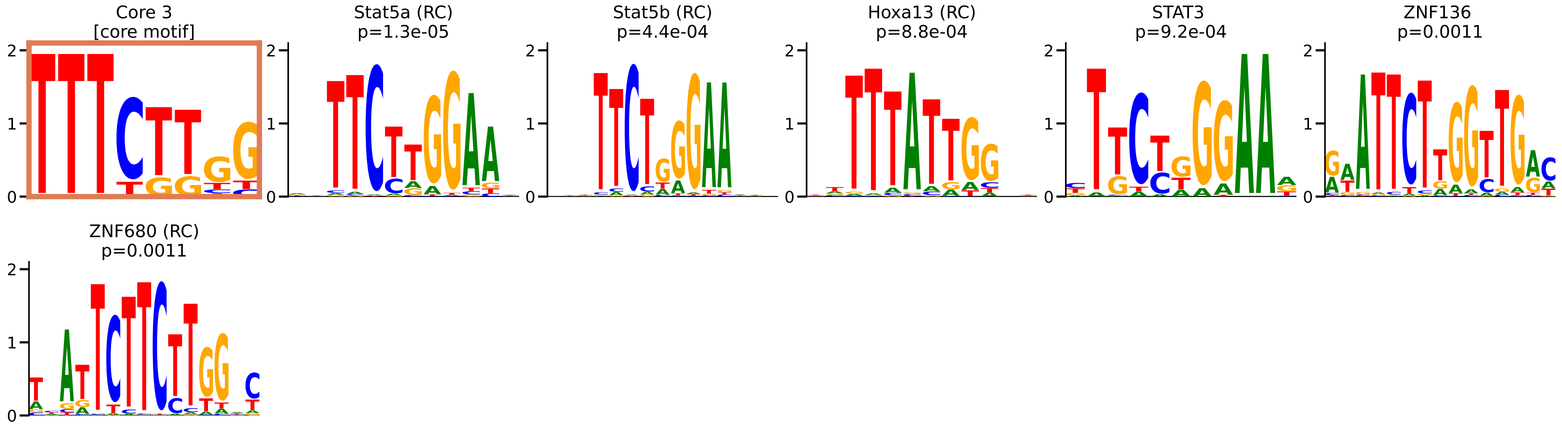

Core 4 — 3 JASPAR matches: Bcl11B, ZNF384, BARX2

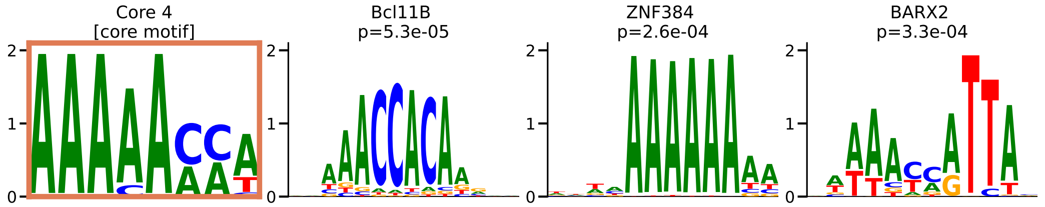

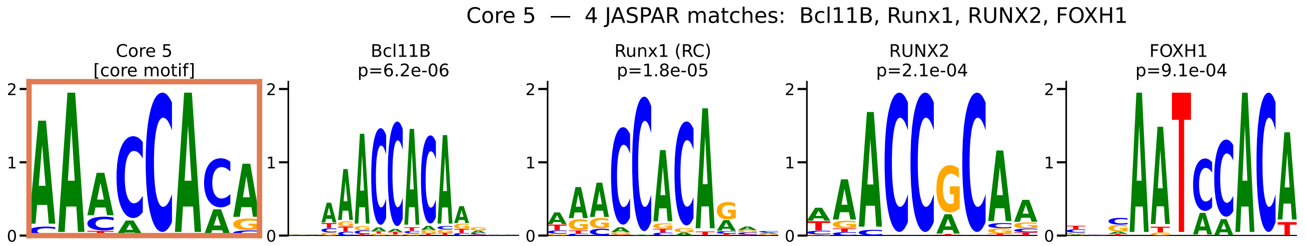
Core 5 — 4 JASPAR matches: Bcl11B, Runx1, RUNX2, FOXH1
Core 5
[core motif]
Bcl11B
p=6.2e-06
Runx1 (RC)
p=1.8e-05
RUNX2
p=2.1e-04
FOXH1
p=9.1e-04

Core 6 — 7 JASPAR matches: HOXB13, Hoxa13, CDX2, HOXD9, Yy1, HOXA10, Hoxd13

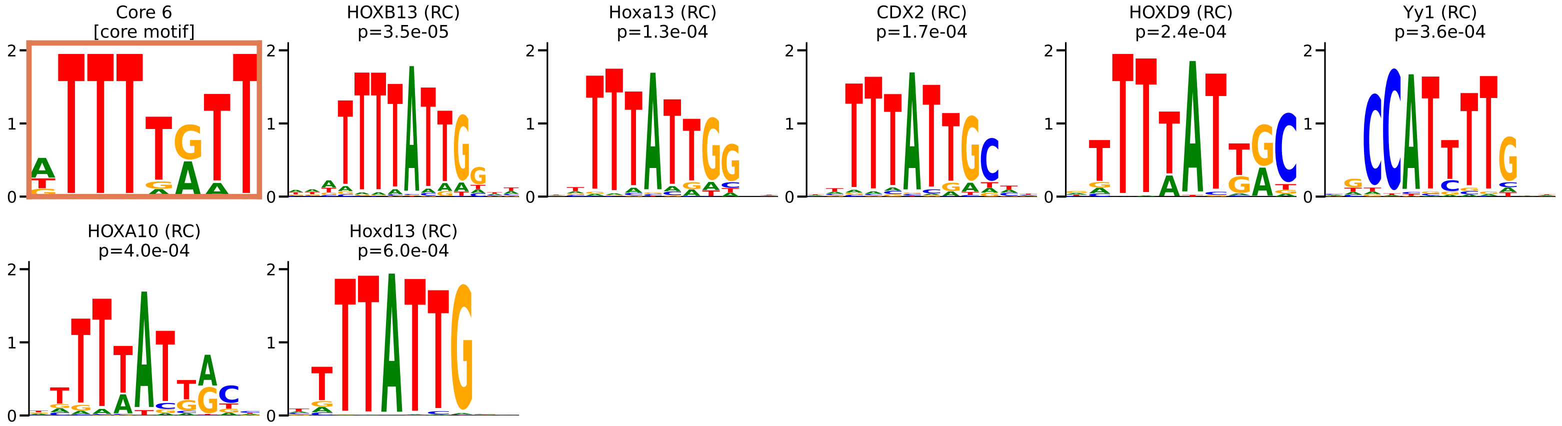

Core 7 — 4 JASPAR matches: PBX2, MEIS2, MEIS1, ONECUT1

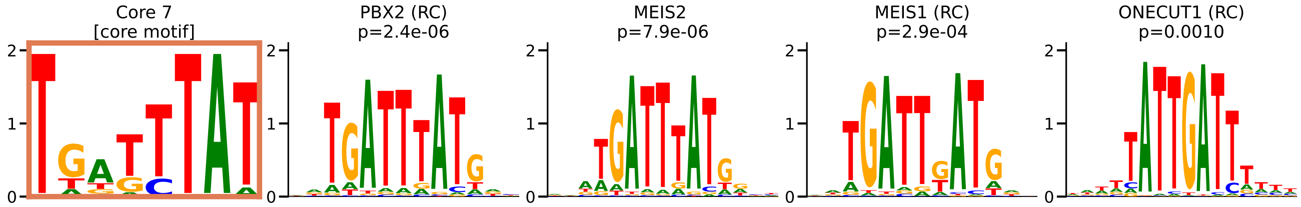

Core 8 — 1 JASPAR match: LIN54

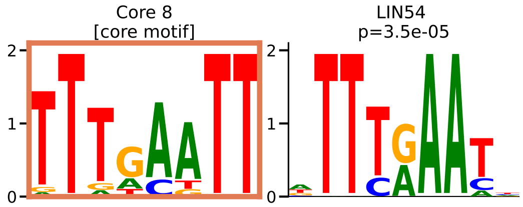

Core 9 — 1 JASPAR match: HINFP

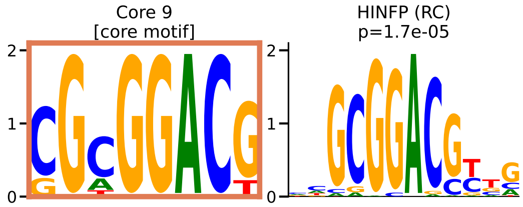

Core 10 — 51 JASPAR matches: MSX1, MSX2, Msx3, ISX, SHOX, RAX2, Shox2, PRRX1, LHX2, LHX9, LBX2, RAX, BSX, PRRX2, UNCX, HESX1, GBX2, EN1, TLX2, ALX3, GSX1, ESX1, LBX1, NKX6-2, EN2, GBX1, MIXL1, VAX2, Nobox, BARX1, HOXA7, ARGFX, PDX1, VSX2, HOXB3, LHX5, Lhx8, NOTO, GSX2, NKX6-1, VENTX, DLX6, Lhx4, Nkx3-1, Dlx3, PAX3, mix-a, EMX2, LHX6, PAX4, ISL2

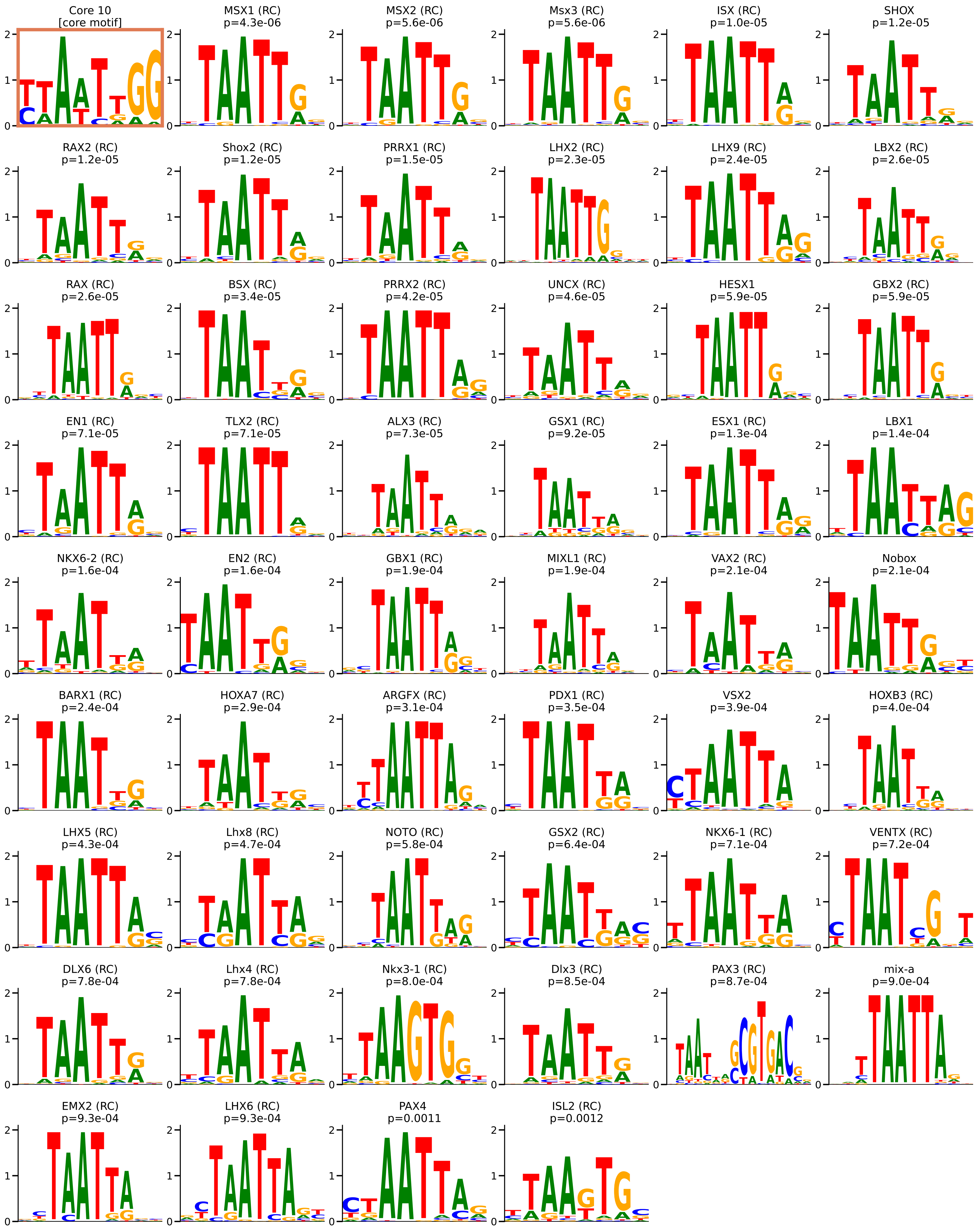

Core 11 — 16 JASPAR matches: CREM, CREB3L4, JUN::JUNB, JDP2, Creb5, FOSB::JUNB, ATF3, JUNB, FOSB::JUN, FOSL1::JUN, FOSL2::JUN, FOS::JUN, ATF7, FOSL2::JUNB, FOSL2::JUND, FOS

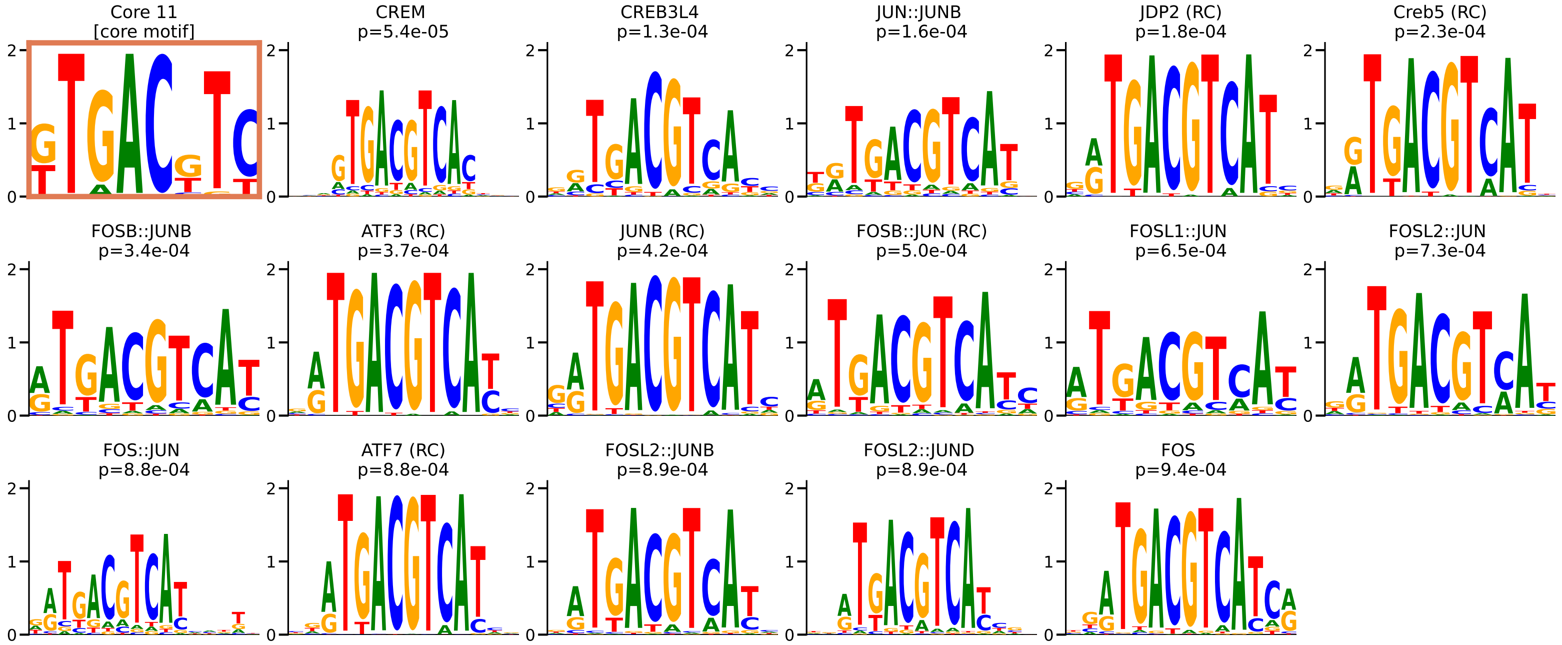

Core 12 — 6 JASPAR matches: ZBTB14, KLF3, HINFP, TCFL5, KLF17, EGR1

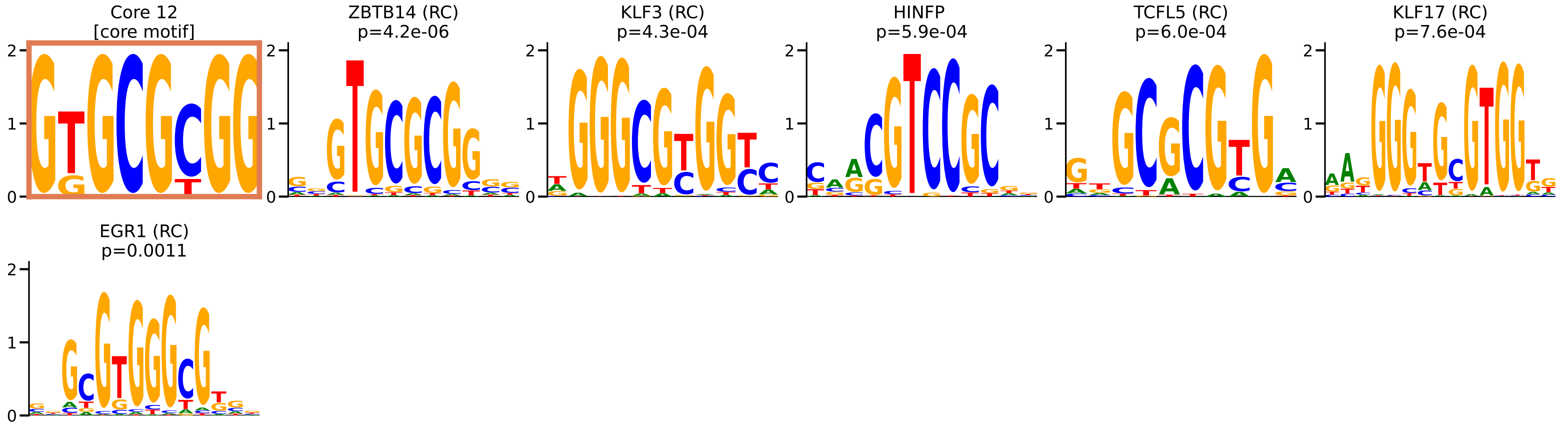

Core 13 — 2 JASPAR matches: ZNF684, ZNF263

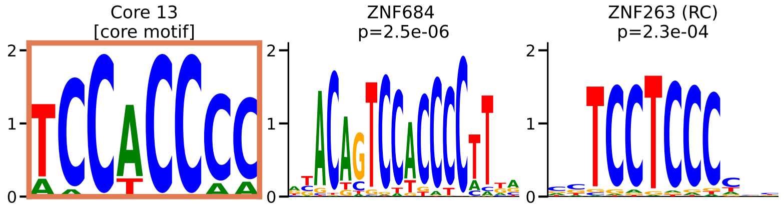

Core 14 — 45 JASPAR matches: ELK1, ETV5, ELK3, ELF4, ELK1::HOXB13, ERF, FEV, ETV3, ETV4, ELK1::HOXA1, ELK1::SREBF2, ZBTB7A, HOXB2::ELK1, ELF2, ELK4, FLI1, ETS1, FLI1::DRGX, ETS2, ERF::NHLH1, ETV6, ERF::HOXB13, ETV5::DRGX, ETV2::DRGX, ETV2, ETV7, HOXD12::ELK1, ETV5::HOXA2, FOXJ2::ELF1, ERF::FOXI1, SPDEF, ETV5::FOXI1, ERF::SREBF2, FOXO1::ELK3, FOXO1::ELK1, ELF1, ETV1, Erg, FOXO1::FLI1, GABPA, Ikzf3, EHF, ELF3, FOXO1::ELF1, ETV2::FOXI1

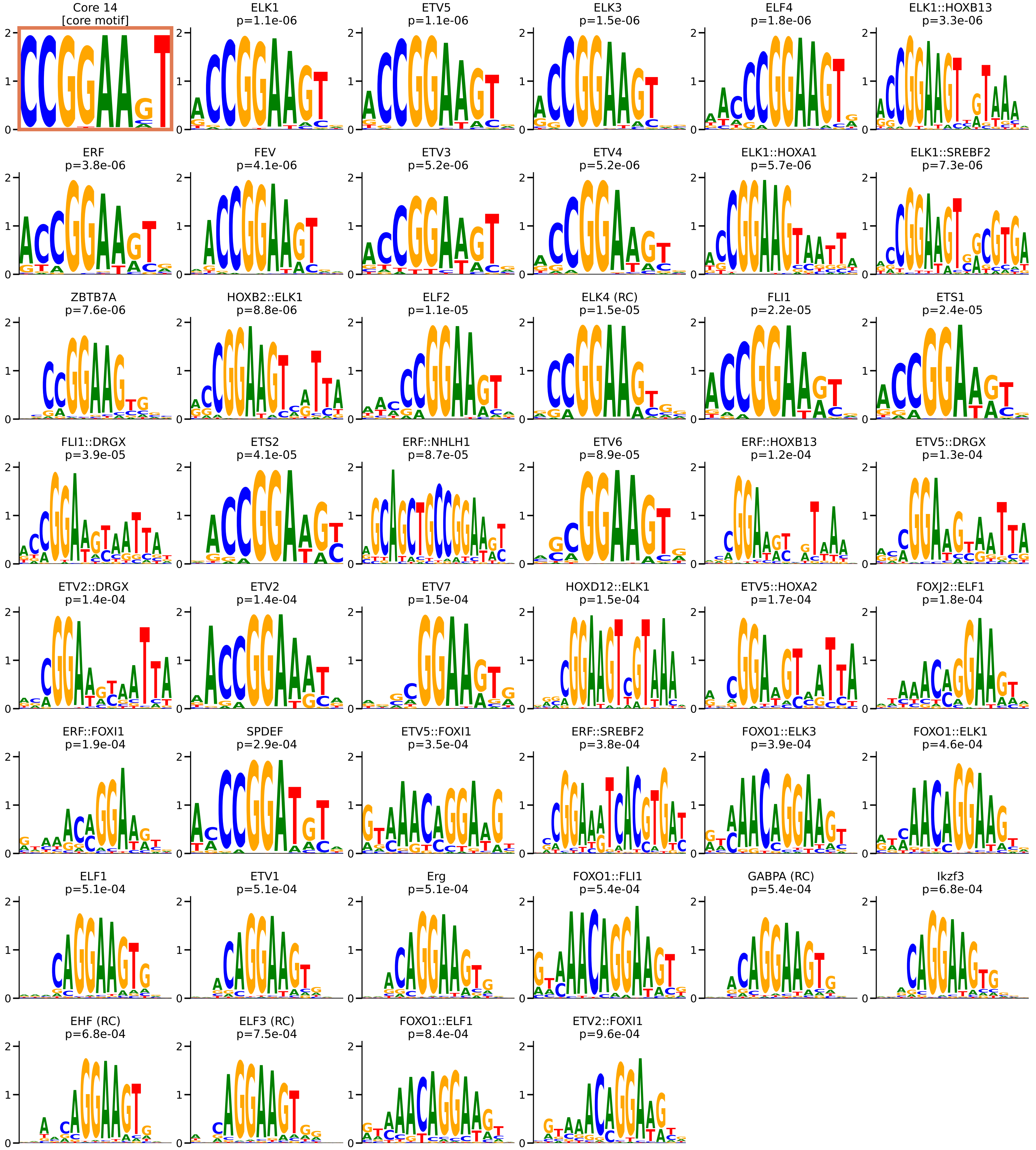

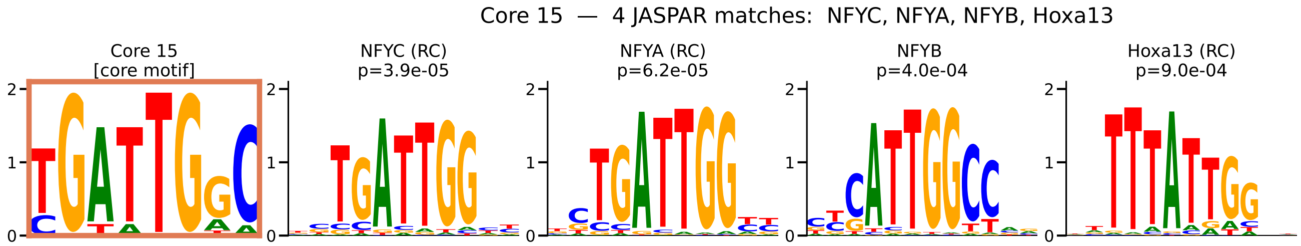
Core 15 — 4 JASPAR matches: NFYC, NFYA, NFYB, Hoxa13
Core 15
[core motif]
NFYC (RC)
p=3.9e-05
NFYA (RC)
p=6.2e-05
NFYB
p=4.0e-04
Hoxa13 (RC)
p=9.0e-04
0
1
2

Core 16  —  4 JASPAR matches:  PRDM1, Spi1, SPIB, Stat2

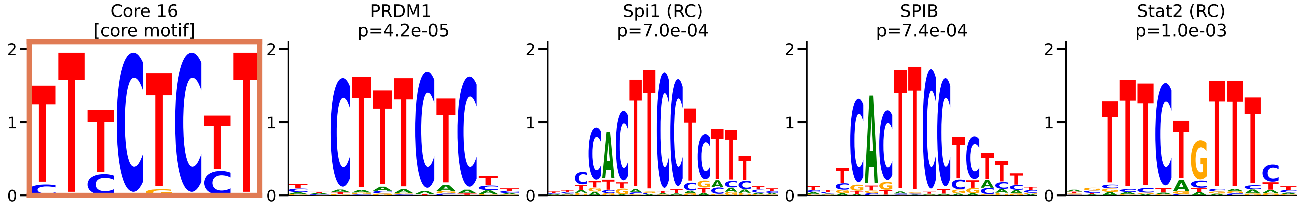

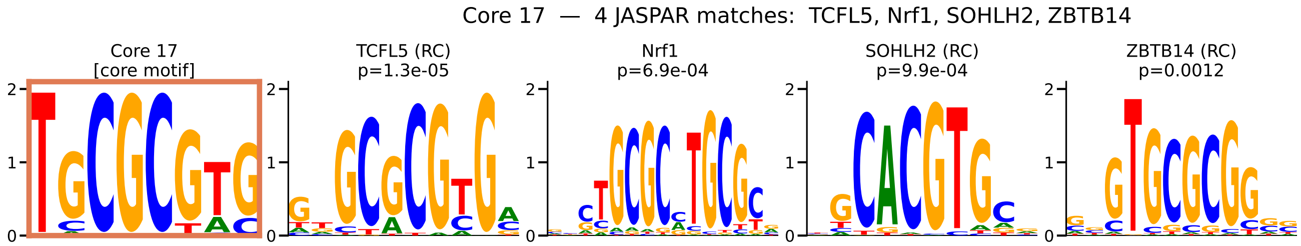
Core 17 — 4 JASPAR matches: TCFL5, Nrf1, SOHLH2, ZBTB14
Core 17
[core motif]
TCFL5 (RC)
p=1.3e-05
Nrf1
p=6.9e-04
SOHLH2 (RC)
p=9.9e-04
ZBTB14 (RC)
p=0.0012

Core 18 — 4 JASPAR matches: SOX10, SOX15, Sox11, SOX4

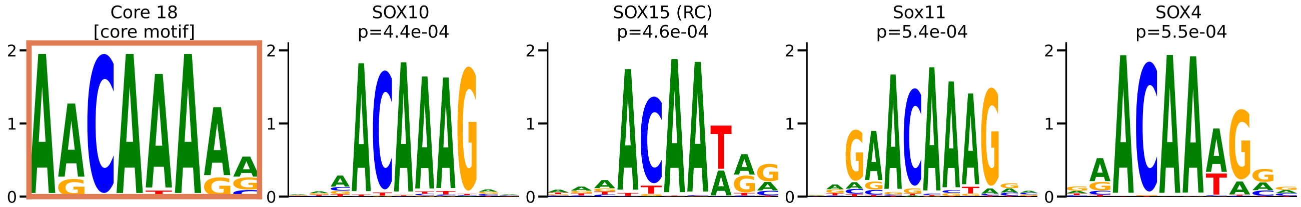

Core 19 — 9 JASPAR matches: ZEB1, TCF3, TCF12, TFAP4::ETV1, ETV2::FIGLA, SNAI1, ETV5::FIGLA, Ptf1A, TFEC

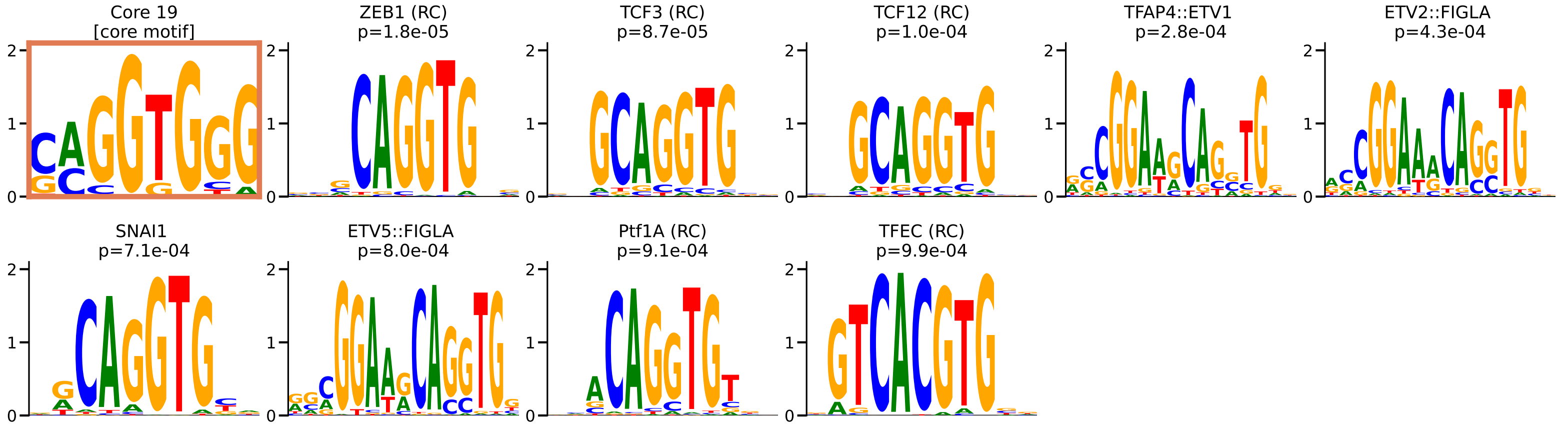

Core 20 — 7 JASPAR matches: E2F3, E2F2, E2F1, E2F4, E2F8, E2F7, ZNF417

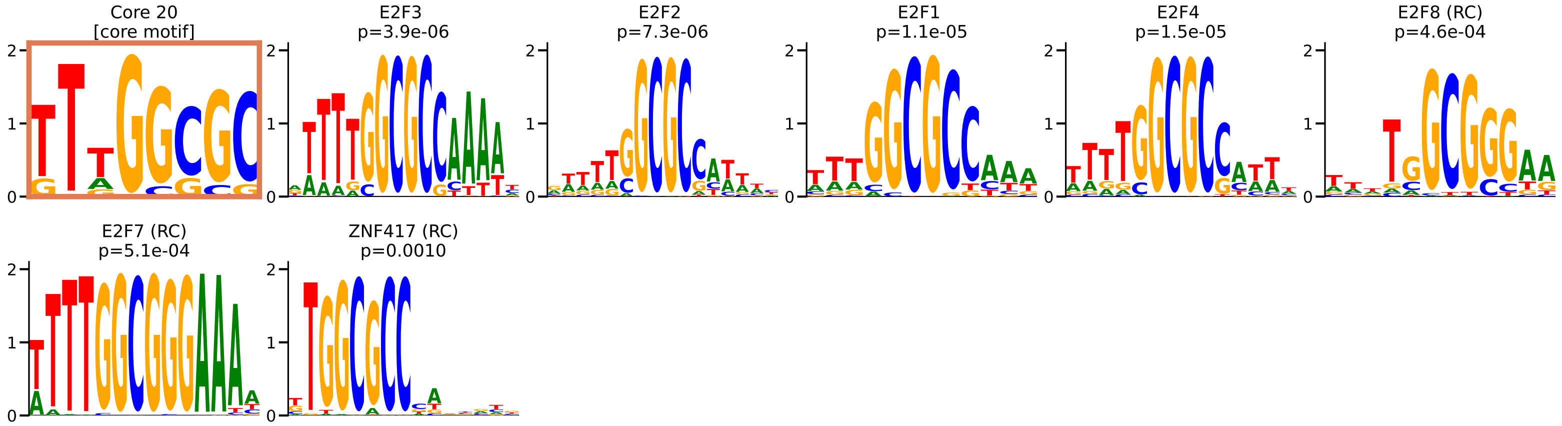

Core 21 — 1 JASPAR match: ZNF331

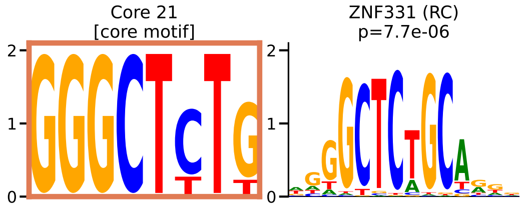

Core 22 — 1 JASPAR match: REST

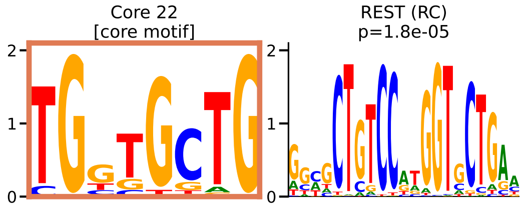

Core 23 — 15 JASPAR matches: TBX15, TBX1, TBX20, TBX18, MGA, TBX4, TBX21, TBX5, TBR1, EOMES, TBX3, Tbx6, TBX2, TBXT, TBX19

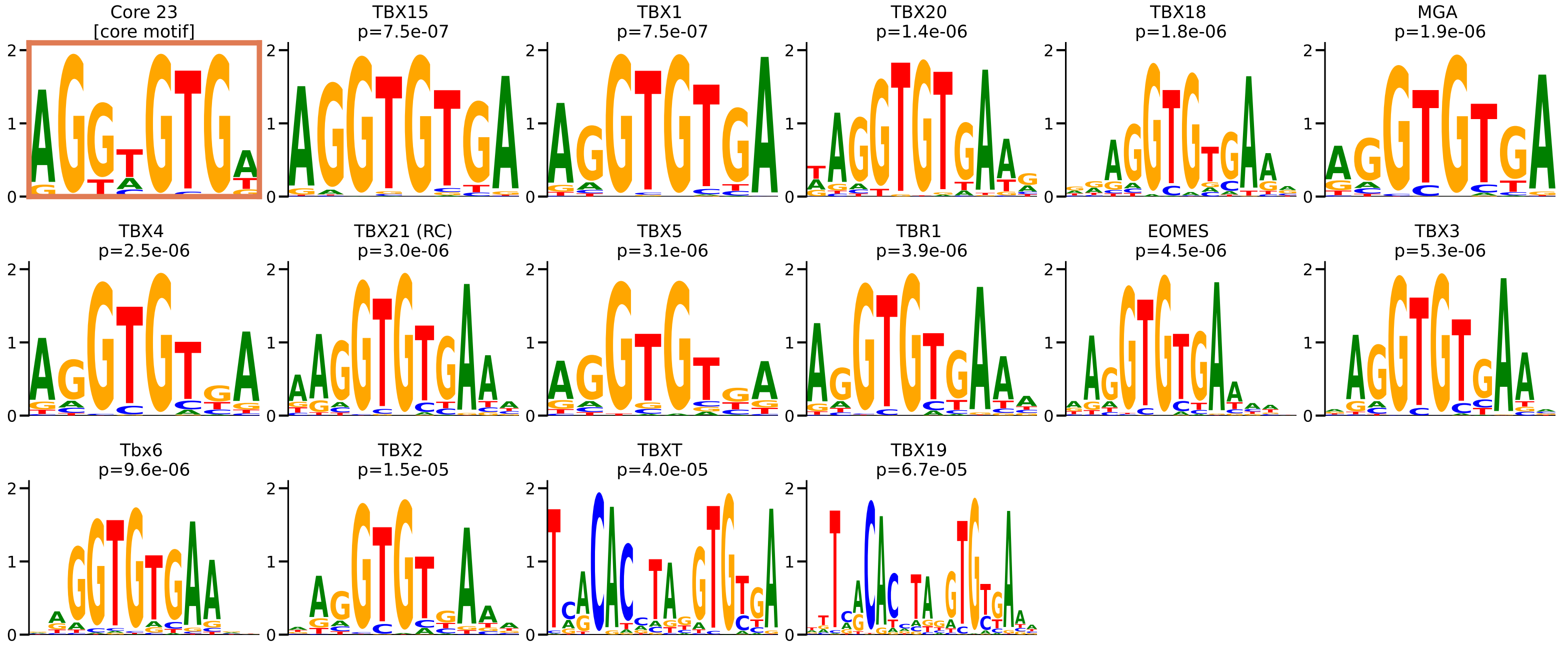

Core 24 — 3 JASPAR matches: MZF1, FOXH1, EBF1

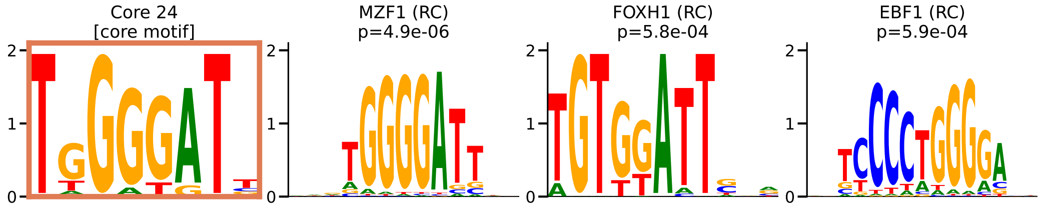

Core 25 — 2 JASPAR matches: ZNF136, Foxo3

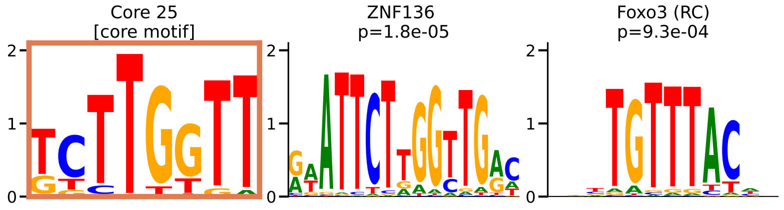

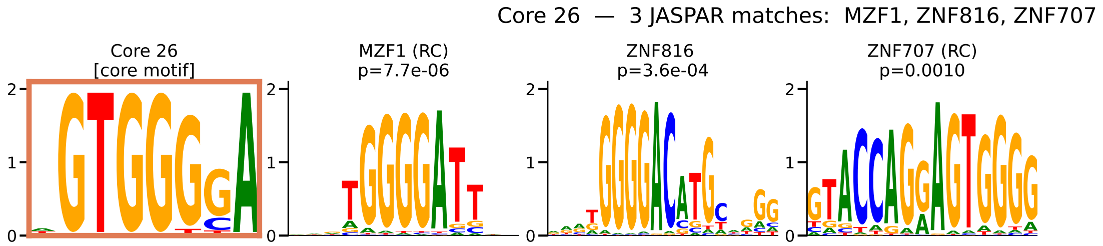
Core 26 — 3 JASPAR matches: MZF1, ZNF816, ZNF707
Core 26
[core motif]
MZF1 (RC)
p=7.7e-06
ZNF816
p=3.6e-04
ZNF707 (RC)
p=0.0010

Core 27 — 2 JASPAR matches: SOX12, SOX14

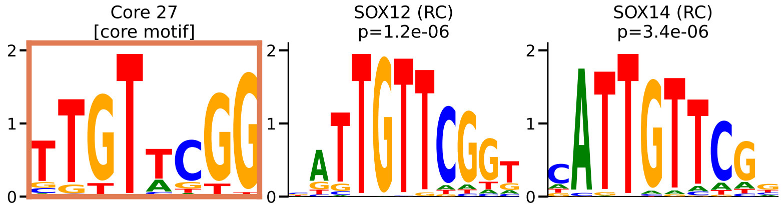

Core 28 — 1 JASPAR match: MSANTD3

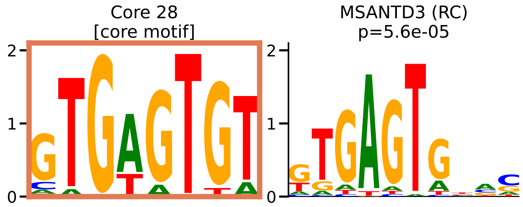

Core 29 — 29 JASPAR matches: ARNT2, Arntl, TFEC, BHLHE41, TFE3, TFEB, USF2, BHLHE40, MITF, MLXIPL, MLX, SREBF2, ERF::SREBF2, Arnt, USF1, Creb3l2, HES6, THRB, ELK1::SREBF2, HES2, CREB3L4, HEY1, HES7, HES1, MNT, MYCN, HEY2, MYC, THRA

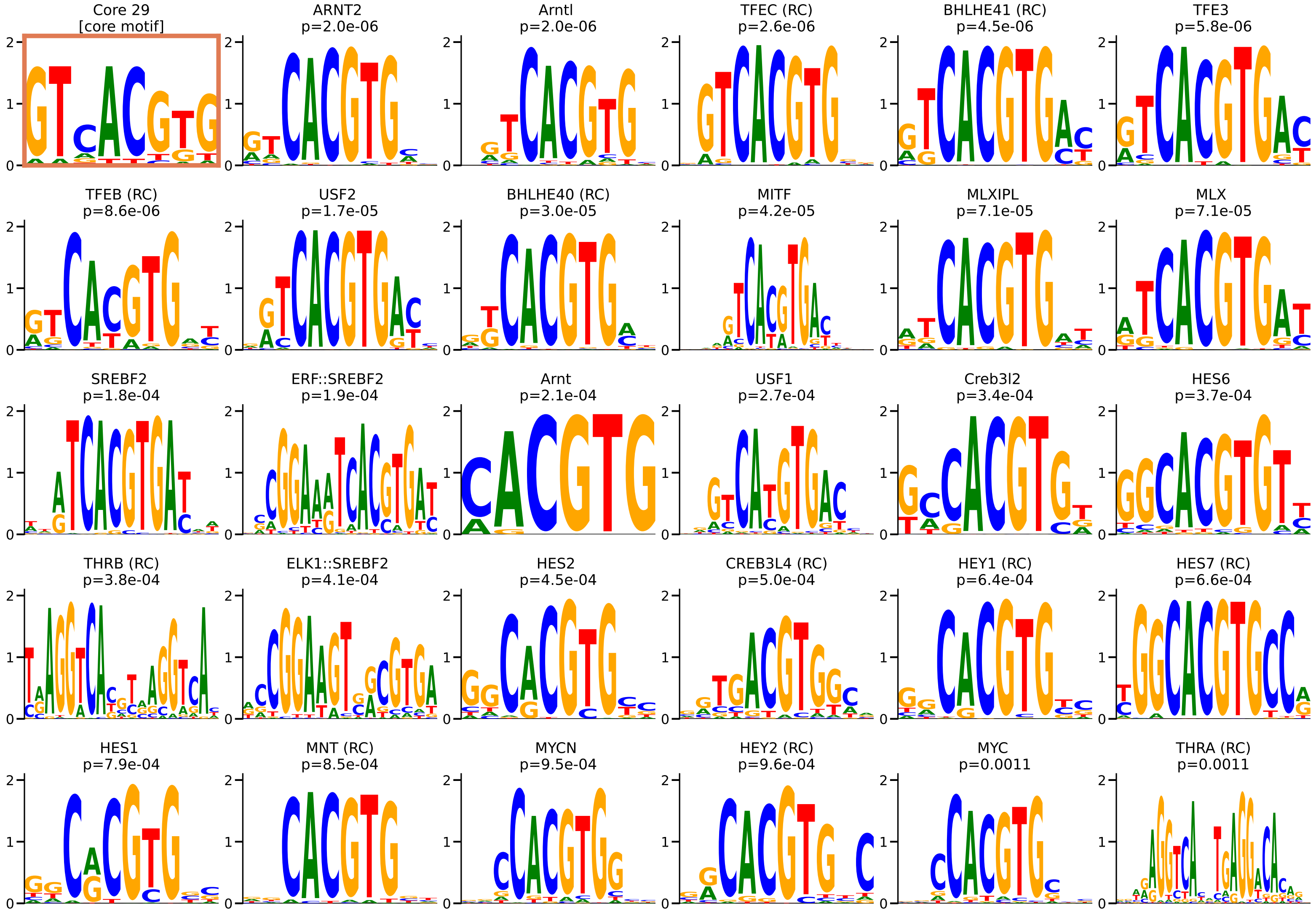

Core 30 — 3 JASPAR matches: ZNF707, KLF9, EGR1

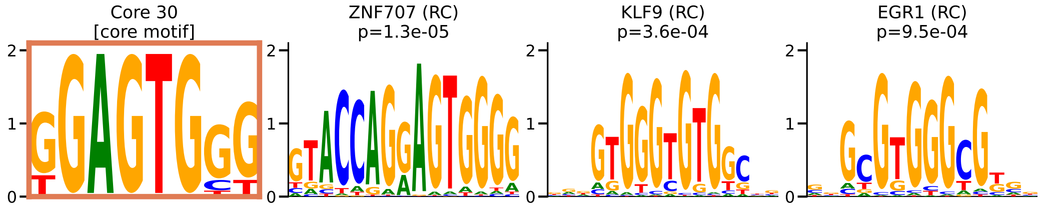

Core 31 — 2 JASPAR matches: SREBF2, SREBF1

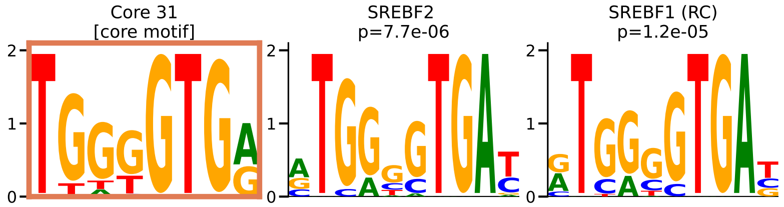

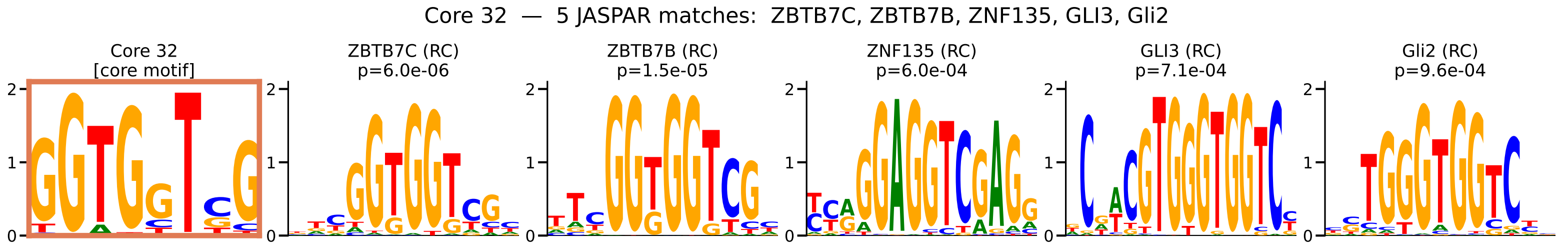

Core 32 — 5 JASPAR matches: ZBTB7C, ZBTB7B, ZNF135, GLI3, Gli2
Core 32
[core motif]
ZBTB7C (RC)
p=6.0e-06
ZBTB7B (RC)
p=1.5e-05
ZNF135 (RC)
p=6.0e-04
GLI3 (RC)
p=7.1e-04
Gli2 (RC)
p=9.6e-04

Core 33 — 1 JASPAR match: ZNF460

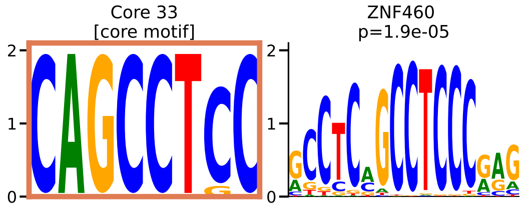

Core 34 — 3 JASPAR matches: ZNF136, ZNF680, Stat5a

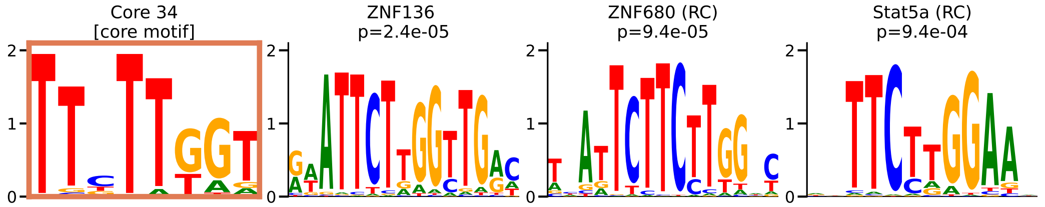

Core 35 — 22 JASPAR matches: NR1H2::RXRA, NR2F2, RARA, Rarb, Rarg, NR2F1, Nr2f6, NR4A1, Nr2f6, TCF7L1, ESRRB, PPARD, RXRG, Rxra, THRB, PPARA::RXRA, Rarb, Hnf1A, RXRB, NR2F1, ESRRA, NR2C2

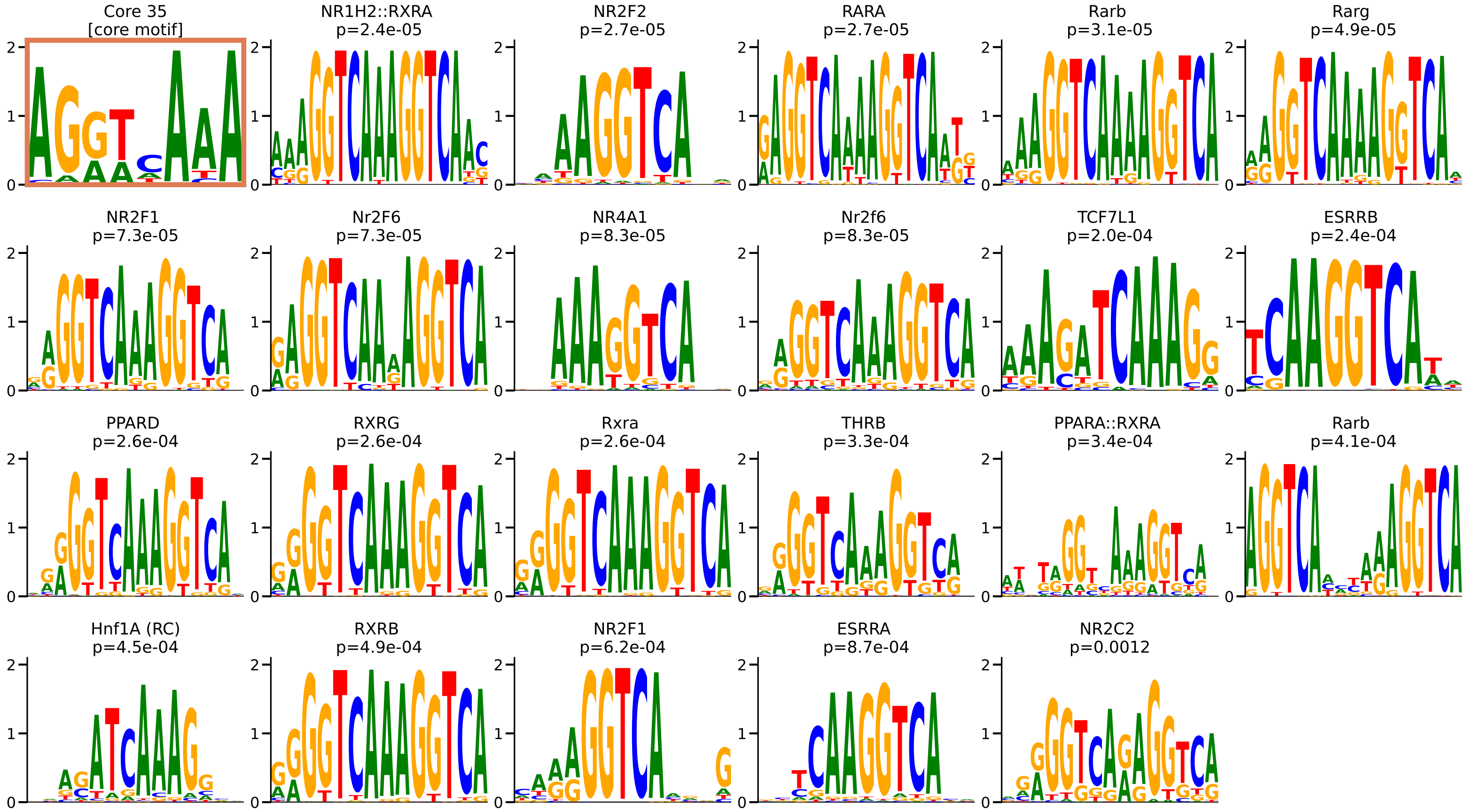

Core 36 — 6 JASPAR matches: E2F1, E2F3, E2F4, E2F2, ZNF417, NFIA

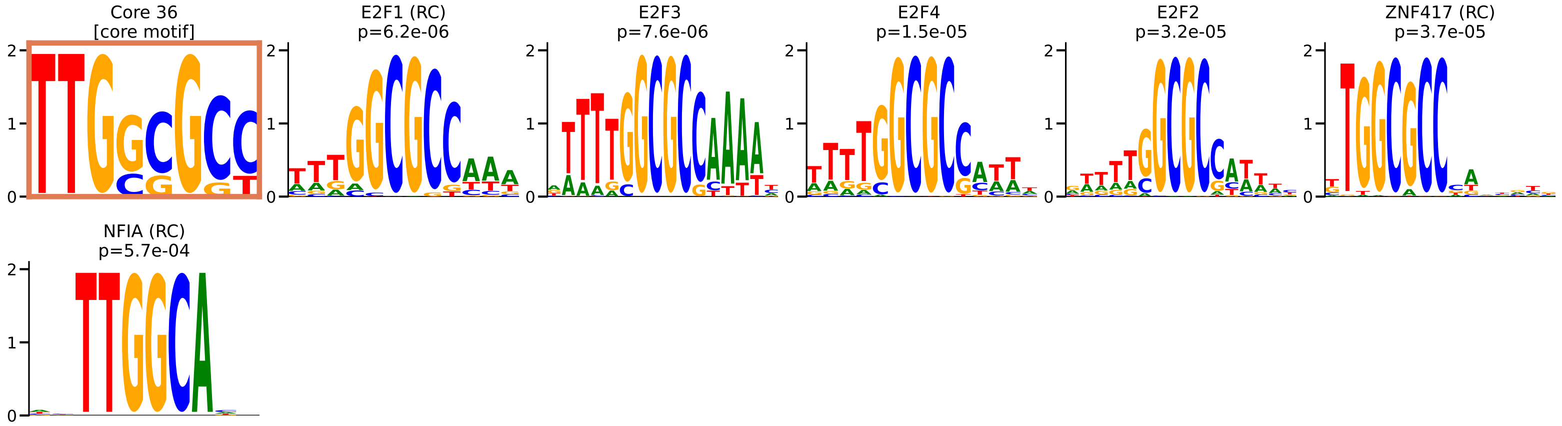

Core 37 — 5 JASPAR matches: PLAGL2, ZNF320, ZNF692, CTCFL, PLAG1

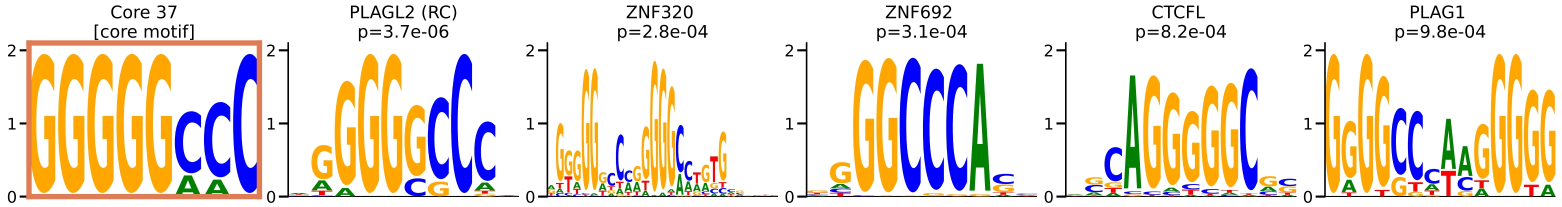

Core 38 — 1 JASPAR match: ZKSCAN1

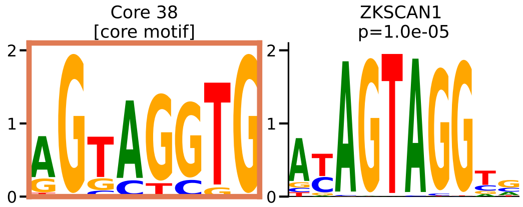

Core 39 — 1 JASPAR match: ZNF768

Core 39
[core motif]

ZNF768
p=5.7e-05

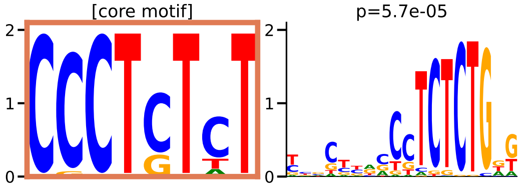

Core 40 — 6 JASPAR matches: GATA6, TRPS1, Mecom, GATA2, GATA4, Gata3

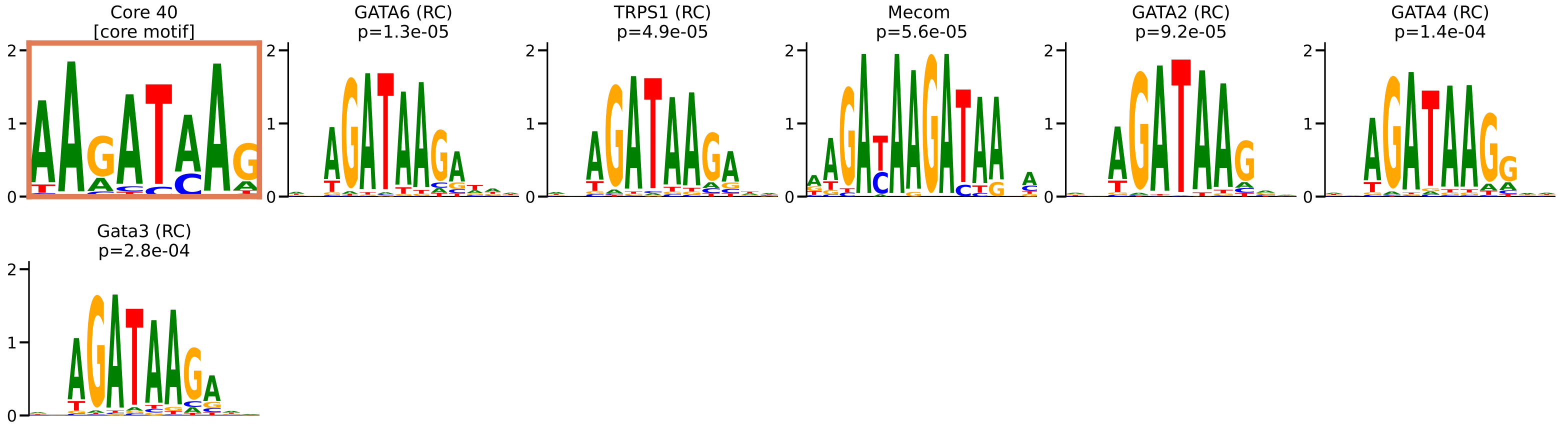

Core 41 — 2 JASPAR matches: ZBTB7B, ZBTB7C

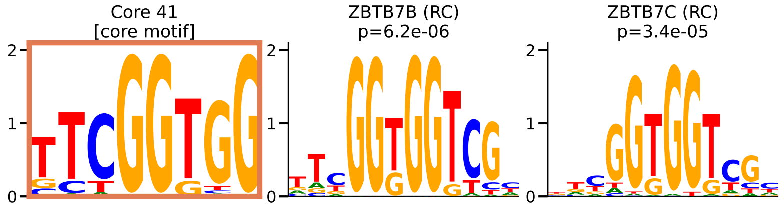

Core 42 — 1 JASPAR match: NFYB

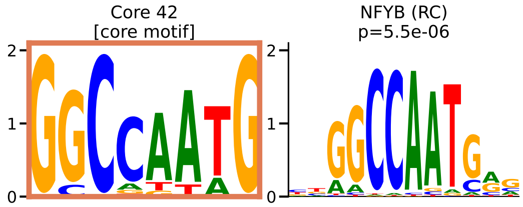
