## Supplemental methods and figures for "A compact motif vocabulary is differentially interpreted across human cell types"

**The PDF file includes:**

Supplementary Methods S1 to S10
Figs. S1 to S14
Captions for tables S1 to S6
Captions for data S1 and S2

**Other Supplementary Materials for this manuscript include the following:**

Tables S1 to S6 (separate machine-readable files)
Data S1 and S2 (separate files)

### Supplementary Methods

Parameter-level descriptions of the procedures that the main Methods summarizes. Software versions and dataset accessions are given at point of use; all code is on GitHub.

S1. Construction of the regulatory sequences

Extraction of promoter sequences

Promoters were defined strictly within the linear window ±10,000 bp around the TSS (the same window applied to genes on the '+' and '−' strands). For each gene, up to ten promoter sequences were assigned. A "fixed" core promoter was defined as ±200 bp around the CAGE-annotated TSS (promoter 0), ensuring a minimum of one promoter per gene regardless of peak annotations. Up to nine additional promoters were selected from EpiMap promoter peak signals within ±10,000 bp of the TSS, requiring that ≥50% of each peak fell within this search boundary. Where more than nine additional peaks were identified, the nine peaks with the widest coordinate spans were retained (promoters 1–9), as peak width positively correlates with regulatory activity. All promoter sequences were extracted from the hg38 reference genome and standardized to exactly 400 bp by symmetric expansion or contraction from the peak center. Genes with fewer than ten promoters received zero-padded placeholder sequences (400 Ns, one-hot encoded as a 400×4 matrix of zeros) to maintain uniform input dimensionality across branches.

Extraction of enhancer sequences

Unlike promoters, which are taken directly from the TSS ±10 kb window, enhancers were not constrained to any fixed window around the gene; they were instead identified through predicted three-dimensional promoter–enhancer contacts. Promoter–enhancer interactions were inferred using EpiTensor, a tensor-decomposition method that predicts 3D genomic interactions from 1D epigenomic signal profiles across cell types (*24*). We retained only strong interactions with an EpiTensor score > 10, yielding 655,133 total promoter–enhancer pairs. For each gene, candidate enhancer regions were identified by intersecting EpiMap promoter peaks with EpiTensor-predicted promoter regions using BEDTools intersect (*49*) (>50% overlap of the promoter peak within the EpiTensor promoter region), and the matched EpiTensor enhancer region was recorded. Active enhancers were then identified by intersecting EpiMap cell-type-specific enhancer peaks with these EpiTensor enhancer regions (>50% overlap), retaining up to ten enhancers per gene selected by peak width. Enhancer sequences were standardized to 400 bp using the same symmetric expansion/contraction procedure as promoters.

Motif-activation vector and thresholds

The first convolutional layer of each branch has 256 filters of width 8 bp and no bias term, applied without padding, so each filter produces 393 activations across a 400-bp sequence. The motif-activation entry for filter m is the arithmetic mean of these 393 raw activations, $a_m = (1/393)\, \Sigma_p W_m \cdot x_{(p..p+7)}$, where $W_m$ is the filter and x the one-hot sequence; no activation function, threshold or binarization is applied at this point, so the entry is a signed continuous value rather than an occurrence count. The contextual-coefficient path applies a ReLU to the same first-layer activations, followed by max-pooling (size 5), three convolutions (64 filters, width 8, same padding) each followed by max-pooling (sizes 2, 3 and 4), flattening (192 features), two dense layers of 256 units with ReLU and dropout 0.15, and a final linear layer giving one coefficient $c_m$ per filter. The branch output is $w_b\, \Sigma_m c_m a_m$ with one trainable scalar $w_b$ per branch, and the prediction for a gene is the sum over its branches; when a gene lacks the sequence for a branch, the one-hot input is all zeros and the branch's contextual coefficients are set to zero. Thresholds enter only in the post hoc interpretation: the PWM of a filter is built from the 1,000 positions with the largest absolute first-layer activation across genes, and the filter's contextual weight is the median of $c_m a_m$ over the genes in which this product exceeds half of its maximum across genes.

Simulated-data positive control

The simulation (fig. S5) was run for five biosamples (A549, GM12878, HeLa, HepG2 and K562) with the same 18,356 genes and the same promoter sequences as the real models. The embedded motif set for a biosample was its own de-replicated promoter motif set (736 to 785 motifs, 3,842 in total), and the target count $C_{g,m}$ was the number of FIMO occurrences of motif m in the real promoter peaks of gene g, so that the simulated genes carry realistic numbers of copies. Each promoter sequence was dinucleotide-shuffled and the gene's copies of each motif were divided as evenly as possible among its promoter sequences (median 4 to 7 non-empty promoters per gene). Each copy was an instance sampled base by base from the motif's PWM and inserted at a uniformly random position that did not overlap a previously inserted copy; a copy was abandoned after ten overlapping draws, which by replay of the placement rule on 1,500 genes per biosample affected 3% (A549) to 15% (HepG2) of the intended copies, so the realised copy number is slightly below C while gene expression y was computed from C. In total 2.0 to 2.8 million copies were placed per biosample into 88,171 to 111,531 promoter sequences, 18 to 31 copies per 400-bp sequence and a median of 101 to 130 per gene; no gene had zero copies. Simulated expression was $y_g = w \Sigma_m C_{g,m} + \varepsilon_g$, with one weight w per biosample (0.020 to 0.026) set so that the signal term had the standard deviation of the biosample's real log2 FPKM, the target centred on the real mean, and$\varepsilon_g \sim N(0, 0.05^2)$; the noise is therefore negligible relative to the signal. Ten promoter-only CNN-CR models per biosample were trained under the production protocol (same hold-out design, optimiser, loss, early stopping and 100-epoch cap; the models stopped after 16 to 100 epochs), and held-out accuracy was the Spearman correlation between the median prediction of the ten models and the simulated expression on the test genes (0.938 to 0.953; median 0.943; Pearson 0.947). First-layer motifs were extracted and de-replicated exactly as for real data, giving 2,014 to 2,233 non-redundant motifs per biosample, and compared with the embedded set by TOMTOM (Pearson distance). TOMTOM reports an E-value that is the match p-value multiplied by the number of motifs in the database being searched, so the same E-value cutoff becomes stricter as that database grows, in direct proportion to its size. The core motifs are matched against JASPAR, which holds 841 motifs, so $E \leq 1$ there means $p \leq 1.2 \times 10^{-3}$. The simulated models yield 2,014 to 2,233 motifs, about 2.5 times as many, so $E \leq 1$ applied here would mean $p \leq 4.5 \times 10^{-4}$ to $5.0 \times 10^{-4}$ and would hold the simulation to a roughly 2.5-fold stricter standard than the JASPAR match it is meant to mirror. Recovery was therefore scored at the common p-value strength, $p \leq 1.2 \times 10^{-3}$, equivalent to $E \leq 2.4$ to $E \leq 2.7$ against these databases. An embedded motif was recovered when at least one extracted PWM matched it at that strength: 687 to 739 per biosample (91.8% to 94.7%; 3,586 of 3,842 pooled, 93.3%). For recovered motifs the best match had a median E-value of 0.28 to 0.35 and median p of about $1.4 \times 10^{-4}$, aligned over all eight positions in 79% to 83% of cases, and about 26% of the recovered motifs reached $E \leq 0.1$.

S2. Within-biosample motif de-replication and annotation

Within-biosample motif de-replication

Redundant motifs were removed in two sequential clustering stages. First, each model's filtered PWMs were clustered by self-comparison using TOMTOM (*32*) (Pearson distance metric; minimum overlap 7 bp; E-value ≤ 0.01; p-value ≤ 0.01; --no-ssc). The resulting pairwise matches were used to build a weighted undirected graph (nodes = motifs; edge weights =

TOMTOM E-values) with NetworkX, and motif groups were identified by Dijkstra shortest-path clustering: motifs joined a group only if all pairwise shortest-path distances were ≤ 0.01. After orienting all members to the same strand, the representative PWM per group was the member closest, in Pearson distance, to the component-wise median of the group. Second, representatives from all 10 models were combined and re-clustered identically, retaining only groups containing representatives from ≥ 2 independent models. This conservative cross-model filter yielded a median of 705 non-redundant promoter motifs and 684 enhancer motifs per biosample, with a maximum of 1,222. The combined motif importance was recorded as the median weight across all instances in the group. These non-redundant sets are the biosample vocabularies counted in Fig. 1E and used as the full vocabularies in the count-based models (Fig. 1H, Fig. 2E). They are not the input of the core-motif derivation: that derivation starts from a smaller, expression-filtered set of representative PWMs built from the same filters (median 73 per biosample, 8,441 pooled; S3).

JASPAR annotation and expression-based filtering

Each representative motif was annotated by TOMTOM against the JASPAR 2022 CORE vertebrates non-redundant database (*25*) (E-value ≤ 0.05; p-value ≤ 0.01). JASPAR TF identifiers were mapped to Ensembl gene IDs via the CIS-BP TF_Information table2. For each motif group, candidate TF annotations were filtered to TFs with biosample expression > 1.66 (log2 FPKM), ensuring annotated TFs are actively expressed; groups sharing JASPAR annotations were merged. A group none of whose members has a JASPAR match to an expressed TF is dropped at this step, so motifs without a catalogue match are not carried into the cross-biosample derivation; they are analysed separately in fig. S9. The retained expressed-TF annotations define the candidate readers used for the constraint analysis (Fig. 3F–H) and the reader analyses (Fig. 5).

S3. Derivation and robustness of the 43 core motif families

Cross-biosample core regulator identification

The cross-biosample derivation does not use the non-redundant vocabularies of S2 (median 705 motifs per biosample); it starts from a smaller, expression-filtered set of representatives built independently from the same first-layer filters. Per biosample, the 2,560 promoter filters of each of the ten models were filtered to total information content > 10 bits and the 750 highest-weighted survivors per model were kept (7,500 PWMs per biosample). These were compared against themselves with TOMTOM (Pearson distance; E-value ≤ 0.01; p-value ≤ 0.01; minimum overlap 7 bp) and against JASPAR 2022 (E-value ≤ 0.05 for annotation), grouped by Dijkstra shortest-path clustering on the self-TOMTOM graph (all pairwise path distances ≤ 0.01), and a group was retained only if at least one of its JASPAR-matched transcription factors is expressed in that biosample (log2 expression > 1.66, from the biosample's own RNA-seq). Groups sharing a JASPAR match were merged, and the member closest to the component-wise median PWM (Pearson distance) was taken as the group's representative. This gave a median of 73 representative promoter PWMs per biosample (range 45–97; the reduction from 7,500 comes mainly from the within-biosample clustering and the expression filter), 8,441 over the 116 biosamples. Cross-biosample clustering. The 8,441 representative promoter PWMs from all 116 biosamples were pooled and subjected to all-vs-all TOMTOM comparison (Pearson distance metric; E-value ≤ 0.5; p-value ≤ 0.01; minimum overlap 7 bp). From the significant matches, an undirected similarity graph was constructed (nodes = motifs; edges between motifs from different biosamples passing the thresholds; edge weight = TOMTOM E-value). Cross-biosample groups were identified by Dijkstra-based network clustering (group_by_network): two motifs were placed in the same group only when all pairwise shortest-path distances among current members were ≤ 0.5. The clustering is therefore determined entirely by graph connectivity in the TOMTOM similarity network; no direct PWM Pearson-distance cutoff gates cluster membership. Per-position Pearson distance was used only (i) to orient all members of a group to a common strand and (ii) to select each group's representative PWM as the member closest to the component-wise median. For each group, the number of distinct contributing biosamples was counted (biosample = the source prefix of each motif name); groups spanning ≥ MIN_CELL_LINES = 25 distinct biosamples were designated core regulators. This yielded 43 core transcriptional regulators, provided in MEME format as data S1.

Robustness of the core-motif cutoff

Two analyses tested whether the >= 25 biosample criterion, and the 43 families it selects, are artefacts of where the threshold was placed (fig. S8).

### S4. Control motif construction

Control motifs were drawn from the JASPAR 2022 CORE vertebrates non-redundant catalogue (841 motifs) in two steps. First, a candidate pool was built by excluding (i) every JASPAR motif with a TOMTOM match to any of the 43 core PWMs (Pearson distance, minimum overlap 7 bp, $E \leq 1$ and $p \leq 0.01$) and (ii) every motif in the same family of a core motif; the pool was re-verified by a second TOMTOM pass against the core motifs and contains 149 motifs. Second, ten independent control sets of 43 motifs were drawn from the pool uniformly at random with fixed seeds; the sets overlap one another by 9–18 motifs and together draw on 141 distinct motifs.

### S5. Binding-site conservation

Conservation at core regulator binding sites was quantified with the 100-vertebrate phyloP track (phyloP100way, hg38) (*33*) from the UCSC Genome Browser. For each FIMO hit in each biosample, the mean phyloP over the exact motif interval was computed with pyBigWig. For each (gene, motif_type) pair, only the maximum phyloP across all hits within the gene's TSS ± 10,000 bp window was retained (best-per-gene), avoiding inflation from genes with many binding sites. The 43 core motifs and the 141 distinct control motifs of the ten control sets were scanned in one FIMO run per biosample (both strands, $p < 10^{-4}$); because FIMO scores each motif independently, the hits of a motif do not depend on which other motifs share the scan, so the ten sets are obtained by subsetting the hits by motif. For the summary comparison in Fig. 3C the test was applied to the per-motif medians (43 core versus the 141 distinct control motifs, each once, and separately 43 versus 43 for each of the ten sets) rather than to the pooled individual sites, because sites contributed by one motif are not independent of one another and pooling them inflates significance without adding information. To test whether the difference merely reflects the higher CG content of core motifs, per-motif median phyloP was regressed on CG content with a common-slope ANCOVA including a motif-set term (Fig. 3D); the CG-adjusted core advantage is reported as that term. The per-biosample consistency shown in Fig. 3E was assessed by a sign test over the 116 biosamples, comparing the core median with the mean over the ten sets of the control median; the number of biosamples in which the core median exceeds every individual set is also reported.

### S6. Fine-mapped eQTL pairing and overlap

Fine-mapped eQTL variants were taken from the GTEx v832 variant-effect benchmark distributed with Enformer (*10*) and Borzoi (*51*) (https://console.cloud.google.com/storage/browser/borzoi-paper/qtl/eqtl/), which supplies, for each of the 49 GTEx tissues, one set of candidate-causal variants already filtered at posterior inclusion probability > 0.9 by SuSiE fine-mapping and one count-matched set of non-causal variants at PIP < 0.01. The complete tissue-level sets were downloaded and used as released; no fine-mapping, thresholding or variant matching was performed in this work.

S7. ENCODE ChIP-seq occupancy

ENCODE4 TF ChIP-seq narrowPeak files (*36*) were obtained for five cell lines (HepG2, K562, GM12878, MCF-7, A549), covering 1,151 unique (TF, cell line) pairs (7,540,192 peaks total; MACS2 calls (*52*)). Peaks within ±10 kb of FANTOM5 TSSs were retained; summit-centered ±150 bp windows were extracted from hg38 with bedtools getfasta (*49*). FIMO (*29*) was applied in a single run to the 43 core PWMs and the 141 distinct control PWMs of the ten control sets against each cell line's peak sequences (match $p < 10^{-4}$). For each (motif, TF, biosample) the fraction_hit = (peaks with ≥ 1 FIMO match) / (total peaks for that TF–biosample pair); each motif was summarized by its mean fraction_hit over all TF– biosample pairs (no occupancy threshold applied), and core versus control distributions were compared by one-sided Mann–Whitney U test, pooling the 141 distinct control motifs (each once) and separately for each of the ten sets; the per-cell-line bars of Fig. 4D show the median over motifs per group, the control value being the mean over the ten sets with its s.d.

S8. External validation on held-out cellular contexts

HEK293 promoter windows. EpiMap promoter peaks do not exist for HEK293, so candidate regulatory summits were pooled from eleven public peak sets for HEK293 and HEK293T (all GRCh38): ENCODE H3K27ac (ENCFF451UZW), H3K4me3 (ENCFF630QMA) and H3K4me1 (ENCFF301UTR) for HEK293; ENCODE DNase-seq for HEK293T (ENCFF285OXK); ENCODE PRO-cap bidirectional, unidirectional and divergent TSS peaks for HEK293 (ENCFF243HMV, ENCFF530NVY, ENCFF387QZO) and HEK293T (ENCFF707DRY, ENCFF624QBA, ENCFF814JPR); and H3K27ac for HEK293T from GEO GSE299613 (GSM9042025). Each peak set was first reduced to one point per peak: for the narrowPeak files (histone marks and DNase-seq) the summit position reported in the file; for the PRO-cap files, which give intervals without a summit, the midpoint of the interval. The points from all eleven sets were then combined into a single list of 445,410 candidate positions. This is a plain union: no set was weighted or given priority, a position was not required to be supported by more than one assay, and identical coordinates reported by several sets were collapsed to one. Windows were then chosen gene by gene. For each gene, the candidate positions within ± 10 kb of its CAGE TSS were sorted by distance from the TSS. The TSS ± 200 bp window was always taken first (bin 0). The sorted positions were then visited in order, and a 400-bp window (position ± 200 bp) was added whenever its centre lay at least 100 bp from the centre of every window already chosen for that gene, until the gene had ten windows or the list was exhausted; windows may therefore overlap, but their centres are at least 100 bp apart. A gene with no candidate position passing this rule kept the TSS window only. This is the same algorithm used for the 116 EpiMap window construction. It yielded 122,248 windows, a mean of 6.66 per gene, with 12.1% of genes having only the TSS window. Windows were scanned on both strands for the 43 core motifs with FIMO at $p < 10^{-4}$, and the count matrix sums occurrences over all windows of a gene. The target was Human Protein Atlas RNA-seq for HEK293 as log2(TPM + 1) over the 18,221 modelled genes with a measurement. Linear (RidgeCV) and random-forest models were trained under the identical untuned protocol and 90/10 gene split (random_state 42) as the internal benchmark, with ten-fold cross-validation within the training split reported alongside; the PWM-initialized CNN-CR used the identical architecture, split and validation fold, restoring the checkpoint with the lowest validation loss, with dropout, per-branch L2 weight and learning rate selected on the validation fold only (the reported model uses dropout 0.3, twice the baseline L2 and learning rate $5 \times 10^{-5}$). Held-out test correlations are referenced to the 116-biosample distributions of the same three model classes (Fig. 4E).

Patient synovial fibroblasts. Twenty fibroblast-like synoviocyte samples from patients with rheumatoid or osteoarthritis (*37*) with ATAC-seq, H3K27ac, H3K4me1 and H3K4me3 ChIP-seq and RNA-seq were used; two further patients lacking RNA-seq were excluded. MACS2 summits of the four assays were pooled per patient (median 497,395 summits) and promoter windows were built under exactly the HEK293 construction above (mean 6.74 windows per gene; 9.1% of genes TSS-window only). The 43 motifs were scanned on both strands at the standard FIMO $p < 10^{-4}$, and the target was the patient's salmon gene TPM as log2(TPM + 1). Linear and random-forest models used the identical untuned protocol; for the CNN-CR, dropout and learning rate were selected per patient on the validation fold only, never on the test genes. Held-out test correlations for all three models are referenced to the same 116-biosample distributions (Fig. 4E). Control-motif models were not run on HEK293 or the patient panel.

S9. Cross-cell-type specificity

To ask whether the core motifs carry cell-type-specific information rather than simply marking genes that are always on, we used the cross-cell-type correlation of Avsec et al.10 Predicted and measured expression were standardized within each biosample (zero mean, unit variance across genes), and each gene was given one Pearson correlation between its predicted and measured values across the 116 biosamples. Biosamples, not genes, are held out: the 116 were split into five folds, and a gene's prediction in biosample b is the average of the 43-motif CNN-CR models trained on the other folds, each reading b's own promoter sequence, so nothing contributing to that prediction has seen a measurement from b. That is the intended design, because the question is whether the motifs predict how a gene's expression changes between cell types, not how genes rank within one, which Fig. 2 already answers. Genes whose measured expression varied by less than 0.01 across biosamples were excluded, as their correlation is undefined. The null permutes the biosample labels of the prediction matrix, one permutation shared by all genes so that each gene keeps its own predicted values but loses its correspondence to the measured biosamples, repeated 100 times. Genes were split into three equal terciles by cross-cell-type variance in standardized expression (Low, Medium, High; $n \approx 6{,}119$ each; Fig. 4F). Marker genes were pre-specified: muscle (MYOD1, ACTA1), erythroid (GATA1, HBB), T cell (CD4, CD8A), liver (ALB, FOXA2), pancreas (INS) and housekeeping controls (GAPDH, SP1). The chromatin-coverage baseline, the number of active promoter windows a gene receives used as a model's only feature, was fitted and scored under the same biosample-held-out scheme, so it differs from the motif model only in the features it is given. Enformer predictions were available for 59 of the 116 biosamples over 18,346 shared genes.

Enformer was trained on genomic intervals rather than on a gene split, so most of the 18,346 shared genes lie inside sequences it was fitted on, and scored there it is partly recalling its training data. The comparison therefore uses Enformer's own split. Each gene was assigned to the split of the published human interval containing its transcription start site, giving 14,626 training, 1,511 validation and 1,831 test genes, of which 1,633 pass the coverage and variance filters and are the genes plotted. CNN-CR was retrained from scratch on the training genes alone, with the same architecture and settings used elsewhere in this work, so the plotted genes are new to both models; CNN-CR is in addition scored with the biosample held out, whereas the 59 cell lines are part of Enformer's training data, so the standard applied to CNN-CR is the stricter of the two. Both were scored on those genes in the 59 biosamples under the same standardization and terciles, with Enformer's predicted CAGE counts log10(x + 1) transformed and its published predictions used as released, since rescaling them onto the measured distribution would feed measured information into the predictions. Each model is scored against the measurement it was trained to predict: Enformer against the measured FANTOM5 CAGE library its matched output track was fitted to, CNN-CR and the chromatin-coverage baseline against EpiMap RNA-seq. Terciles were fixed once on measured EpiMap RNA-seq for every series, because terciles defined on CAGE would regroup the genes. Enformer is applied zero-shot from its published weights whereas CNN-CR is fitted to EpiMap expression, so the comparison measures cross-biosample resolution rather than two equally trained models.

### S10. Reader assignment and functional reader switching

#### Reader assignment at individual motif sites

We assigned readers at the level of individual motif sites in five ENCODE ChIP-profiled cell lines (K562, HepG2, A549, GM12878, HEK293). Core-motif occurrences were taken from the FIMO scans ($p < 10^{-4}$) already computed on each gene's promoter windows for the linear/CNN models, and each occurrence was mapped back to its genomic coordinate (accounting for window strand). Each cell line was scored on the motif copies inside its own promoter windows, which differ between lines because the chromatin annotation that defines the windows is biosample-specific (246,571 to 317,970 gene-motif loci per line). Pairwise comparisons are made on the motif copies that fall inside the promoter window in both cell lines of the pair, so each comparison reads the same DNA in both cell lines. For ChIP-seq we used each factor's top-intensity peaks — the top decile of a factor's own signalValue distribution (a within-factor percentile, since absolute signalValue varies ~10–15-fold between experiments), keeping the full peak interval and requiring overlap with a TSS ± 10 kb promoter (peak-selection summary for all five lines in fig. S12). A candidate reader factor (a motif's JASPAR-annotated TF) was scored as binding a motif site when one of its top-intensity peaks overlapped that copy; where a motif occurred several times at a gene, the factor's overlapping peak with the highest signalValue was used. Binding is binary from this point on. Peak signal is used only in the within-factor top-decile filter above, and no signalValue is compared between factors, experiments or cell lines, because absolute ChIP-seq intensity depends on the antibody or epitope tag, the crosslinking and the sequencing depth of each experiment and is therefore not comparable across them. Occupancy is summarised by breadth: for a motif in a cell line, each candidate reader is scored by the number of that motif's bound genes at which it binds, and the dominant reader and the switch rate are defined from these counts below. Fig. 5B encodes the same count twice, as the fraction of the motif's bound genes at which a reader binds (bubble size) and as that reader's share of all candidate-reader binding at the motif in that line (shade); the two have the same numerator and differ only in the denominator, so they rank the readers of a line identically and differ only between lines, where the share also reflects how many candidate readers that line has profiled.

Two quantities are reported per motif: which factor reads it most widely in a cell line, and how far the readers present at a gene differ between cell lines. For a motif in one cell line, every candidate reader is scored by the number of that motif's bound genes at which it has a qualifying peak, and the factor with the largest count is the dominant reader of that motif in that line, provided it has at least 20 target genes; a line with no candidate reaching 20 genes contributes no dominant reader for that motif. The dots in Fig. 5C are the number of distinct factors that hold this position across the five lines, so one dot means the same factor reads the motif most widely in every line in which it can be called. Switching is assessed between cell lines two at a time. For each pair we use only the genes whose motif copy falls inside the promoter window in both cell lines, and only those candidate readers with ChIP-seq data in both, so that a reader difference cannot simply mean that a factor was never measured in one of them. For a gene carrying the motif and bound by at least one of those readers in both lines, we take the set of readers bound at it in each line and count the gene as switched when the two sets are not identical; a reader gained or lost counts as well as one exchanged for another. The switch rate of a motif is the number of switched genes divided by the number of genes scored, pooled over all cell-line pairs, and Fig. 5C reports it for the 12 motifs with at least 20 scored genes. The two quantities are independent: a motif may keep one dominant reader in every line (one dot) while the readers accompanying it change at most genes (a long bar), as for motifs 29 and 20, so the bar measures turnover in the reader set and the dots measure turnover in the leading factor. The four motifs drawn in Fig. 5B are those with more than one distinct dominant reader and the highest switch rate, excluding motif 15, whose two readers NFYA and NFYB are subunits of the same NF-Y complex rather than competing paralogues; reader spectra for the switching motifs not drawn in Fig. 5B are in fig. S12F. This gene-level definition is distinct from the aggregate occupancy metric used for the ChIP-seq enrichment of Fig. 4D; the functional Perturb-seq analysis below uses the same positional rule, a FIMO motif instance inside one of the factor's ChIP-seq peaks within a promoter, but over all of a factor's peaks rather than its top decile.

<u>Functional reader switching by genome-wide Perturb-seq</u>

Reader function was tested in the two cell types that have both ENCODE TF ChIP-seq and a genome-wide Perturb-seq screen, K562 and HEK293/HEK293T. Binding was taken from ENCODE4 TF ChIP-seq narrowPeak calls for K562 and HEK293 (HEK293 rather than HEK293T because only about 17 factors have ChIP-seq in HEK293T, whereas HEK293T is the line with a genome-wide screen; the analysis therefore assumes that a factor binding a motif copy in HEK293 also reads it in HEK293T). For this functional analysis all of a factor's peaks were used. A TF was a positional reader of core motif M at gene g when a FIMO match to M ($p < 10^{-4}$) lay inside one of its summit-centred (± 150 bp) peaks within g's promoter (TSS ± 10 kb). The genes so defined are the reader's ChIP-defined target genes.

<u>Perturb-seq data and effective knockdown</u>

Knockout effects for K562 were the published normalized profiles of Replogle et al.41 (expression as z-scores relative to non-targeting controls; 11,258 perturbations); for HEK293T, the X-Atlas/Orion genome-wide Perturb-seq atlas (*40*) (18,312 perturbations), which we converted to the same representation as the Replogle profiles: for each perturbation and gene, the difference between the mean expression in the perturbed cells and in the non-targeting-control cells, divided by the standard deviation of that gene across the non-targeting single cells (a z-score against the non-targeting controls). A perturbation was used only where its knockdown was effective in the screen being analysed, defined as the factor's own transcript decreasing on its knockout (self $z < -0.1$); this excludes factors not expressed in that line, whose guides yield only sampling noise.

<u>Quantile normalization</u>

The two screens differ in the shape of their genome-wide effect distributions (median $|z|$ about 0.030 in HEK293T against 0.058 in K562), reflecting the different platforms and cell types. To make effects directly comparable, both distributions were quantile-normalized to a common reference defined as the average of the two screens' quantile functions, computed on the 8,236 genes measured in both, so that their marginal distributions become identical. The mapping is rank-based, so it is robust to the small number of extreme HEK293T values arising from genes with near-zero control variance; genes with exactly zero control variance in HEK293T (about 5% of that matrix, largely unexpressed genes) were treated as non-finite and excluded. All effects below are quantile-normalized absolute knockout effects.

Within-line dominance (Fig. 5D,E; fig. S13)

Within one cell line every reader of a motif is perturbed in the same screen and read out against the same reference, so readers can be compared directly. For each core motif and cell line, the readers are the positional readers of that motif with an effective knockdown in that screen. The panel's gene set is the union of the readers' ChIP-defined target genes, so every reader of the panel is scored on the same genes, including genes it does not itself bind; a panel required at least two readers and at least 100 genes, which left 19 motif and cell line panels. Readers were ranked by their median absolute effect over the panel's genes. The lead is the leader's median divided by the median of the other readers' medians, and separately by the runner-up's median; a 95% interval for each reader's median was obtained from 400 bootstrap resamples of the genes. The leader and the runner-up were compared by a paired Wilcoxon signed-rank test over genes, and the share of genes at which the leader has the larger effect is drawn as the strip above each curve panel. In the curve panels genes are ordered by how responsive each gene is to perturbation in general: for each gene we took the absolute knockout effects of all ChIP-profiled TFs that do not bind that gene and used their median as the gene's general perturbability. Genes are sorted by this value, from most to least responsive, so the ordering is fixed before any reader curve is drawn and never depends on the readers being compared. No threshold on effect size is applied anywhere in this analysis; every reader with any target gene at the motif is included.

Cross-cell-type switch (Fig. 5F,G; fig. S14)

The cross-cell-type comparison uses core motifs with at least 300 ChIP-defined target genes and at least two candidate readers with an effective knockdown in both Perturb-seq screens. Every such reader of a motif is included. The motif's gene set is the union of its candidate readers' ChIP-defined target genes, each reader's targets taken from the line in which it was ChIP-profiled, and every reader is scored on that union in both screens, so the readers differ only as perturbations. Thirteen motifs qualified. For every reader and every gene in that set, the absolute effect in K562 was divided by the absolute effect in HEK293T and the ratio taken in log2; the mean of this per-gene log2 ratio over the gene set gives the reader's cross-line ratio, and its absolute value is the reader's switch magnitude. The mean is taken in log2 space because the arithmetic mean of raw ratios is not symmetric under exchanging the two screens and is dominated by small denominators. A gene is used where both screens give a finite, non-zero magnitude, which excludes 0.02% of observations; no floor, pseudocount or cutoff is applied. In Fig. 5F every reader of a motif is drawn along the gene axis as its own curve, coloured by its median ratio from violet (larger in HEK293T) through grey (no preference) to teal (larger in K562), with the colour saturating at a 2.8-fold median ratio and a running median over 2.5% of the genes applied for display only; the marginal at the right names each reader at its median ratio. A motif whose readers fall on both sides of zero has readers that favour different cell lines; a motif whose readers all lie on one side differs between the lines in overall knockout size rather than in reader identity. Fig. 5G places all thirteen motifs on one axis, one lane per motif, each reader as a dot at its cross-line ratio, so the spread of a lane is the difference between the motif's most K562-leaning and most HEK293T-leaning reader. fig. S14 draws the nine motifs not shown in Fig. 5F, each as its ratio panel beside the absolute knockout magnitudes the ratio is formed from, so that a ratio driven by one screen alone can be distinguished from one in which both screens move. No comparison with other TFs is made.

**Fig. S1.**

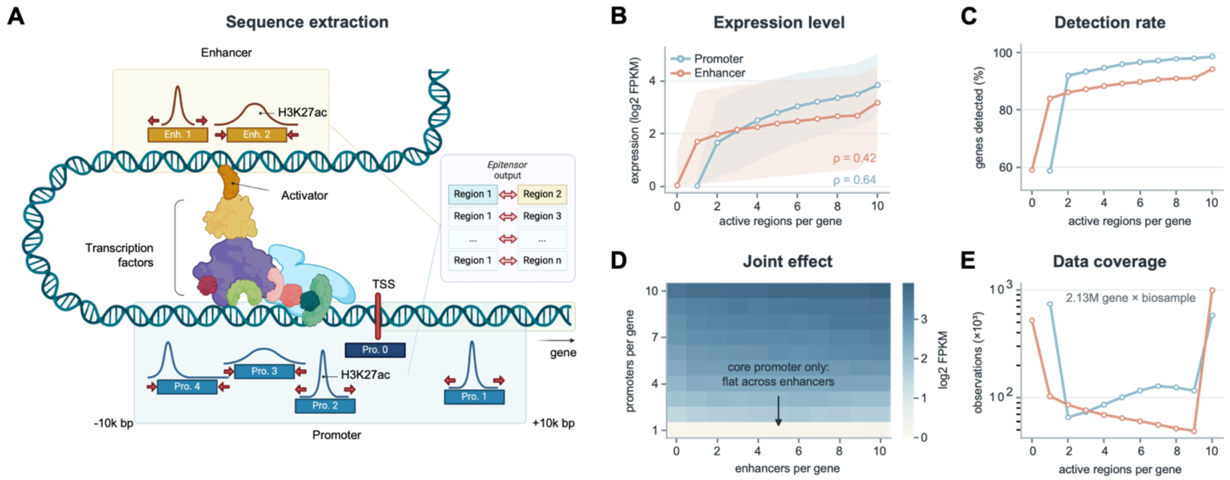


Construction of the model input and its relationship to expression. **(A)** Definition of the up to twenty regulatory sequences supplied to CNN-CR for each gene. Promoter sequences (blue) are drawn from a fixed window of TSS ± 10 kb and comprise a mandatory core promoter centred on the CAGE-annotated TSS (Pro. 0) together with up to nine EpiMap concatenated promoter peaks (Pro. 1 onwards), so every gene receives at least one promoter sequence. Enhancer sequences (yellow) are identified through EpiTensor-predicted three-dimensional contacts (right), which pair the gene's promoter region with distal regions, and are retained where an EpiMap enhancer peak overlaps the contacted region. Red arrows denote standardization of every selected peak to 400 bp by symmetric expansion or contraction around the peak centre. Looping brings the bound enhancer and its TFs into contact with the promoter, which is the configuration EpiTensor infers. **(B)** Median expression against the number of active regions per gene, for promoters (teal) and enhancers (coral), with the interquartile range shaded; Spearman rho is given on the panel. All panels use the same 2,129,296 gene × biosample observations (18,356 genes × 116 biosamples), with expression as log2 FPKM. Promoter counts begin at 1 rather than 0 because of the mandatory core promoter, so promoter 1 and enhancer 0 both denote a gene with no called peak. **(C)** The same relationship split into whether a gene is expressed at all: the percentage of gene × biosample observations with any detected expression (FPKM > 0 in the EpiMap RNA-seq; no other cutoff is applied), plotted against the number of active regions. Promoters rise from 59% to 99% and enhancers from 59% to 94%, so region number governs whether a gene is expressed at all, not only how strongly. **(D)** Median expression for every combination of promoter and enhancer count. Each of the 110 cells contains at least 1,892 observations. Genes carrying only the core promoter (bottom row) stay near zero across the entire enhancer range (median 0.00 to 0.07), whereas genes with ten promoters span 1.06 to 1.18, indicating that enhancer number does not compensate for an absent promoter peak. **(E)** Number of observations contributing to each bin of (B) to (D), on a logarithmic scale. The bins are very unevenly populated: 35% of all gene × biosample observations have only the mandatory core promoter (promoter count 1, no called promoter peak), and 47% sit at the cap of ten enhancers, so the medians in (B) to (D) rest on tens of thousands of observations in the middle bins but on hundreds of thousands at the two ends, and the enhancer axis is truncated by the cap.

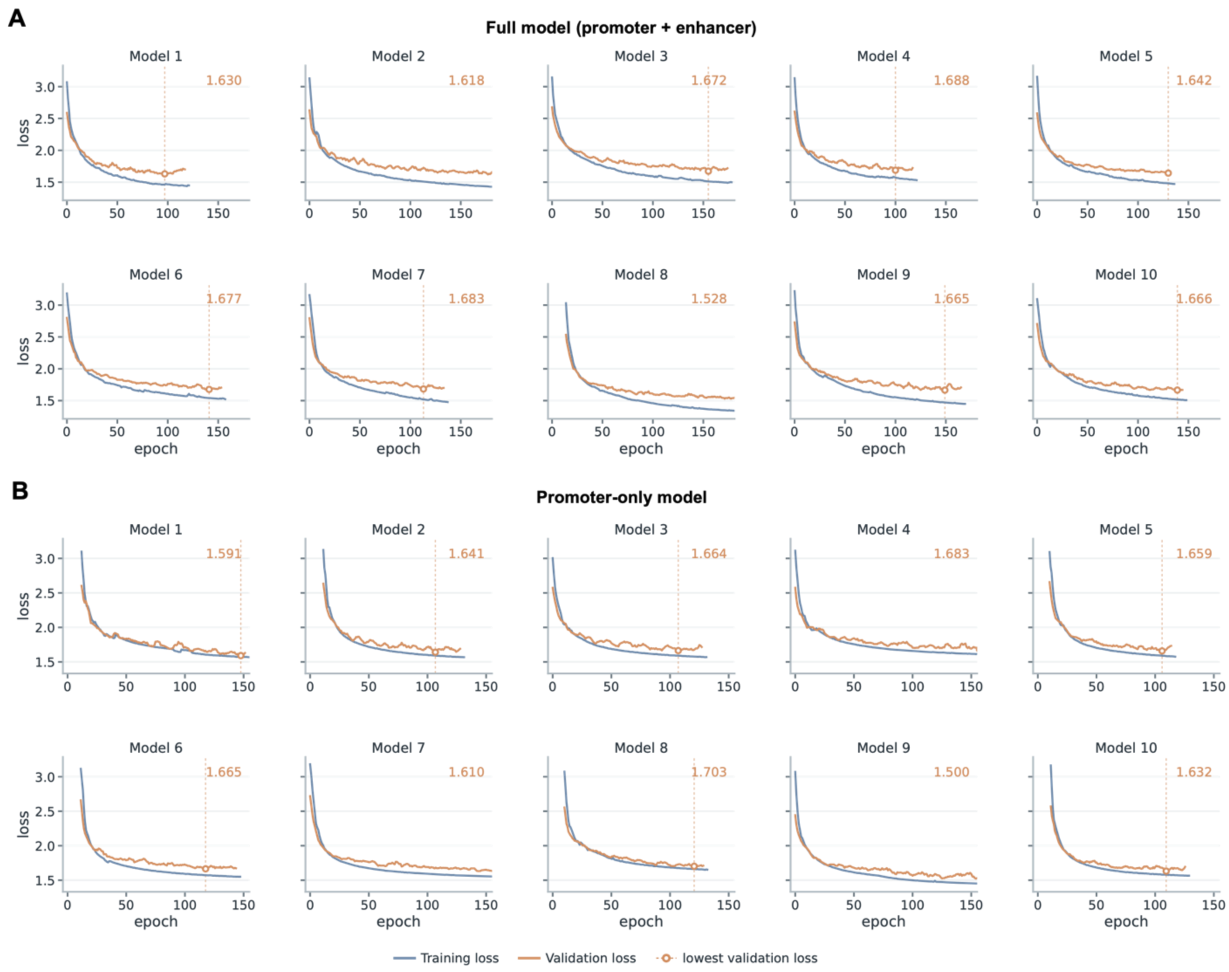


**Fig. S2.**

Training and validation loss for the ten CNN-CR models. **(A)** The full promoter-plus-enhancer model. **(B)** The promoter-only model. Each shows training loss (blue) and validation loss (orange) against epoch for the ten independently initialized CNN-CR models trained on K562, one panel per model. Open circles and dashed vertical lines mark the epoch of lowest validation loss, the point at which training was stopped, and that value is printed in each panel. Loss is mean squared error on log2 FPKM. Validation loss is smoothed with a five-epoch moving average. Within each of (A) and (B) all ten panels share a common axis range, so models can be compared directly. Training ran for 110 to 177 epochs in (A) and 104 to 151 in (B), reaching lowest validation losses of 1.528 to 1.688 and 1.500 to 1.703 respectively.

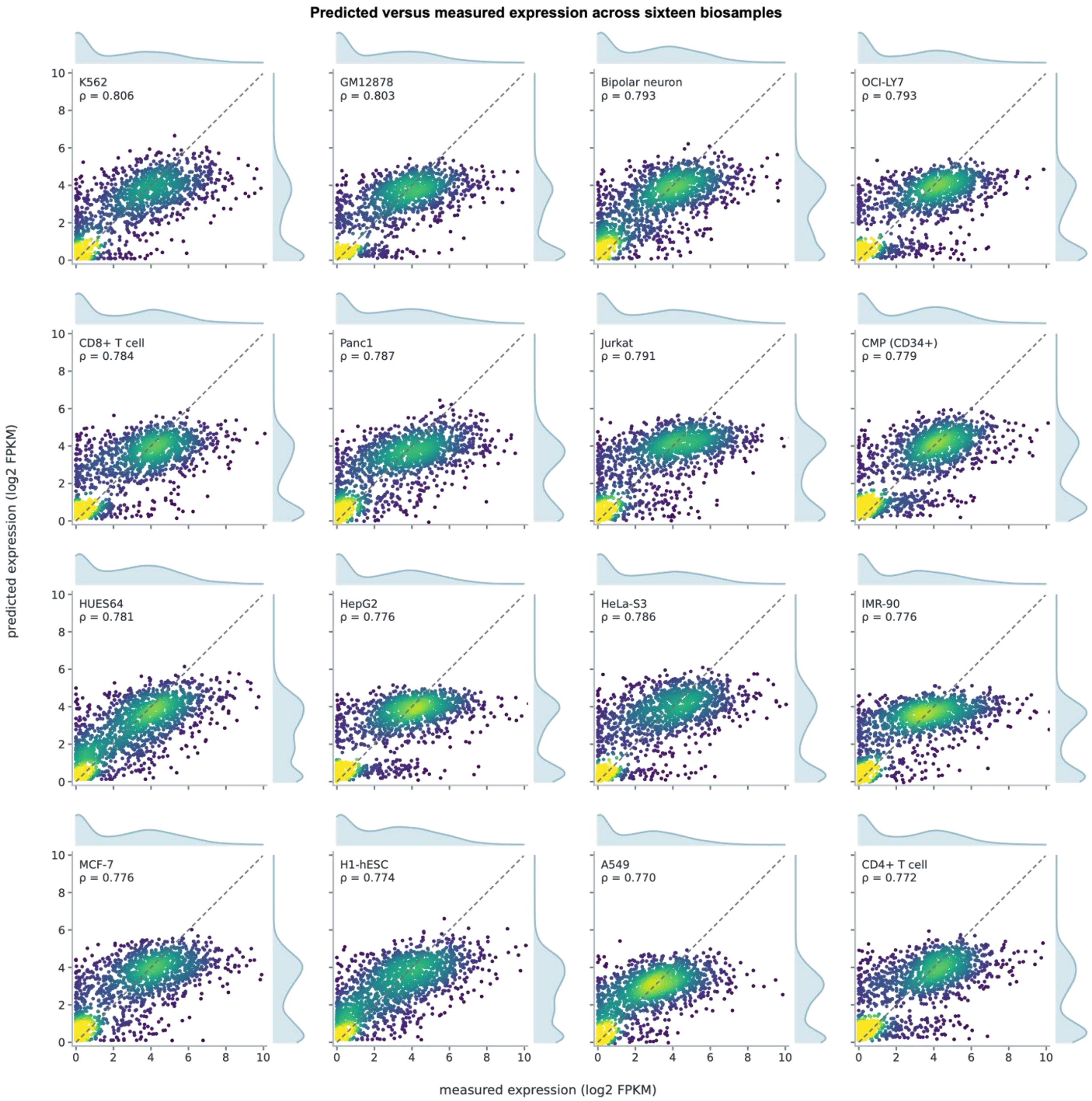


**Fig. S3.**

Predicted versus measured expression across sixteen biosamples. Held-out predicted expression against measured expression (log2 FPKM) for sixteen high-performing biosamples of the 116, ordered by test Spearman rho and chosen to span established cancer lines, primary immune cells, a haematopoietic progenitor, embryonic stem cells, a differentiated neuron and a fibroblast. Each panel shows the 1,836 held-out genes of that biosample, coloured by local point density with the densest points drawn last; the dashed line is the identity. Predictions are the median across the ten models, the same aggregation used in Fig. 1D, so the K562 panel reproduces that figure exactly (rho = 0.806). Spearman rho ranges from 0.770 to 0.806 with a median of 0.782 across the sixteen, against a median of 0.743 across all 116 biosamples.

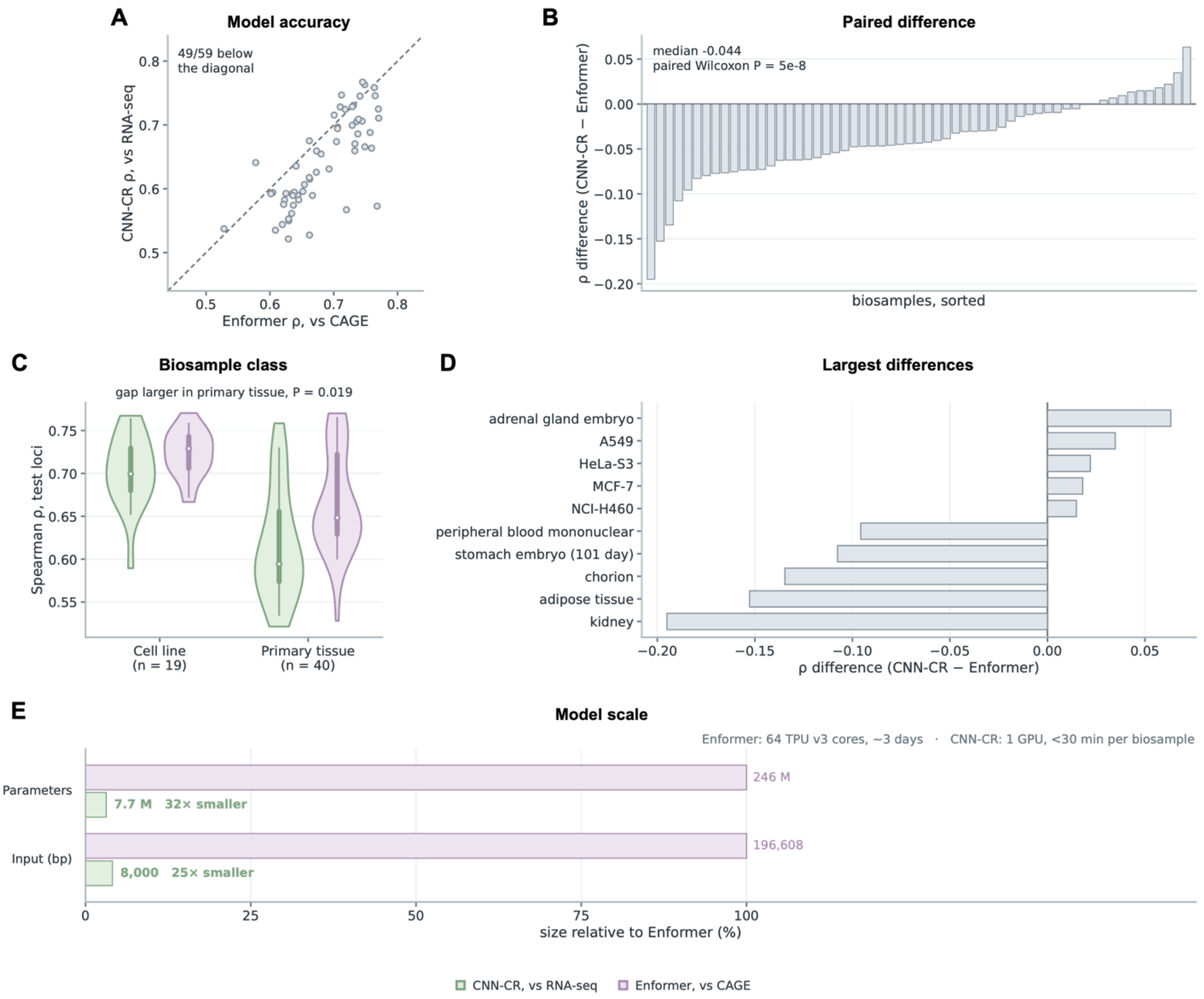


**Fig. S4.**

CNN-CR versus Enformer across 59 matched biosamples. Every panel uses the 1,634 genes that lie inside the genomic intervals Enformer held out for testing; CNN-CR was retrained to exclude these genes. **(A)** Spearman rho for CNN-CR against Enformer, one point per biosample, for the 59 biosamples with Enformer prediction. CNN-CR reaches a median 0.641 against Enformer's 0.680 and sits below the diagonal in 49 of the 59 biosamples. **(B)** The same comparison as a paired difference per biosample, sorted (median -0.044, paired Wilcoxon P = 4.6 x 10-8, CNN-CR higher in 10 of 59). **(C)** The comparison split by biosample class. A biosample was called primary when its EpiMap name carries a donor or developmental descriptor or an explicit tissue term, and an established cell line otherwise. CNN-CR trails Enformer by 0.03 across cell lines (0.699 versus 0.729, n = 19) and by 0.05 across primary tissue (0.594 versus 0.648, n = 40); the gap is significantly larger in primary tissue (Mann-Whitney P = 0.019). **(D)** The five biosamples at each extreme, named. Four of the five in which CNN-CR is ahead are established cancer cell lines, and the five largest shortfalls are all primary tissue. **(E)** Model scale, each quantity expressed as a percentage of Enformer's. CNN-CR has 7,719,700 parameters, being 20 independent branches of 385,984 and 20 branch scalers; the branches do not share weights, so a full model contains 20 x 256 = 5,120 first-layer motif filters and a promoter-only model 10 x 256

= 2,560. Enformer's parameter count is not reported in Avsec et al. and the value shown is computed from its released architecture (1,536 channels, a six-block convolutional tower from 768 to 1,536 channels, 11 transformer layers with 8 heads, key size 64 and value size 192, and the 5,313-track human head). CNN-CR is 32-fold smaller and reads a 25-fold shorter input, 8,000 bp per gene against 196,608 bp. Enformer was trained on 64 TPU v3 cores for approximately 3 days, CNN-CR on one GPU in under 30 minutes per biosample.

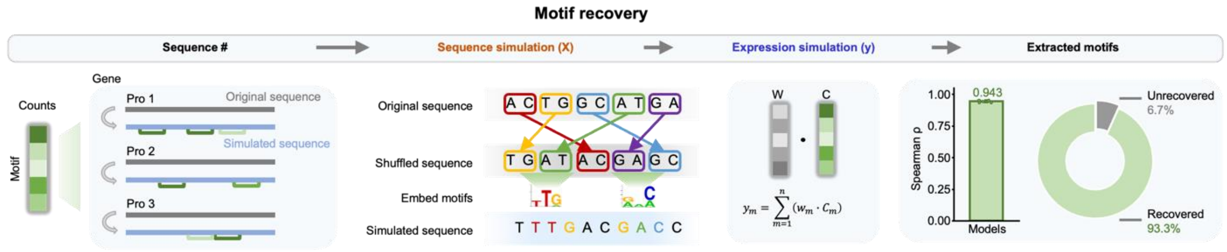


**Fig. S5.**

Recovery of embedded motifs from simulated data. Validation that CNN-CR's first-layer filters learn genuine sequence patterns rather than positional or spurious correlations, shown left to right as the simulation pipeline and its outcome (Supplementary Methods S1). Sequence #: for each of the 18,356 genes, every promoter sequence is replaced by a simulated sequence carrying a defined number of copies of each motif, taken from the per-gene count matrix C at left (green, one row per motif), which records how many times each of the biosample's own non-redundant promoter motifs (736 to 785 per biosample) occurs in the gene's real promoter peaks. Sequence simulation (X): each real promoter is dinucleotide-shuffled, preserving dinucleotide frequency while destroying its original motif content, and the motif copies are then embedded at random non-overlapping positions; across the five biosamples this placed 2.0 to 2.8 million copies into 88,171 to 111,531 promoter sequences (18 to 31copies per 400-bp sequence; median 101 to 130 copies per gene). Expression simulation (y): simulated expression is a linear function of the per-motif count vector with uniform weights plus Gaussian noise, $y_g = \Sigma_m w_m C_{g,m} + \varepsilon_g$, where $C_{g,m}$ is the number of embedded copies of motif m in the promoters of gene g, $w_m = w$ for all m, with w chosen per biosample so that the signal term has the standard deviation of the real log2 FPKM, and $\varepsilon_g \sim N(0, \sigma^2)$ with $\sigma = 0.05$. Extracted motifs: ten promoter-only CNN-CR models per biosample were trained on the simulated data using the settings applied to real data, across five simulated datasets (A549, GM12878, HeLa, HepG2, K562). The models reached a median Spearman correlation of 0.943 against simulated expression (bar; 0.938 to 0.953 across the five biosamples). Motifs were extracted and de-replicated as for real data (2,014 to 2,233 non-redundant motifs per biosample) and compared with the embedded set by TOMTOM; an embedded motif counted as recovered when at least one extracted PWM matched it at $p \leq 1.2 \times 10^{-3}$, the same p-value strength as the $E \leq 1$ cutoff used against JASPAR (Supplementary Methods S1). A median of 93.3% of the embedded motifs per biosample was recovered (donut; 91.8% to 94.7% across biosamples) and the remainder had no significant match. The bar shows the median over the five biosamples with the full range as the error bar and one point per biosample.

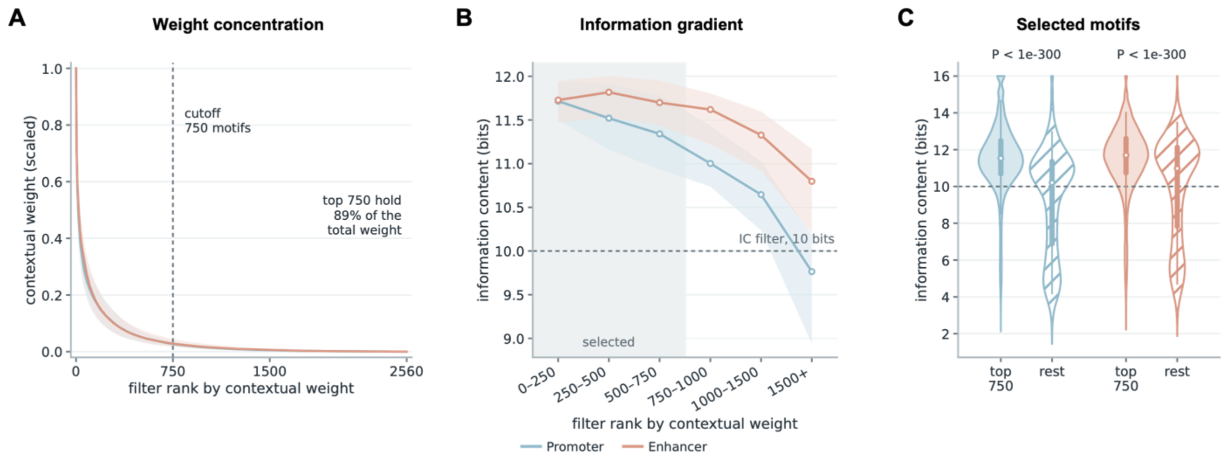


**Fig. S6.**

Contextual weight selects the motif vocabulary. The contextual-weight module supplies the criterion by which each full model's 5,120 first-layer filters (20 branches x 256, or 10 x 256 for the promoter-only model) are reduced to the compact vocabulary used throughout. **(A)** Contextual weight against filter rank, each model min-max scaled and then summarized rank by rank across all 1,160 models (116 biosamples x 10 models; line, median; band, interquartile range). The distribution is steeply concentrated: the 750 highest-weighted filters carry 89% of the total weight in promoters and 88% in enhancers, so the great majority of filters contribute almost nothing. **(B)** Median information content of the same filters, binned by that contextual-weight rank (116 biosamples, model 0; line, median across biosamples; band, interquartile range). Information content falls monotonically with rank, from 11.7 bits in the top 250 to 9.8 bits beyond rank 1,500 for promoters, so the ranking recovers information-dense, motif-like filters without ever being told to. Shaded region, the 750 filters retained. **(C)** Information content of the retained filters against the discarded remainder. Selected filters are markedly more information-dense (promoter median 11.53 versus 10.20 bits, enhancer 11.70 versus 10.98; one-sided Mann-Whitney $P < 1e\text{-}300$ for both), and 87.6% of selected promoter filters exceed the 10-bit threshold applied next, against 53.3% of those discarded. Dashed line, that 10-bit filter. After both steps an average of 657 promoter and 649 enhancer motifs survive per model, which merge across the ten models into the roughly 700 non-redundant motifs per biosample reported in the main text.

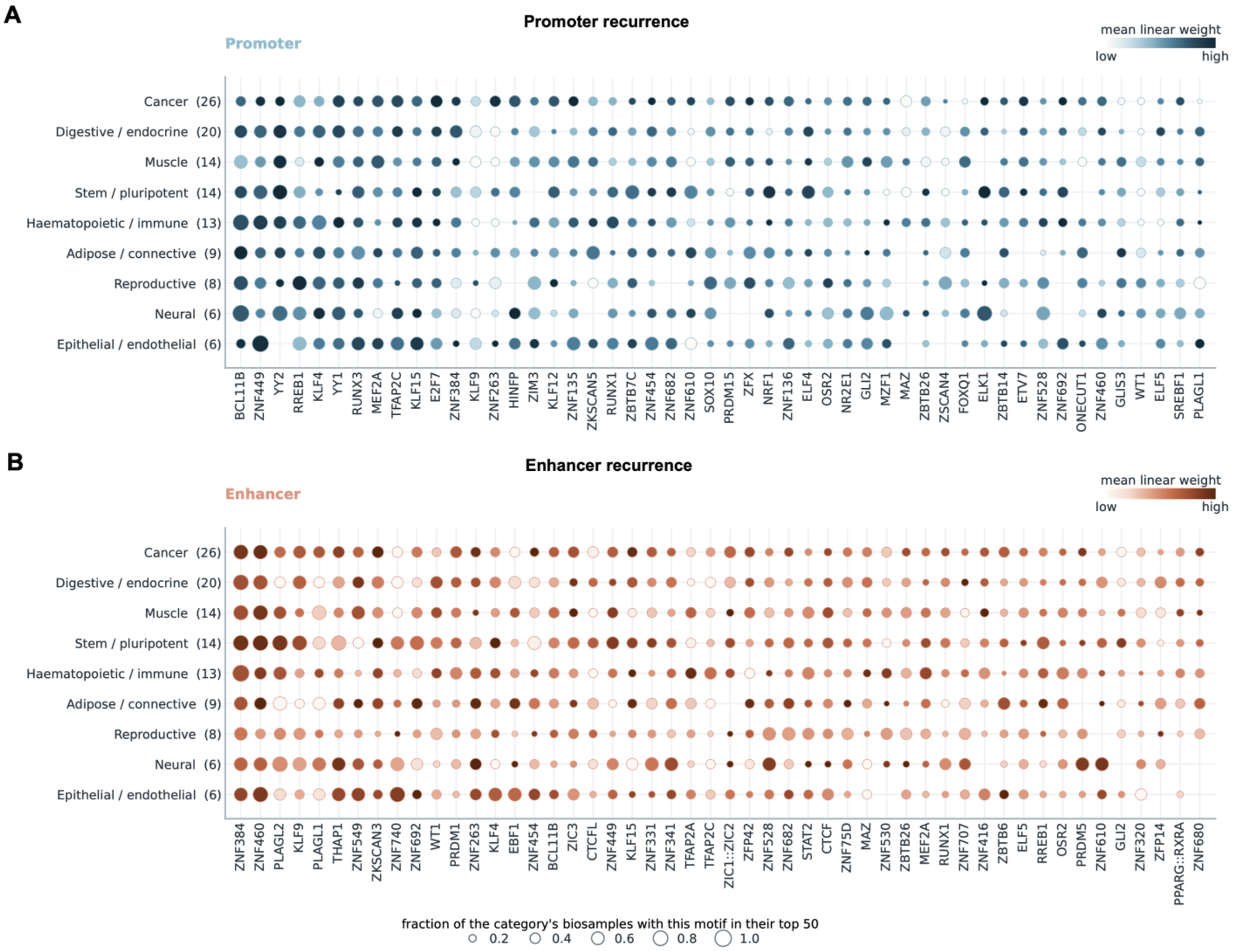


**Fig. S7.**

Motif recurrence across cell-type categories. Recurrence of individual JASPAR-matched motifs across the nine cell-type categories into which the 116 biosamples were grouped, for promoter motifs (a) and enhancer motifs (b). Within each biosample, matched motifs sharing a TF symbol were collapsed by summing their contextual weight, ranked by absolute weight, and the 50 highest retained; the 50 TF symbols appearing in the most top-50 lists are shown. Dot area gives the fraction of that category's biosamples whose top 50 contains the motif, and dot colour the mean linear weight of the motif in that category, square-root scaled. Category sizes are given in parentheses. Motifs are ordered by how many biosamples they recur in, so the leftmost are the most broadly recurrent. Members of the ELF, ELK, KLF, ZNF and YY families are present in every category rather than being confined to the lineage they are usually associated with: in promoters YY2, YY1, KLF4, KLF15, ZNF449 and BCL11B are near-ubiquitous, and in enhancers ZNF384, ZNF460 and PLAGL2 recur across all nine. A minority are category-restricted, for example MEF2A in muscle and EBF1 in haematopoietic and immune samples, but these are the exception.

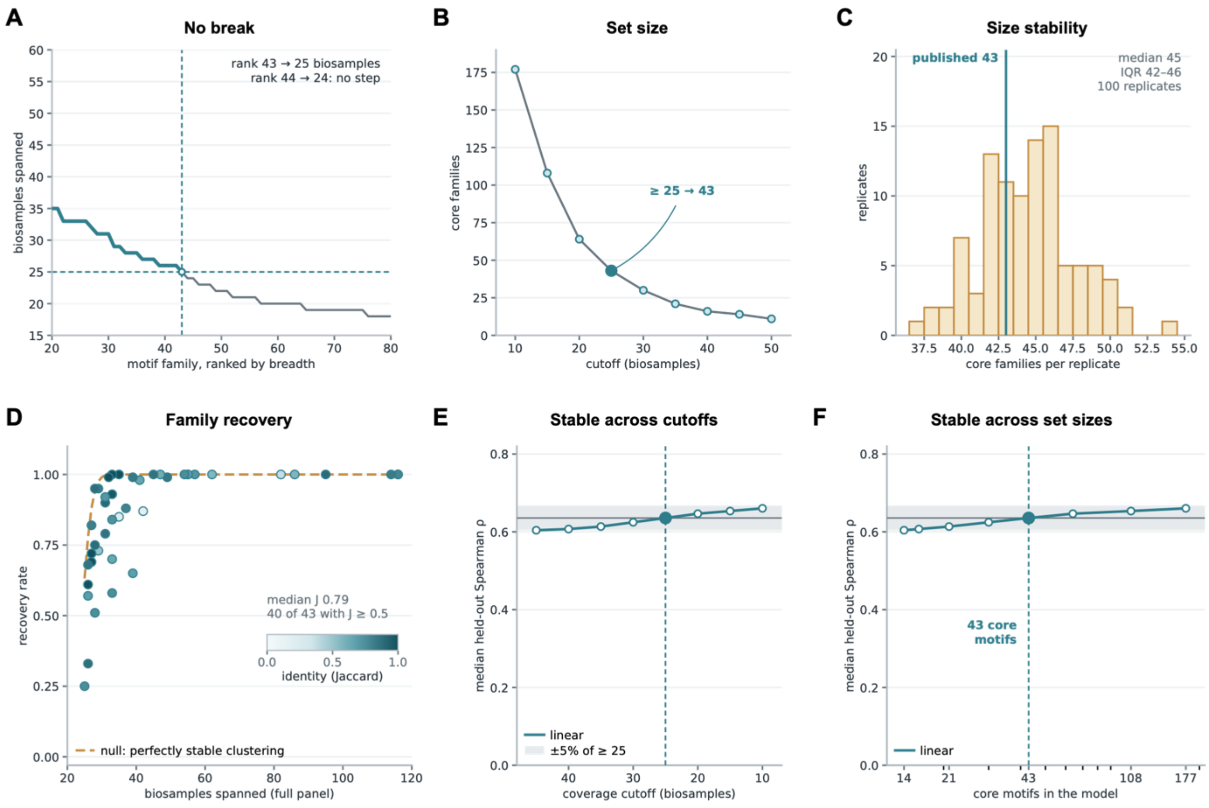


**Fig. S8.**

The core motif set is reproducible even though the cutoff that defines it is a convention. **(A)** Cross-biosample coverage around the cut: the family ranked 43rd spans 25 biosamples and the 44th spans 24. Teal, the 43 retained families; dashed lines, the >= 25 criterion. **(B)** Core families obtained at each of eight cutoffs from >= 10 to >= 45 biosamples (177 to 14 motifs); filled marker, the published 43. **(C)** Core families per replicate when the derivation is repeated on 100 random subsamples of 93 of the 116 biosamples under a proportionally scaled cutoff (Methods): median 45, interquartile range 42–46; teal line, the selected 43. **(D)** Fraction of replicates in which each published family was again called core (recovery), against the number of biosamples it spans; fill, median Jaccard index (shared members divided by the union of members) between the published family and its best-matching replicate family, a measure of how much of the family's membership returns: median 0.79 over the 43 families; 40 of 43 have a median Jaccard of at least 0.5, meaning that at least half of the combined membership is shared, and the three below it are families that are recovered but split in two. In a typical replicate 36 of the 43 families are called core again (range 32– 42; table S1). Dashed line, the expectation for a perfectly stable clustering. **(E and F)** Median held-out Spearman rho across all 116 biosamples of an ordinary least squares model trained on the core motifs surviving each cutoff, under the genes and split of Fig. 2E, read against the coverage cutoff (e) and the number of motifs (f). Horizontal line and shaded band, the >= 25 result (rho = 0.636) and +/- 5% of it; dashed line and filled marker, the published 43. Regardless of the cutoff and the resulting motif-set size (14 to 177 motifs), performance remains within a small range, within 5% of the 43-motif result (rho 0.604 to 0.660).

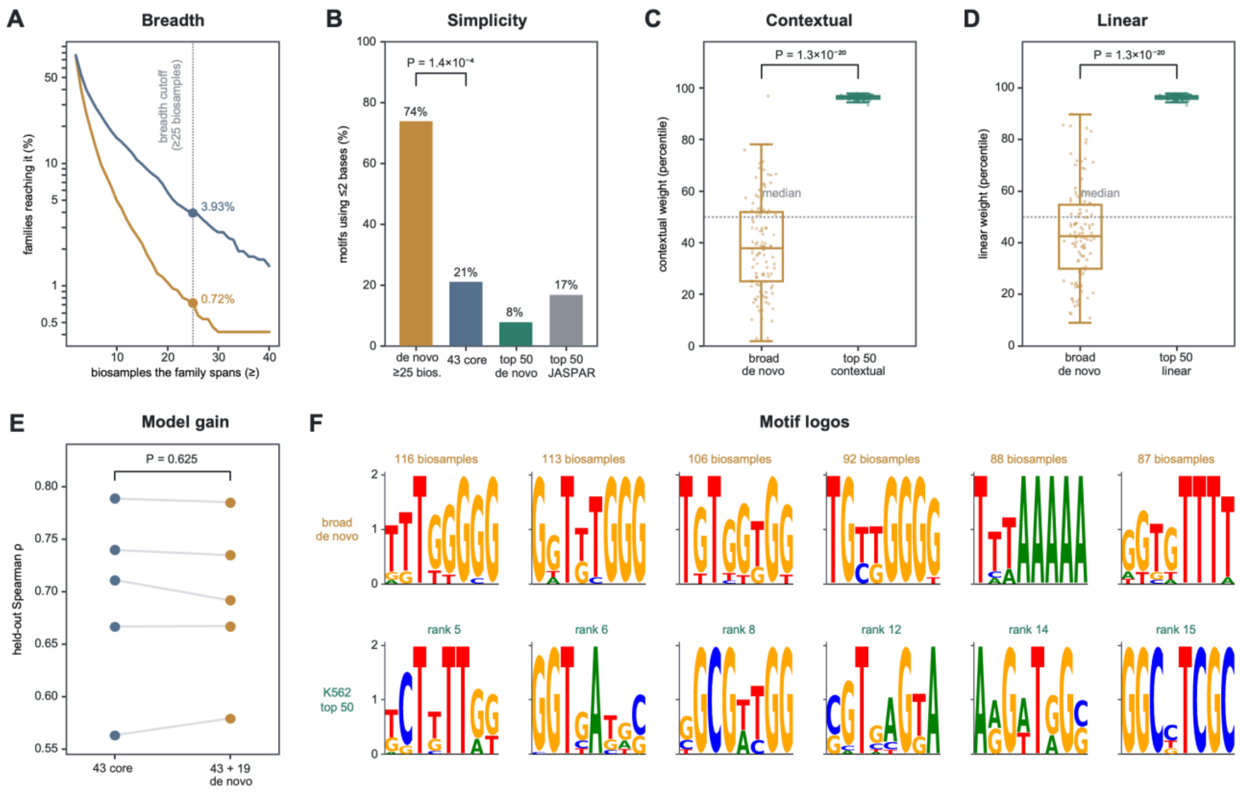


**Fig. S9.**

Uncatalogued de novo motifs are cell-type-restricted, and the broad exceptions are compositionally simple, weakly weighted and add no predictive power. **(A)** Percentage of motif families spanning at least N biosamples, for the 2,623 families formed by uncatalogued (JASPAR-unmatched) de novo motifs carried through the cross-biosample clustering with the expression requirement lifted (orange) and the 1,095 catalogued groups of the published derivation (blue). At the ≥ 25-biosample cutoff (dashed line), 0.72% of de novo families qualify against 3.93% of catalogued groups. **(B)** Percentage of motifs whose consensus uses at most two distinct bases, for the 19 broad de novo families, the 43 core motifs, and the de novo ($n = 26$) and JASPAR-matched ($n = 24$) motifs of K562's promoter top 50; two-sided Fisher's exact test. **(C and D)** Within-biosample percentile rank of absolute weight for the promoter motifs matching one of the 19 broad families (orange; $n = 460$ across 115 biosamples), against the 50 highest-weighted motifs of the same biosample (green); contextual weight and its top 50 in (C), ridge weight and its top 50 in (D), so each panel is drawn on a single weight scale. Points are per-biosample medians; dashed line, the biosample median. The families fall below their biosample's median in 80/115 biosamples for contextual weight ($P = 1.1 \times 10^{-8}$) and 79/115 for ridge weight ($P = 1.2 \times 10^{-4}$), and below its 50 highest-weighted motifs in 114/115 and 115/115 (P values on panel); two-sided Wilcoxon signed-rank throughout. **(E)** Held-out Spearman ρ of the motif-initialized CNN-CR given the 43 core motifs or all 62 (43 core plus the 19), in five biosamples spanning the performance range. The 62-motif model alone was hyperparameter-tuned (Methods), so the comparison favours the larger set; the 19 did not increase performance (mean $\Delta\rho = -0.002$, improved in 2 of 5; paired two-sided Wilcoxon P as shown). **(F)** Sequence logos (information content, bits): the six broadest de novo family representatives with the number of biosamples each spans (orange), and the six highest-ranked de novo motifs of K562's promoter top 50 with their ranks (green).

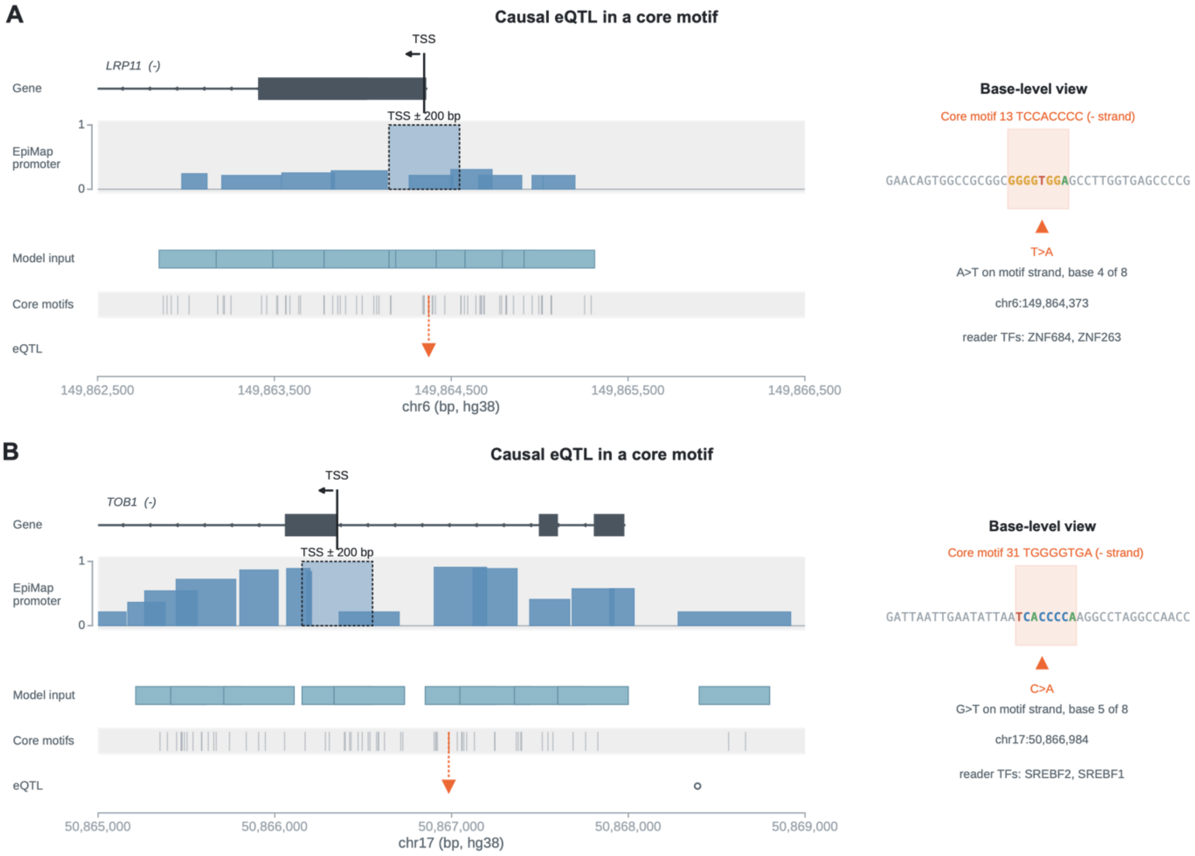


**Fig. S10.**

Two further fine-mapped causal eQTLs inside core-motif occurrences at active promoters. As in Fig. 4C, each locus shows the gene model with its TSS, the matched biosample's Epimap promoter signal, the promoter windows supplied to the model, all core-motif occurrences, and the eQTL positions (orange arrow, the fine-mapped causal variant; open circles, non-causal variants of the same tissue). The base-level view at right gives the 40-bp sequence around the causal variant, the overlapping core-motif occurrence with its consensus and strand, the substitution the variant introduces, and the motif's candidate reader TFs. **(A)** LRP11 in thyroid: causal variant chr6:149,864,373 T>A, 26 bp upstream of the TSS, inside a core-motif occurrence (consensus TCCACCCC) with candidate readers ZNF684 and ZNF263. **(B)** TOB1 in tibial nerve: causal variant chr17:50,866,984 C>A, 631 bp upstream of the TSS, inside a core-motif occurrence (consensus TGGGGTGA) with candidate readers SREBF2 and SREBF1.

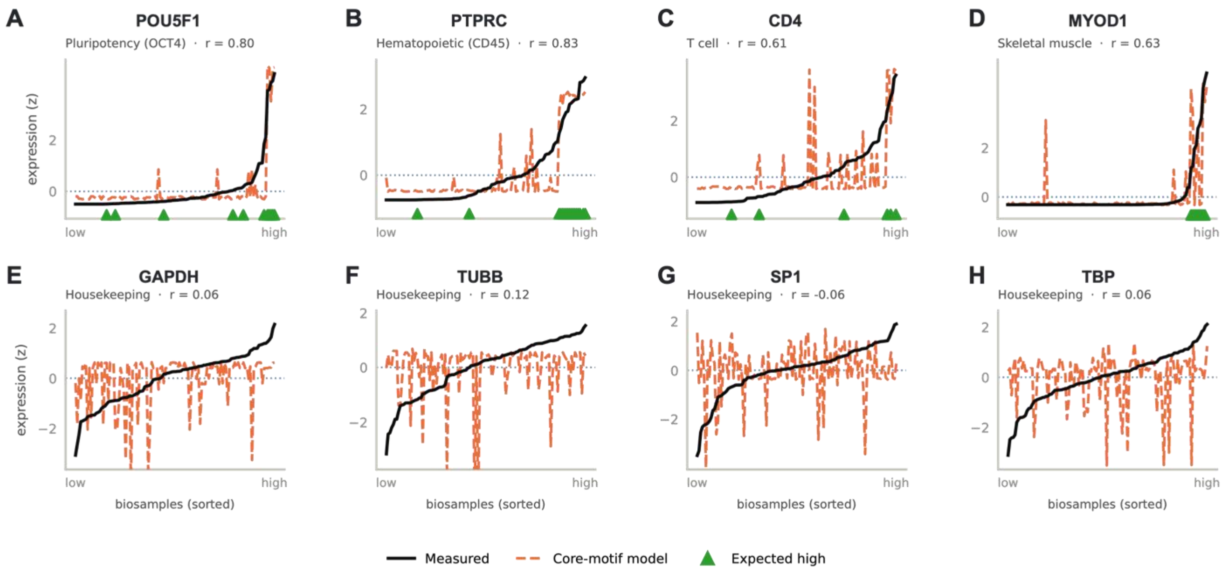


**Fig. S11.**

Cell-type-specific markers are tracked while housekeeping genes are not. As in Fig. 4G: measured (black) and core-motif-predicted (orange dashed) expression across the 116 biosamples, both z-scored per gene and sorted by measured level; green triangles mark biosamples expected to express the marker; the cross-cell-type Pearson r above each panel is computed under the standardization used for Fig. 4F (Methods). **(A to D)** four representative lineage markers: the pluripotency factor POU5F1 (OCT4, r = 0.80), the pan-hematopoietic marker PTPRC (CD45, 0.83), the T-cell marker CD4 (0.61) and the muscle regulator MYOD1 (0.63); the prediction rises in the biosamples expected to express each marker. **(E to H)** four housekeeping controls: GAPDH (0.06), TUBB (0.12), SP1 (−0.06) and TBP (0.06); with no cell-type-specific pattern to follow, the predictions do not track the measured curves.

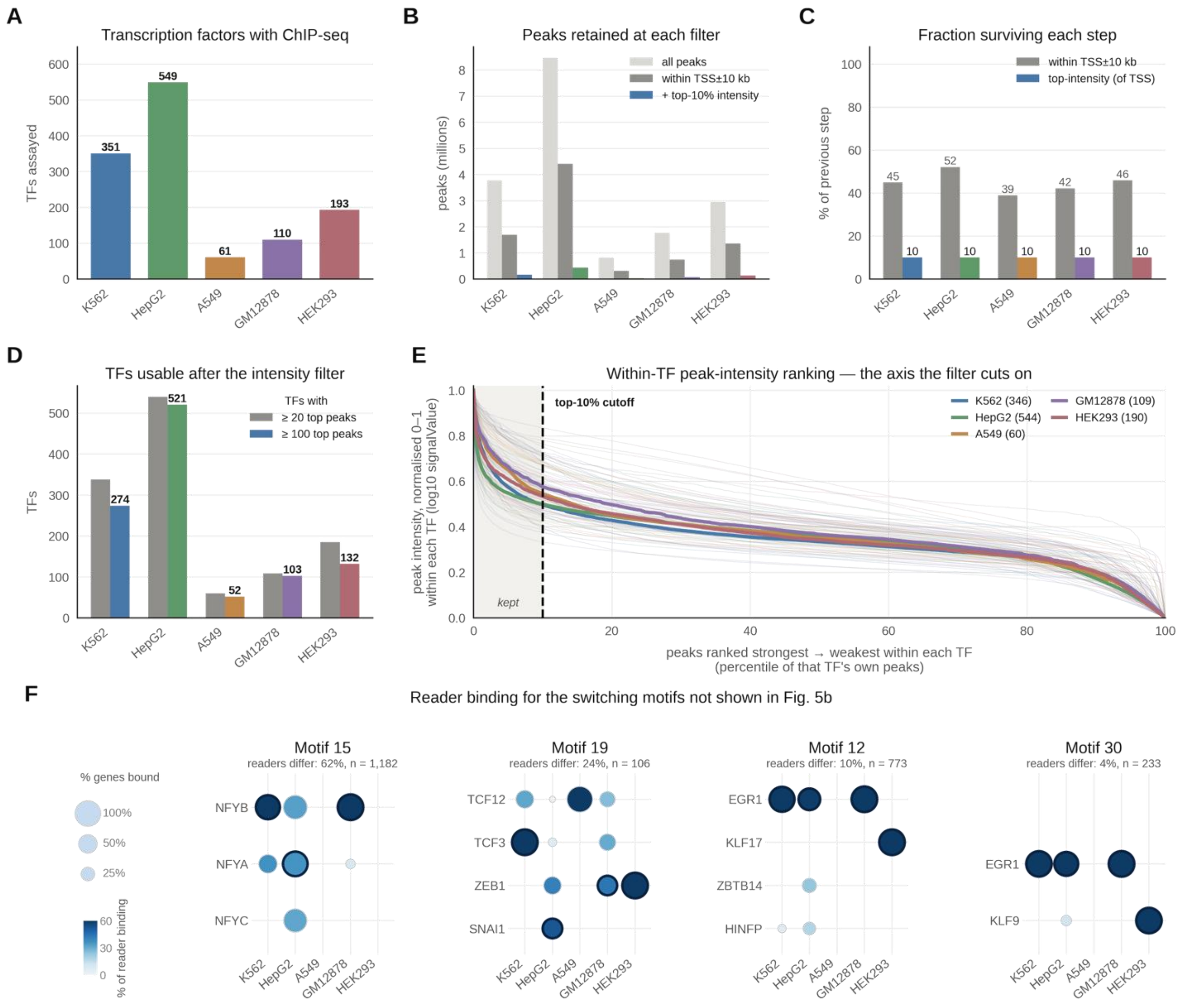


**Fig. S12.**

ChIP-seq peak selection for the site-level reader assignment. How the peaks behind the reader assignment of Fig. 5B,C were chosen (a–e), and the reader spectra of the switching motifs not shown in Fig. 5B (f), for each of the five ENCODE ChIP-profiled cell lines: a factor's peaks are kept when they overlap a TSS ± 10 kb promoter window and their binding signal falls in the strongest 10% of that factor's own peaks (a within-factor percentile, because absolute signalValue varies ~10–15-fold between experiments; Methods). **(A)** Transcription factors with ChIP-seq per line (K562 351, HepG2 549, A549 61, GM12878 110, HEK293 193). **(B)** Peaks retained at each filter: all peaks, peaks within TSS ± 10 kb of a promoter, and the top-10% intensity subset. **(C)** Fraction of peaks surviving each step: 39–52% of each line's peaks are promoter-proximal, and the intensity filter keeps 10% of those by construction. **(D)** Factors usable after the intensity filter, with at least 20 (grey) or at least 100 (colour) top-intensity promoter peaks (≥ 100: K562 274, HepG2 521, A549 52, GM12878 103, HEK293 132). **(E)** Peak intensity (log10 signalValue, normalized 0–1 within each factor) against the percentile rank of the factor's own peaks, strongest first; thin lines, individual factors; thick lines, per-line medians; dashed line and shading, the top-10% cutoff the filter cuts on. **(F)** Reader-binding spectra for the switching motifs not shown in Fig. 5B, encoded exactly as in Fig. 5B: rows, candidate readers; columns, the five cell lines; bubble area, the percentage of the motif's bound genes at which the reader binds; shade, that reader's share of all candidate-reader binding at the motif in that line; dark ring, the dominant reader of that line, the reader bound at the most genes (at least 20). Motifs 15, 19, 12 and 30 complete the eight motifs of Fig. 5C whose dominant reader changes between lines. Above each grid, under the motif number, is the motif's switch rate, defined as in Fig. 5C: over all pairs of the five cell lines, the percentage of genes bound in both lines of a pair at which the readers bound there are not the same, comparing only readers ChIP-sequenced in both lines; n is the number of gene-pair observations scored, a gene bound in several lines contributing once per qualifying pair. Motif 15 (NFYB in K562 and GM12878, NFYA in HepG2; 62%, n = 1,182), motif 19 (TCF3 in K562, SNAI1 in HepG2, ZEB1 in GM12878 and HEK293; 24%, n = 106), motif 12 (EGR1 in K562, HepG2 and GM12878, KLF17 in HEK293; 10%, n = 773) and motif 30 (EGR1 in K562, HepG2 and GM12878, KLF9 in HEK293; 4%, n = 233).

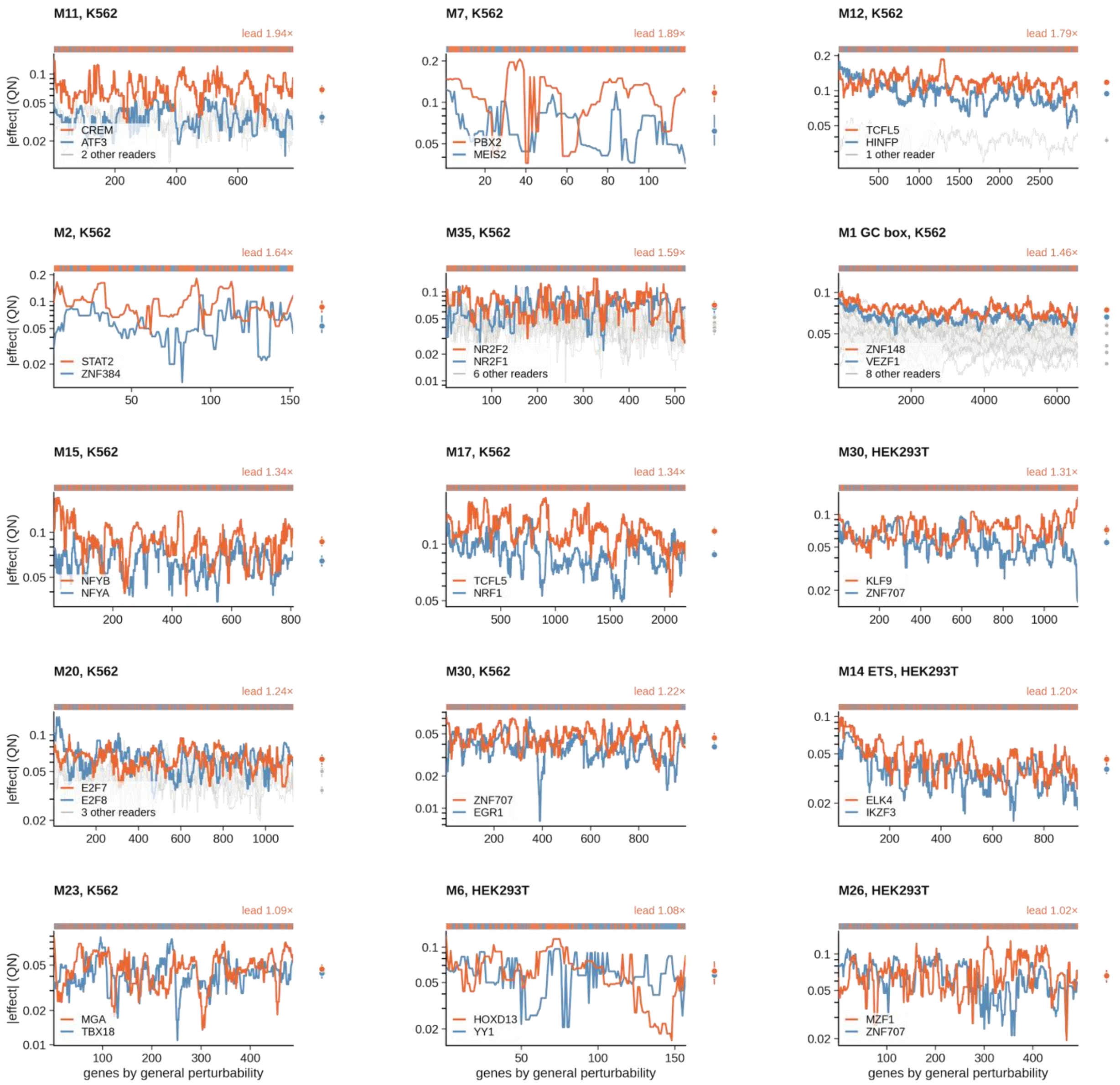


**Fig. S13.**

Reader curves for every motif and cell line panel not shown in Figure 5D. Figure 5E places all nineteen panels on one scale but draws curves for only four of them; the remaining fifteen are drawn here in the same way and in the same order, from the largest lead to the smallest. Each panel is one core motif in one cell line. Every curve is one reader's quantile-normalized absolute knockout effect across that motif's target genes, and a gene enters the set when a copy of the core motif inside one of that reader's ChIP-seq peaks falls in the gene's promoter (Methods). All readers of a panel are drawn on the union of those sets, so they are scored on the same genes. Genes are ordered by general perturbability, how responsive each gene is to perturbation in general (the median absolute knockout effect on that gene across the ChIP-profiled TFs that do not bind it; Methods), so the ordering is fixed before any curve is drawn. Orange, the leading reader; blue, the runner-up; grey, the remaining readers, whose 10th to 90th percentile range is shaded. The strip at the right of each panel gives every reader's median effect over genes, on the same axis, with its 95% bootstrap interval over 400 resamples of genes. The strip above each panel is coloured gene by gene, in the same order as the curves, for whichever of the leading reader and the runner-up has the larger effect at that gene. Lead, the leader over the median of the other readers. The panels run from a clear leader in the cAMP-response motif in K562 (CREM over ATF3, 1.94 fold) down to pairs that are level, where the leader's advantage is 1.02 fold or less and no reader can be called the reader of that motif.

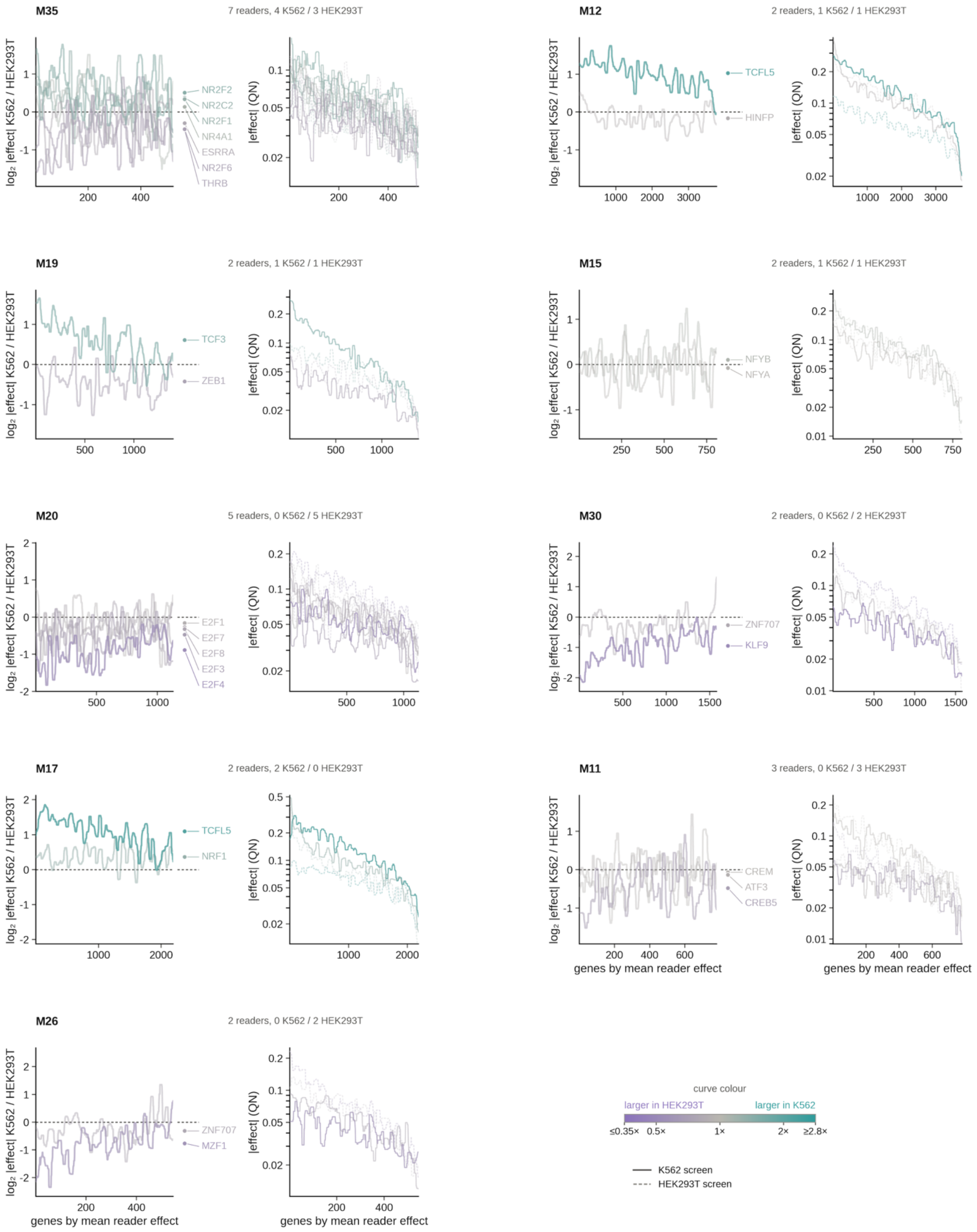


**Fig. S14.**

Cross-cell-type reader preference for every core motif not drawn in Figure 5F. Figure 5F draws every candidate reader of four core motifs; the nine remaining motifs whose readers are perturbed in both screens are given here, in the same way and on the same colour scale. Each motif occupies two panels. Left, every curve is one reader's log2 ratio of its quantile-normalized absolute knockout effect in K562 to its effect in HEK293T, across that motif's target genes; a gene enters the set when a copy of the core motif inside one of that reader's ChIP-seq peaks falls in its promoter, and all readers of a motif are drawn on the union of those sets so that they are scored on the same genes. Genes are ordered by the mean effect of the motif's readers, and curves are smoothed for display with a running median over 2.5% of the genes. Teal, a reader whose effect is larger in K562; violet, larger in HEK293T; the colour saturates at a 2.8-fold median ratio and readers without a cross-line preference are drawn faint, so a motif read by the same factors in both lines appears as a flat grey bundle on the zero line. The marginal at the right of each ratio panel names every reader at its median ratio, with the labels spaced apart where medians coincide. Right of each ratio panel, the two quantities every ratio is formed from: the quantile-normalized absolute knockout effect of each reader on a logarithmic axis, solid for the K562 screen and light dashed for the HEK293T screen, in that reader's ratio colour, so that a ratio driven by one screen alone can be told from one in which both screens move. Above each pair of panels, the number of readers and how many fall on each side; the colour key and the solid/dashed key are at the bottom right, and the colour key is the one used in Figure 5F,G. Motif 20 (five E2F readers, all larger in HEK293T) and motif 17 (NRF1 and TCFL5, both larger in K562) show a whole reader set moving together, which a change of reader would not produce; motif 35 is the clearest split, with four of seven nuclear-receptor readers leaning to K562 and three to HEK293T.

**Table S1. (separate file)**

Robustness of the core-motif set to the recurrence cutoff. **(A)** Cutoff sweep: for each recurrence cutoff from ≥ 10 to ≥ 50 biosamples, the number of motif families, how many of the published 43 they contain (the sets are nested, so every cutoff below 25 contains all 43 and every cutoff above 25 is a subset of them), the median held-out Spearman ρ of the linear and random-forest models across the 116 biosamples on the gene split of Fig. 2E and its ratio to the 43-motif result, and the JASPAR TF entries matched by each set (TOMTOM E ≤ 1, p ≤ 0.01; 287 for the 43 motifs) with the distinct TFs among them (254 for the 43); because the sets are nested, the TFs of every cutoff are a superset or a subset of those of the 43. **(B)** Per-family stability over 100 random subsamples of 93 biosamples with the cutoff scaled to ≥ 20 of 93: for each of the 43 families, its rank and breadth on the full panel, its number of member motifs, the fraction of replicates in which its best-matching replicate family was again called core (under the scaled cutoff and under the literal ≥ 25), the median fraction of its surviving members recovered in that family, the median Jaccard index to it, and its matched JASPAR TFs. Recovery is close to 1 for the 25 broadest families and falls towards the cutoff boundary; 40 of 43 families have a median Jaccard ≥ 0.5, three (universal_4, universal_16, universal_18) are recovered but split into two families. **(C)** Per replicate: the number of families in the subsample derivation, the set size under the scaled and the literal cutoff, how many of the 43 were recovered, and the median Jaccard over the 43.

**Table S2. (separate file)**

GTEx tissue to EpiMap biosample pairing behind Fig. 4A,B. All 26 curated pairs with the number of fine-mapped variants each contributes, whether it cleared the > 500-variant threshold, and the core-versus-control overlap counts; the 10 excluded pairs are included so that the threshold can be seen not to select the direction of the effect.

**Table S3. (separate file)**

The 340 EpiMap biosamples downloaded (EpiMap identifier and name).

**Table S4. (separate file)**

The 116 non-redundant biosamples retained for modelling after removing replicates, sex-specific counterparts and two samples failing quality control.

**Table S5. (separate file)**

Core motif families matching previously reported epigenetic motifs. **(A)** Histone-mark matches: each core motif family with its best match among the published histone-modification motifs, the match statistics and the mark concerned. **(B)** DNA-methylation matches: the same comparison against the published DNA-methylation-associated motifs. Written for this submission; the manuscript reports the counts drawn from these two sheets in the Discussion.

**Table S6. (separate file)**

Audit of the within-biosample merge by shared expressed TF. **(A)** Per biosample: similarity groups formed by the 7,500 promoter filters (TOMTOM E ≤ 0.01, path distance ≤ 0.01), groups retained by the expression filter, groups after the transitive merge by shared expressed JASPAR TF, the number of merged groups and of groups absorbed, and the largest merge. **(B)** Per representative PWM (8,441): the number of similarity groups merged into it, its number of filters and TF names, and the TF names shared between the merged groups. **(C)** Per core family: the number of member representatives, how many of them are merges, the mean number of similarity groups per member, and the TF names that most often drove the merges.

**Data S1. (separate file)**

The 43 core regulator motifs as position weight matrices in MEME format.

**Data S2. (separate file)**

Logo atlas of the core motifs: one page per core motif showing the core position weight matrix and every JASPAR 2022 motif it matches by TOMTOM, drawn in the orientation of the core motif.
